# Metagenomics survey unravels diversities of biogas’ microbiomes with potential to enhance its’ productivity in Kenya

**DOI:** 10.1101/2020.04.23.048504

**Authors:** S.M. Muturi, L.W. Muthui, P.M. Njogu, J.M. Onguso, F.N. Wachira, S.O. Opiyo, R. Pelle

## Abstract

The obstacle to optimal utilization of biogas technology is poor understanding of biogas’ microbiome diversities over a wide geographical coverage. We performed random shotgun sequencing on twelve environmental samples. A randomized complete block design was utilized to assign the twelve biogas reactor treatments to four blocks, within eastern and central regions of Kenya. We obtained 42 million paired-end reads that were annotated against sixteen reference databases using two ENVO ontologies, prior to β-diversities studies. We identified 37 phyla, 65 classes and 132 orders of micro-organisms. *Bacteria* dominated the microbiome and comprised of 28 phyla, 42 classes and 92 orders, conveying substrate’s versatility in the treatments. Though, *fungi* and *Archaea* comprised of only 5 phyla, the *fungi* were richer; suggesting the importance of hydrolysis and fermentation in biogas production systems. High β-diversity within the taxa was largely linked to communities’ metabolic capabilities. *Clostridiales* and *Bacteroidales*, the most prevalent guilds, metabolize organic macromolecules. The identified affiliates of *Cytophagales*, *Alteromonadales*, *Flavobacteriales, Fusobacteriales*, *Deferribacterales*, *Elusimicrobiales*, *Chlamydiales*, *Synergistales* to mention but few, also catabolize macromolecules into smaller substrates to conserve energy. Furthermore, *δ-Proteobacteria*, *Gloeobacteria* and *Clostridia* affiliates syntrophically regulate *P*_*H2*_ and reduce metal to provide reducing equivalents. *Methanomicrobiales* and other *Methanomicrobia* species were the most prevalence *Archaea*, converting formate, CO_2(g)_, acetate and methylated substrates into CH_4(g)_. *Thermococci*, *Thermoplasmata* and *Thermoprotei* were among the sulfur and other metal reducing *Archaea* that contributed to redox balancing and other metabolism within treatments. Eukaryotes, mainly fungi were the least abundant guild, comprised largely *Ascomycota* and *Basidiomycota* species. *Chytridiomycetes*, *Blastocladiomycetes* and *Mortierellomycetes* were among the rare species, suggesting their metabolic and substrates limitations. Generally, we observed that environmental and treatment perturbations influenced communities’ abundance, β-diversity and reactor performance largely through stochastic effect. The study of the diversity of the biogas’ microbiomes over wide environmental variables and the productivity of biogas reactor systems has provided insights into better management strategies that may ameliorate biochemical limitations to effective biogas production.

**Author Summary:** The failure of biochemical reactions in biogas producing systems is a common problem and results from poor functioning of the inhabiting micro-organisms. A poor understanding of the global diversities of these micro-organisms and lack of information on the link between environmental variables, biogas production, and community composition, contrains the development of strategies that can ameliorate these biochemical issues. We have integrated sequencing-by-synthesis technology and intensive computational approaches to reveal metacommunities in the studied reactor treatments. The identified communities were compared with the treatment’s phenotypic and environmental data in an attempt to fill the existing knowledge gaps on biogas microbiomes and their production capacities. We present 132 biogas taxonomic profiles systematically and comparatively, linking the abundance with the identified environmental variables. The local composition of microbiome and variations in abundance were also linked to the observed differences in biogas productivity, suggesting the possible cause of the observed variations. The detailed information presented in this study can aid in the genetic manipulation or formulation of optimal microbial ratios to improve their effectiveness in biogas production.

## Introduction

The optimal utilization of micro-organisms is a critical strategy in the development of new products and improvement of the existing bioprocesses [1]. One of the widely applied technologies that utilises micro-organisms is anaerobic digestion (AD) of waste materials. The process generates biogas and digestate from wastes, the former being an energy carrier molecule [2]. Over 30,000 industrial installations globally, have been revealed to utilize AD to convert organic wastes into power, producing up to10,000 MW [3]. Apart from power production, stable AD processes provide one of the most efficient environmental models for organic waste valorization that enables nutrient recovery from the digestate [4], and hence providing additional economic benefits [5], when compared to other methods of waste management [6]. Many African countries, including Kenya produce huge amounts of waste [5] with high biogas generating potential. In spite of the efforts made to exploit this resource through the Biogas Partnership Programme [7], the biogas systems in African countries remain fairly rudimentary. The Programmes have installed approximately 60,000 biogas plants in Africa which include 16,419 in Kenya, 13,584 in Ethiopia, and 13,037; 7518, and 6504 in Tanzania, Burkina Faso and Uganda, repectively [5, 7]. Though Kenya leads in biogas exploitation in Africa, adoption of the technology remains poor, and is not commensurate with the country’s energy needs. However, because of the relative popularity of the technology in the country, compared to its’ counterparts, Kenya was justified as the appropriate pilot site for biogas studies among other African countries. Similar guidelines were also followed to select the Kenyan regions in this study.

Overally, it is thought that the poor adoption of the biogas technology in Africa is partly due to poor knowledge on the microbiomes that mediate the AD process. These processes comprise thermodynamic unfavorable reactions that contribute to stability and performance of the treatments. The microbial populations and the reactions involved are highy sensitive to environmental and reactor operation conditions; traits that prompt the need for development of sustainable management strategies for effective biogas production. Without such strategies, a complete breakdown of the whole system occurs. In the event of complete failure, farmers are forced to abandon the systems or invest in costly restoration processes, all of which end up reducing the economic and environmental benefits of the AD systems; thereby constraining efforts towards greening the economy.

Although, previous reports from other continents have indicated dominance of *Bacterial* and *Archaeal* species in tolerating reactor perturbations, limitations of traditional culture-dependent and independent methods e.g 16S rRNA gene [8–9], still limit the knowledge on global biogas microbiomes’ diversities within these systems. The 16S rRNA techniques used thus far provide low resolution [10, 11, 12], and are not robust enough to unravel the high level of microbial diversity in biogas ecosystems. All these limit the information available on microbial composition and diversity which would be critical for improvement and better management of biogas systems. The 16S rRNA gene does not reveal the full extent of microbial diversity in an ecosystem [13–17] and most current diversity estimates are extrapolations of empirical scaling laws [13], and other theoretical models of biodiversity [14]. The above limitation constrains the capacity to scrutinize and apply the biodiversity scaling laws, macroecological theories and other ecological theories [12–14] in the context of biogas production. This ends up directly limiting the available microbial diversity knowledge which would be useful to develop viable mitigition strategies to address biochemical reaction failures. Despite covering a small fraction of microbial diversities in a dataset, marker gene based studies, annotate the detected species against biological databases that comprise sequences of interest. This approach has the potential to lead to conflicts on the links between microbial populations and reactor performance. Such conflicts were reported by Wittebolle *et al*. [18] and Konopka *et al* [19]. Wittebolle *et al*. considered higher microbial diversity as a reservoir of redundant metabolic pathways that possess desirable traits that respond to reactor pertubations [18] and other environmental conditions. In contrast, Konopka *et al*. argued that less diverse communities confer system stability by expressing complementary pathways [19] that avoid direct competition over the available resources. These kinds of contrasting views prompted us to ask whether microbial diversity in biogas reactors matters or it is the existence of the core microbiomes that determine the AD stability and maximum biogas production. In an effort to seek answers to these questions, we were provoked to further characterize the biogas system’s micro-organisms using next generation sequencing platforms also termed “massively parallel sequencing” over a wide geographic area. Specifically, the technology employed sequencing-by-synthesis chemistry to generate the dataset. Previously, this technology has enabled massive sequencing of environmental genomes with high quality and accurancy [20]. However, the application of the technology in the biogas systems is still at its infancy. Furthermore, eukaryotes, mainly *fungi* have normally been neglected by the majority of the studies, yet these species, are also crucial in the metabolism of large macromolecules in the AD systems. Our study was therefore designed to address the limitations enumerated above. We expect our findings to contribute to the formation of testable hypotheses for future application of metacommunity theory on the links between microbal diversity and environmental conditions. Further the study will contribute to a greater understanding of biogas microbiomes for improvement of biogas production.

## Results

### Annotations against the 16 biological databases

We used Illumina based shotgun metagenomics to identify and compare microbial communities within and among the biogas reactor treatments. In total 12 sludge samples assigned to twelve different treatments operating at a steady-state were assayed. The procedure yielded a total of 44 million raw reads and out of these, 2 million reads were of poor quality. We revealed 24,189,921 reads of 35-151bps from the remaining 42 million reads, which were filtered based on the % G:C content bias and duplicates according to Gomez-Alvarez *et al*. [21]. Approximately 23,373,513 reads conveying 1,261833 to 3,986675 reads per the treatment were realized (SFig. 1a). Thereafter, we trimmed and assembled the contigs using *de novo* approach and the obtained high quality scaffolds (3,819,927 scaffolds, 1,119,465,586 bps, with mean average length of 291bps, and mean average standard deviation of 410 bps, STable 1) were annotated against sixteen databases using two ENVO ontologies. Taxonomic assignments conveyed by SEED database were the highest (180, 000 scaffolds), while the assignment of NOG, LSU, RDP and COG databases were the least (<8000 scaffolds). Among the databases, IMG, Genbank and TrEMBL conveyed equal scaffold annotations (144,000 scaffolds). Other databases annotated the reads as indicated in Fig. 1. Between the two ontologies, the large lake biome revealed the highest annotation compared to the terrestrial biome with a precision of *P ≤ 0.05* across the treatments. Furthermore, 39.75% to 46.70% and 53.07% to 60.11% genes were conveyed to known and unknown proteins.

**Fig. 1:**
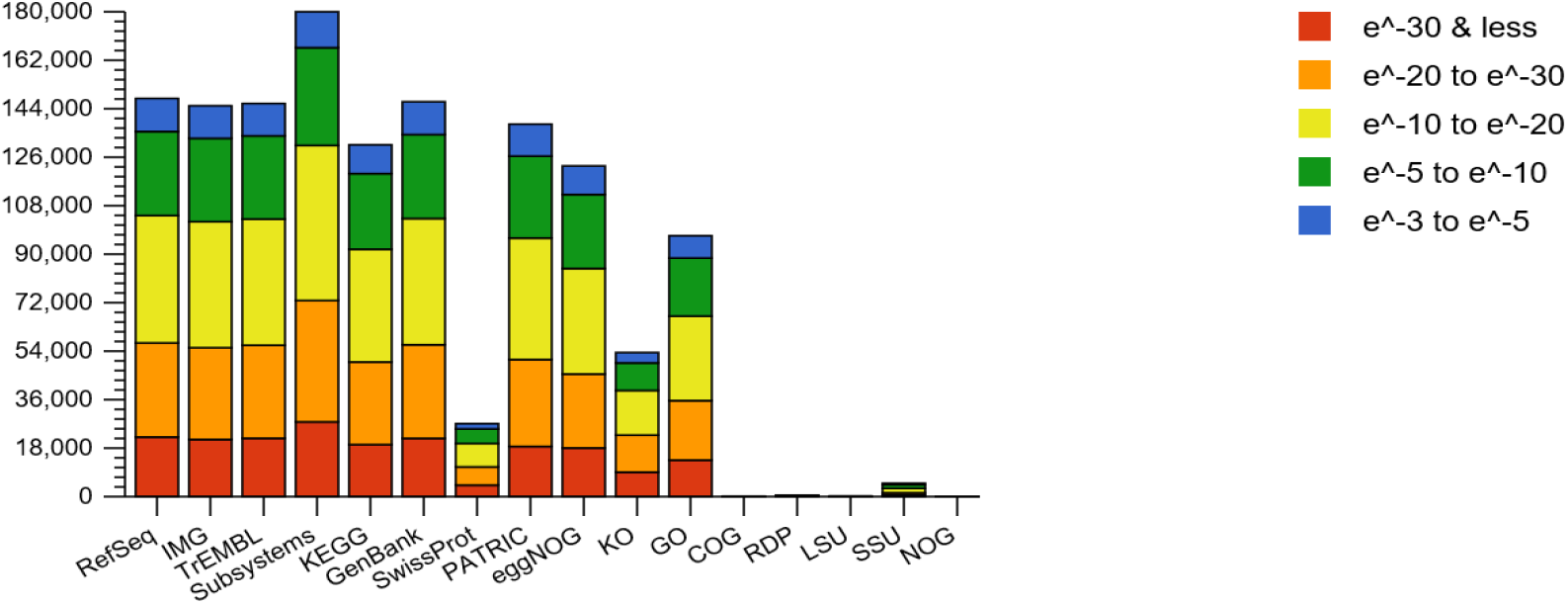
Stacked bar charts showing the source of hits distributions from the sixteen databases: These include nucleotide and protein databases, protein databases with functional hierarchy information, and ribosomal RNA databases. The bars representing annotated reads are colored by e-value range. Different databases have different numbers of hits but can also have different types of annotation data.

**Table 1:** The Krona Software generated *Bacterial* phyla biomes of the twelve treatments within eastern and central regions of Kenya.

| **Phylum** | **S_11.10** | **S_9.10** | **S_12.10** | **S_6.10** | **S_8.10** | **S_3.10** | **S_1** | **S_4.10** | **S_2.0** | **S_5.10** | **S_7.10** | **S_10.10** |
| --- | --- | --- | --- | --- | --- | --- | --- | --- | --- | --- | --- | --- |
| *Firmicutes* | 27% | 34% | 26% | 21% | 25% | 22% | 22% | 31% | 15% | 30% | 29% | 15% |
| *Proteobacteria* | 26% | 19% | 20% | 27% | 24% | 27% | 25% | 19% | 31% | 25% | 21% | 32% |
| *Bacteroidetes* | 15% | 18% | 22% | 11% | 18% | 11% | 15% | 20% | 10% | 16% | 18% | 10% |
| *Actinobacteria* | 4% | 3% | 5% | 5% | 4% | 4% | 4% | 4% | 7% | 5% | 7% | 6% |
| *Chloroflexi* | 3% | 2% | 3% | 6% | 4% | 6% | 4% | 3% | 7% | 2% | 3% | 7% |
| *Spirochaetes* | 2% | 3% | 1% | 3% | 2% | 3% | 4% | 2% | 2% | 1% | 1% | 1% |
| *Verrucomicrobia* | 2% | 2% | 1% | 2% | 1% | 2% | 2% | 1% | 2% | 0.9% | 1% | 2% |
| *Cyanobacteria* | 2% | 1% | 2% | 2% | 2% | 2% | 2% | 2% | 3% | 1% | 2% | 3% |
| *Planctomycetes* | 2% | 1% | 2% | 3% | 2% | 3% | 3% | 2% | 3% | 2% | 2% | 3% |
| *Chlorobi* | 1% | 1% | 1% | 1% | 2% | 1% | 1% | 1% | 1% | 0.9% | 1% | 1% |
| *Acidobacteria* | 1% | 1% | 1% | 2% | 1% | 2% | 1% | 0.9% | 2% | 0.8% | 1% | 2% |
| *Thermotogae* | 1% | 0.9% | 1% | 1% | 1% | 1% | 1% | 0.9% | 1% | 0.8% | 1% | 1% |
| *unclassified reads* | 0.6% | 0.6% | 1% | 1% | 2% | 1% | 0.9% | 1% | 0.6% | 1% | 1% | 0.7% |
| *Synergistetes* | 0.8% | 0.6% | 1% | 0.7% | 0.7% | 0.6% | 0.6% | 0.8% | 0.8% | 0.6% | 0.8% | 0.9% |
| *Lentisphaerae* | 0.8% | 0.5% | 0.3% | 0.4% | 0.3% | 0.4% | 0.4% | 0.3% | 0.3% | 0.2% | 0.4% | 0.2% |
| *Fusobacteria* | 0.6% | 0.8% | 0.6% | 0.5% | 0.6% | 0.5% | 0.5% | 0.7% | 0.4% | 0.7% | 0.7% | 0.4% |
| *D. Thermus* | 0.5% | 0.4% | 0.6% | 0.9% | 0.6% | 0.8% | 0.7% | 0.6% | 1% | 0.5% | 0.6% | 1% |
| *Aquificae* | 0.5% | 0.4% | 0.4% | 0.5% | 0.5% | 0.5% | 0.5% | 0.4% | 0.4% | 0.3% | 0.4% | 0.5% |
| *Tenericutes* | 0.3% | 0.6% | 0.3% | 0.2% | 0.4% | 0.3% | 0.2% | 0.3% | 0.2% | 1% | 0.5% | 0.3% |
| *Deferribacteres* | 0.3% | 0.3% | 0.3% | 0.4% | 0.4% | 0.4% | 0.3% | 0.3% | 0.3% | 0.2% | 0.3% | 0.3% |
| *Nitrospirae* | 0.3% | 0.2% | 0.3% | 0.4% | 0.4% | 0.4% | 0.4% | 0.3% | 0.4% | 0.2% | 0.3% | 0.4% |
| *Chlamydiae* | 0.3% | 0.2% | 0.1% | 0.2% | 0.2% | 0.1% | 0.2% | 0.1% | 0.2% | 0.08% | 0.2% | 0.1% |
| *Fibrobacteres* | 0.2% | 0.5% | 0.1% | 0.2% | 0.1% | 0.2% | 0.2% | 0.2% | 0.08% | 0.2% | 0.2% | 0.08% |
| *Dictyoglomi* | 0.2% | 0.2% | 0.2% | 0.4% | 0.3% | 0.4% | 0.3% | 0.3% | 0.3% | 0.2% | 0.2% | 0.3% |
| *Elusimicrobia* | 0.2% | 0.2% | 0.1% | 0.1% | 0.1% | 0.2% | 0.1% | 0.1% | 0.1% | 0.1% | 0.2% | 0.1% |
| *Chrysiogenetes* | 0.08% | 0.06% | 0.07% | 0.07% | 0.07% | 0.09% | 0.06% | 0.07% | 0.09% | 0.05% | 0.07% | 0.09% |
| *Gemmatimonadete* | 0.06% | 0.06% | 0.07% | 0.1% | 0.1% | 0.1% | 0.1% | 0.07% | 0.2% | 0.6% | 0.08% | 0.2% |
| *C. Poribacteria* | 0.03% | 0.03% | 0.03% | 0.05% | 0.03% | 0.05% | 0.03% | 0.03% | 0.05% | 0.02% | 0.03% | 0.04% |

### Annotations against the archived taxonomic profiles

Taxonomic assignments revealed four broad but relatively simple distributions of microorganisms, with the species of three domains being responsible for methanogenesis. In summary, 2, 617,791 reads hits were generated, with 213681 hits being assigned to *Archaea*, 2379293 hits to *Bacteria* and 21886 hits to *Eukaryotes* (SFig. 1a). Thus, the revealed abundance hits ratio of *Bacteria* to *Archaea* to *Eukaryotes* in the reactor treatments was 109:9:1. The higher abundance of *Bacterial* species was linked to their metabolic capabilities. Unlike other cellular organisms, *Eukaryotes* mainly *fungi* had the least abundances (< 1% of the total hits) among the treatments. Despite these variations of species abundances, we observed consistent microbial richness among the studied treatments. In total 37 phyla, 73 classes and 132 orders were indentified. A majority of the reactor treatments exhibited high community diversity, except for the communities detected in reactor 1, 3 and 6 that were found to cluster partially on the upper right quadrant of the plot (Fig. 2), at a *P ≤ 0.05*. All the identified phylotypes were further annotated and β-diversity determined in details.

**Fig. 2:**
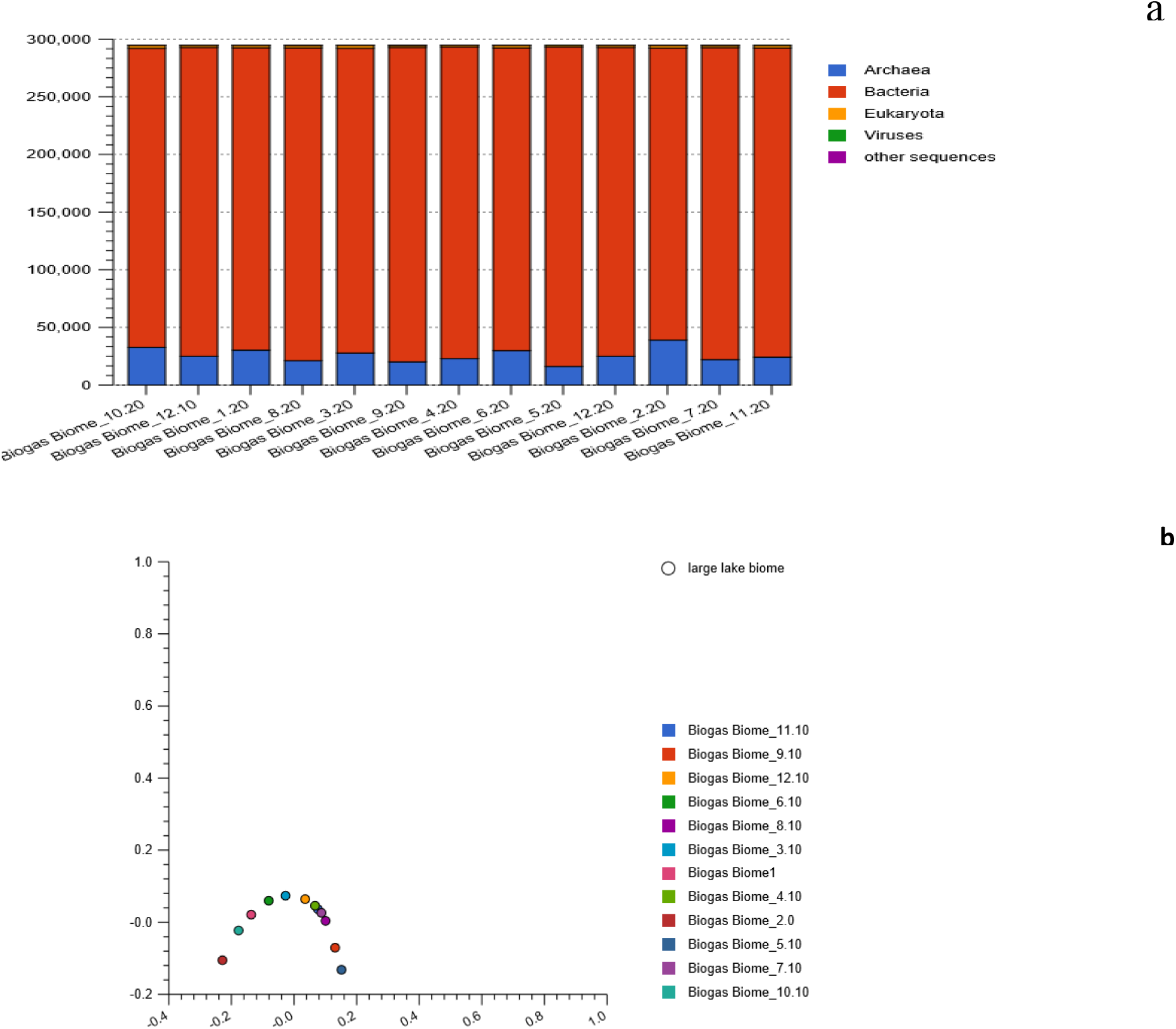
The stacked barchart (a) showing the three main domains, proportion of relative abundances and their PCoA plot (b) based on the Euclidean model. The PCoA plot revealed dissimilarities among the studied reactors. However, only three reactors partially clustered in the upper right quadrant of the plot. Though the Viruses nucleotide contributed to the curve they were not considered in the downstream analysis.

### The identified *Bacterial* biomes

We evaluated the variation in abundance and composition of 92 *Bacterial* domain orders that were assigned to 41 classes and 21 phyla (SFig. 2a, Table 1). Generally we observed significantly higher *Bacterial* abundances (89-93% of the total hits), compared to other domains. Similarly, significantly high β-diversities of the *Bacteria* communities were also observed at the phylum level. However, the communities of reactor 2 and 10 (within the same block) and reactor 4 and 9 (on different blocks) were found to reveal low β-diversity (SFig. 2b). In addition, we further observed low divesity variabilities between communities of reactor 3 and 6 and those of reactor 8 and 11 when compared at lower (class and order) taxonomic ranks (SFig. 3a and 3b). Hence, only four out of twelve treatments had high β-diversities, which could indicate genetic plasticity due to species evolution.

Specifically, we revealed *δ-Proteobacteria* species as the most abundant guild, constituting 9-14% of the total microbial communities. Globally the species were the second most abundant after the *Clostridial* species that composed 11-26% of the total communities. The *δ-Proteobacteria* communities were classified into 7 orders (SFig. 6a). Among them were *Syntrophobacteriales* that comprised 2-4% of the total identified communities, followed by *Desulfuromonadales* and *Myxococcales* species that comprised 2-4% and 0.9-4% of the total microbial compositions respectively. *Desulfobacterales* and *Desulfovibronales* communities were found to comprise 1-2% of the total species abundances. Among the least abundant *δ-Proteobacterial* species were the affiliates of *Desulforculales* and *Bdellovibrionales* that comprised only < 1% of the identified total communities. The non-classified reads derived from the class *δ-Proteobacteria*, also composed < 1% of the total communities. However, the PCoA plots indicated high sample to sample variation of the *δ-Proteobacteria* communities, although the composition of reactor 4 and 7 were found to be similar (SFig. 6b).

Among the identified species, the affiliates of *Firmicutes* were the most abundant in seven out of twelve reactor treatments and comprised 15-34% of total hits (Table 1) while the *Proteobacteria* species (Fig. 3) dominated in the other five treatments, comprising of 19-32% of total hits, (Table 1). *Proteobacteria* richness was significantly higher compared to that of *Firmicutes* species. The *Proteobacteria* communities were classified into 38 orders, which were affiliated to 7 classes (SFig. 4a). We linked their versatility with their ability to circumvent the oxidative stress in the systems. Significantly high levels of similarity of microbial composition were revealed among the treatments 1, 3 and 6, and treatments 4, 7 and 12, all located on the upper and lower right quadrant of the plot, (SFig. 4b and 5; at the class and order level). However, low β-diversities were observed for the identified *Proteobacteria* communities of reactor 4, 7, 8 and 9, (SFig. 5) at the order level. Other treatments were found to have high β-diversities of *Proteobacteria* species.

**Fig. 3:**
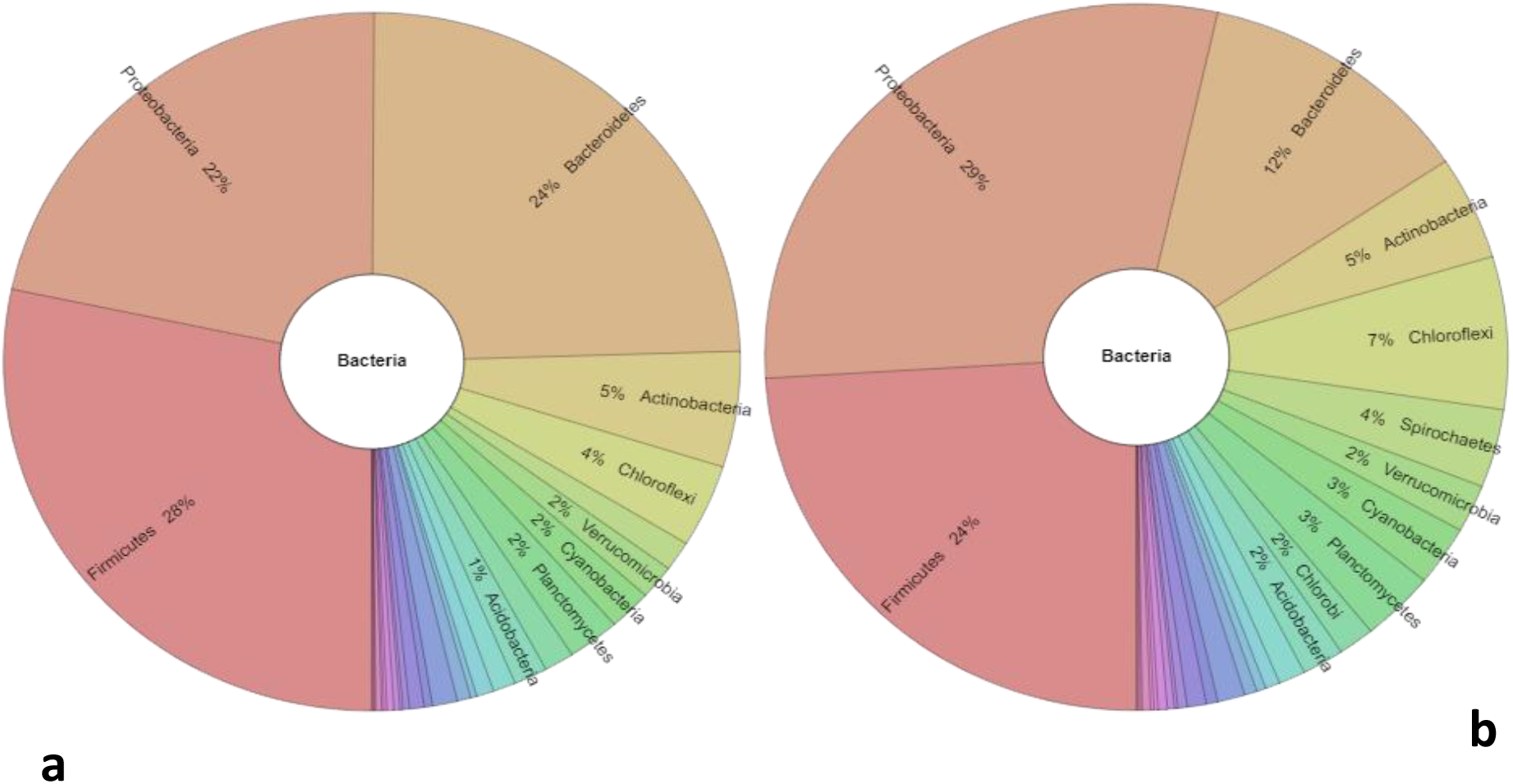
The Krona radial space filling display showing the source evidence of the stated relative abundances of the identified communities in the text (e.g Table 3 and within the text). The display reveals the abundances of the Bacterial reads for the communities identified in reactor 12 and 3: a) Reveals community abundances of reactor 12 while b) reveals community abundances of reactor 3, where *Firmicutes* and *Proteobacteria* dominated.

The communities’ affilliated to the class *γ-Proteobacteria* comprised 4-11% of the total identified reads. The affiliates were the fifth most abundant group in the entire microbiome and were classified into 14 orders (SFig. 7a). Among the identified orders was *Pseudomonadales*, that dominated the class, comprising of <1 to 6% of the total identified communities. This phylotype was followed by *Alteromonadales* and *Enterobacterales* species in abundance. The members of *Chromatiales*, *Vibrionales*, *Methylococcales*, *Oceanspirillales* and *Pasteurelales* respectively were also among the detected species. Other identified orders were *Xanthomonadales*, *Thiotrichales*, *Legionellales*, *Aeromonadales*, *Acidithiobacillales* and *Cardiobacterales*, all comprising <1% of the total identified communities. Unclassified *γ-Proteobacteria* reads were also identified in this study. Our results also revealed significantly high β-diversity (*P≤ 0.05*) of the microbial communities in the following six reactor treatments (reactor 2, 5, 6, 10, 11, 12). In contrast, the communities identified in reactors 1, 3, 4, 7, 8 and 9 were found to cluster on the upper right quadrant of the PCoA plot (SFig. 7b), indicating homogeneity and genetic composition similarity within the microbial populations.

Communities of *β-Proteobacteria* were detected in low abundances (2-4% of the identified microbial communities). Affiliates of *Burkholderiales* comprised of 1-2% of the microbial populations while the affiliates of *Rhodocyclales*, *Neisseriales*, and *Nitrosomonadales* species were found to comprise of <1% of the total identified populations across the treatments. Other identified *β-Proteobacteria* species were assigned to the order *Methylophilales*, *Gallionellales* and *Hydrogenophilales* which comprised of <1% of the total identified communities in the twelve treatments. High sample to sample variation was also observed for the *β-ProteoBacteria* species (SFig. 8b).

The affiliates of *α-Proteobacteria* were revealed to comprise of 3-8% of the total microbial communities, and were classified into 7 orders (SFig. 9a). The *Rhizobiales* dominated this group, comprising of 1-2% of the total microbial population in the treatments. The relative abundances of *Rhodobacterales*, *Rhodospirillales*, and *Sphingomonadales* followed (representing <1% of the total population). Other *α-Proteobacteria* orders identified included *Caulobacterales*, *Rickettsiales* and *Parvularculales*. We also found significantly high dissimilarities among the *α-Proteobacteria* within the studied treatments. Only the *α-Proteobacteria* communities of reactor 3 and 11 were revealed to express low β-diversity (SFig. 9b), a trait that we postulate was attributed by similar reactor operating conditions rather than the observed environmental conditions (Table 1 and 2).

**Table 2:** The Geographic positioning systems’ co-ordinate for the sampled reactor treatments, assigned to the four blocks of eastern and central regions of Kenya.

| Reactor Number | Sample id | Blocks | Treatment | Northings | Eastings | Altitude | Location |
| --- | --- | --- | --- | --- | --- | --- | --- |
| EA12/11/09/004 | S_1 | Kiambu | L-C | S(00°15.477') | E (37°38.828') | 1468 M | Ngenda |
| EA12/10/07/003 | S_2 | Kiambu | S-C | S(00.21347°) | E(37.61867°) | 1674 M | Karimoni |
| EA12/10/04/001 | S_3 | Meru | S-E | S(00°08.180') | E(37°39.477') | 1443 M | Kaithe |
| CE20/12/01/020 | S_4 | Kirinyaga | S-C | S(00°32'45.97") | E(37°26'40.52") | 1364 M | Murinduko |
| EA12/10/08/004 | S_5 | Meru | L-E | N(00°01.997') | E(37°35.353') | 1958 M | Kithaku |
| EA12/10/09/002 | S_6 | Meru | M-E | N(00°04.046') | E(37°38458') | 1650 M | Ntima |
| EA12/11/12/002 | S_7 | Meru | L-E | N(00.02946°) | E(37.10077°) | 1987 M | Ontulili |
| EA14/11/05/001 | S_8 | Embu | L-E | S(00°033.269') | E(37°28.247') | 1306 M | Municipality |
| EA14/11/09/001 | S_9 | Embu | M-E | S(00°27.824') | E (37°32.537') | 1417 M | Gaturi South |
| CE12/10/06/012 | S_10 | Kiambu | L-C | S(00.24874°) | E(37.62438°) | 1589 M | Kianyau |
| CE12/10/07/013 | S_11 | Kiambu | S-C | S(00°16'17.57") | E(37°38'40.16") | 1577 M | Dururumo |
| CE14/11/11/001 | S_12 | Kirinyaga | M-C | S(00.48637°), | E (37.44057°) | 1539 M | Ngandori |

**Legend:** CE: Central regions, EA: Eastern regions, L-C-Large size, Central regions, M-C: Medium size, Central region, S-C: Small size, Central regions, L-E: Large size, Eastern regions, M-E: Medium size, Eastern regions S-E: Small size, Eastern regions.

The ε and *Z*-*Proteobacteria* were among the least abundant *Proteobacteria* phylotypes in our study and were found to comprise ≤1% of the total identified microbial populations in the treatments. The *ε-Proteobacteria* communities were classified into the orders *Campylobacteriales* and *Nautiliales* (SFig. 10a), both comprising <1% of the total identified populations. We found similarities between the *ε-Proteobacteria* communities of reactor 4 and 8 (clustered) and those of reactor 5 and 9 (clustered partially) revealing low β-diversity (SFig. 10b). All the identified *Z*-*Proteobacteria* were from the order *Mariprofundales*, and accounted for <1% of the total microbial populations in the studied treatments.

The *Firmicutes* communities were the second most diverse community identified in this study. They comprised of 15-34% of the total microbial communities identified. The community was classified into 8 orders and 4 classes (SFig. 11a and 12a). The *Firmicutes* in the three treatments of reactor 2, 10 and 12 revealed significant genetic similarity and were revealed to cluster in the lower left quadrant of the PCoA plot (SFig. 11b). The *Firmicutes* in reactor 1 and 7 were found to cluster partially on the upper right quadrant of the PCoA plot (significant threshold; *P≤0.05*; SFig. 11b), indicating low β-diversity. At the order level, the identified *Firmicutes* of reactor 2 and 10; 3 and 6; 7 and 12; and reactor 1 and 9 also exhibited low β-diversities (SFig. 12b). Other local communities were revealed to express high β-diversities at class and order levels.

The *Clostridia* was revealed to be one of the most abundant *Firmicutes* phylotype, comprising 8-22% of the identified total microbial communities. It was classified into 4 orders (SFig. 13a). The order *Clostridiales* dominanted the phylum and constituted 8-22% of the total communities. *Thermoanaerobiales* species constituted 2-3% of the total identified communities. Other identified *Clostridia* species were the affiliates of *Halanaerobiales* and *Natranaerobiales*, all represented <1% of the total microbial populations, except for the *Halanaerobiales* communities in reactor 11 that comprised of 2% of the total identified microbial populations. Among the twelve biogas reactor treatments, three treatments (i.e reactor 7, 11 and 12) clustered on the lower left quadrant of the PCoA plot, (SFig. 13b), indicating high community similarities. Communities of reactor 5 and 9 were found to cluster partially on the upper left quandrant of the PCoA plot, revealing significantly low β-diversity.

The other seven treatments had high β-diversity, implying metabolic and genetic plasticity among the identified *Clostridia* species. The class *Bacilli* comprised of 4-9% of the total identified communities and was the fourth most abundant *Clostridia*. The *Bacilli* were classified into *Bacillales* and *Lactobacillales* (SFig. 14a), which Comprised 3-4% and 1-4% of the total identified microbiome communities, respectively. Among the *Bacilli* communities, those of reactor 5 and 8 were found to exhibit significant dissimilarities, while the others exhibited a low β-diversity (SFig. 14b). Other detected low abundance *Firmicutes* were *Erysipelotrichales* (*Erysipelotrichi*) and *Selenomonadales* (*Negativicutes*) which comprised of <1% of the total identified communities. The *Erysipelotrichales* species in reactor 5 and 9 were however found to comprise of 1% of the total identified microbial populations.

*Bacteroidete*s were the third most abundant phylotype comprising of 10-22% of the total identified microbial populations. These were classified into 4 classes and orders (SFig. 15a and SFig. 16a). Among them, were the *Bacteroidia* that comprised of 5-12% of the identified microbial communities. All the *Bacteroidia* belonged to the order *Bacteroidales*. Other members of *Bacteroidete*s were *Flavobacteria* that belong to the order *Flavobacterales*. The communities of *Cytophagia* were all assigned to *Cytophagales* species and they comprised of 1-2% of the total identified microbial communities. The other *Bacteroidete*s were *Sphingobacteria*, mainly of the order *Sphingobacteriales*, which comprised of 2-3% of the total identified microbial communities. We observed partial clustering of the *Bacteroidete*s’ communities particularly in reactor 2 and 10 and in reactor 8 and 12, which indicated low β-diversity between the respective communities (SFig. 15b and 16b). The *Bacteroidete*s communities of reactor 4 and 9 also clustered on the lower left quandrant of the PCoA plot, (SFig. 15b and 16b), implying significant similarities of the *Bacteroidete*s in the two treatments.

We identified 6 orders of *Actinobacteria* species (SFig. 17a) which comprised of 3-7% of the identified microbial reactor communities. The *Actinomycetales*’ were among the identified affiliates, comprising of 2-6% of the total identified communities. The species of order *Coriobacteriales* and *Bifidobacteriales* accounted for <1% of the total identified microbial composition. Other less abundant *Actinobacteria* species were assigned to the order *Rubrobacteriales*, *Solirubrobacteriales* and *Acidimicrobiales*. Out of the twelve biogas reactor treatments, the microbial communities of the seven treatments (reactor 1, 3 and 12; reactor 5 and 11 and reactor 7 and 8; SFig. 17b) were found to exhibit significantly (*P≤0.05*) low β-diversities. Only *Actinobacteria species* of the remaining three out of the twelve treatments were found to exhibit high β-diversities.

The *Chloroflexi* were categorized into 4 classes (SFig. 18a) and 5 orders (SFig. 19a) and comprised of 2-7% of the total microbial communities identified (Table 3). The communities belonged to Chloroflexi (1-4% of the identified communities) and *Dehalococcoidetes* (≤1-3% of the identified communities) as the most abundant phylotypes. Although all the *Dehalococcoidetes* reads were unclassified at a lower level, we identified *Chloroflexales* and *Herpetosiphonales* as the affiliates of the class *Chloroflexi* (SFig. 20). The two orders comprised of 1-2% and <1% of the total identified microbiomes, respectively. Other identified classes of the phylum were *Thermomicrobia* and *Ktedonobacteria*; all of which comprised of <1% of the total identified communities. The *Thermomicrobia* were classified into orders *Sphaerobacterales* and *Thermomicrobiales* (SFig. 21a). All the identified *Ktedonobacteria* belonged to the order *Ktedonoacteriales* (also representing <1% of the identified communities). We identified two treatments (reactor 1 and 7) that revealed low β-diversities of their local *Chloroflexi* communities (SFig. 18b). Four treatments (reactor 4, 5, 10 and 12), out of the remaining ten were revealed to also cluster on the lower right quandrant of the PCoA plot (SFig. 18b and 19b).

**Table 3:** The ArcGIS Software and the Questionnaire generated environmental variables and Phenotypic traits for the sampled treatments within the identified blocks of Kenya.

|  | Alt | S-T | AEZ | Ann. Av. R-f | Ann. Mean Temp °C | Substrate source |  |  | Family size |  |  | Reactor Performance |  |  |  |  |
| --- | --- | --- | --- | --- | --- | --- | --- | --- | --- | --- | --- | --- | --- | --- | --- | --- |
| Sample id |  |  |  |  |  | CO | PO | PI | M | F | T | IG | ACG | RV | FACoG | GGV |
| S_1 | 1468 M | Lithosol | UM 4 | 800- 900 | 20.7-19.9 | 4 | - | - | 5 | 4 | 9 | 4.8 | 4.0 | 16 M<sup>3</sup> | 14 M<sup>3</sup> | 2 |
| S_2 | 1674 M | Ferralsol | UM 2 | 1100-1400 | 19.5-18.4 | 3 | - | - | 2 | 1 | 3 | 2.4 | 2.0 | 8 M<sup>3</sup> | 7 M<sup>3</sup> | 1 |
| S_3 | 1443 M | Nitosol | UM 3 | 1400-2200 | 20.6-19.2 | 11 | - | - | 3 | 4 | 7 | 2.4 | 2.0 | 8 M<sup>3</sup> | 5 M<sup>3</sup> | 3 |
| S_4 | 1364 M | Andosol | UM 2 | 1220-1500 | 20.1-19.0 | 6 | - | - | 4 | 2 | 6 | 1.8 | 1.5 | 6 M<sup>3</sup> | 4.5 M<sup>3</sup> | 1.5 |
| S_5 | 1958 M | Phaeozem | L H 1 | 1700-2600 | 17.4-14.9 | 3 | - | - | 3 | 1 | 4 | 4.8 | 4.0 | 16 M<sup>3</sup> | 12 M<sup>3</sup> | 4 |
| S_6 | 1650 M | Nitosol | UM1 | 1650-2400 | 19.2-17.6 | 2 | 30 | - | 2 | 2 | 4 | 3.0 | 2.5 | 10 M<sup>3</sup> | 6 M<sup>3</sup> | 4 |
| S_7 | 1987 M | Phaeozem | LH 1 | 1700-2600 | 17.4-14.9 | 5 | - | - | 5 | 2 | 7 | 4.8 | 4.0 | 16 M<sup>3</sup> | 10 M<sup>3</sup> | 6 |
| S_8 | 1306 M | Nitosol | UM 3 | 1000-1250 | 20.7-19.6 | 5 | - | 6 | 5 | 4 | 9 | 4.8 | 4.0 | 16 M<sup>3</sup> | 11 M<sup>3</sup> | 5 |
| S_9 | 1417 M | Andosol | UM 2 | 1200-1500 | 20.1-18.9 | 2 | - | - | 3 | 2 | 5 | 3.0 | 2.5 | 10 M<sup>3</sup> | 8 M<sup>3</sup> | 2 |
| S_10 | 1589 M | Acrisols | UM 2 | 1100-1400 | 19.5-18.4 | 3 | - | - | 2 | 2 | 4 | 4.8 | 4.0 | 16 M<sup>3</sup> | 13 M<sup>3</sup> | 3 |
| S_11 | 1577 M | Cambisol | UM 3 | 800-1200 | 19.9-19.5 | 14 | - | - | 5 | 4 | 9 | 2.4 | 2.0 | 8 M<sup>3</sup> | 6 M<sup>3</sup> | 2 |
| S_12 | 1539 M | Ferralsol | UM1 | 1400-1700 | 19.3-17.5 | 3 | - | - | 4 | 3 | 7 | 3.0 | 2.5 | 10 M<sup>3</sup> | 8 M<sup>3</sup> | 2 |

**Key:** Alt: Altitude (M amsl); **S-T:** Soil type; **AEZ:** Agro-ecological zones; **UM 3:** Marginal Coffee zone; **UM 1:** Coffee-Tea Zone; **UM 2:** Main coffee zone; **LH 1:** Tea-Dairy Zone; **R-F:** Rainfall (mm); **Temp:** Temperature (°C); **CO:** Cow; **PO:** Poultry; **PI:** Pigs; **M:** Male; **F:** Female; **T:** Total; **IG:** Ideal gas (M<sup>3</sup>); **ACG:** Actual calculated gas (M<sup>3</sup>); **RV:** Reactor Volume (M<sup>3</sup>); **FACoG:** Farmers Actual Collected gas (M<sup>3</sup>); **GGV:** Difference between the RV and FACoG.

Similarly, the *Chloroflexi* communities of reactors 1 and 5 and those of reactors 3 and 7 were found to cluster on the upper right and lower right quandrants of the PCoA plot (SFig. 20b). The *Chloroflexi* class communities of reactor 1 and 10 were found to cluster partially with the cluster of microbes from reactor 3 and 7. We also identified partial clustering of *Chloroflexi* from reactor 8, 11 and 12 indicating low β-diversities among the communities in these reactors (SFig. 20b). The *Chloroflexi* from the rest of the treatments revealed high β-diversities. In contrast, the *Thermomicrobia* communities were significantly dissimilar among the treatments as indicated in SFig. 21b.

The *Cyanobacteria* comprised of 1-3% of the total microbial composition and majority in this group were unclassified. Only <1% of the total population were assigned to *Gloeobacteria* (SFig. 22a). Unlike other phylotypes, only the *Cyanobacteria* communities of reactors 3, 7 and 10 were found to exhibit significantly high β-diversity (*P≤ 0.05*), (SFig. 22b) while the communities in the other treatments were found to express partial similarities between or among themselves. The *Gloeobacteria* communities were all classified as *Gloeobacteriales* order. Other identified communities (unclassified reads) included members of the following orders; *Chroococcales*, *Nostocales*, *Oscillatoriales* and *Prochlorales* (SFig. 23a). Overally, a majority of the local *Cyanobacteria* communities were found to reveal low β-diversities. The *Cyanobacteria* communities in reactors 4, 6, 9 and 12, however exhibited high β-diversities (SFig. 23b). Similarly the identified four phylotypes at the order level (derived from unclassified reads, SFig. 24a) were also revealed to express high local compositional dissimilarities among the twelve biogas treatments (SFig. 24b).

We also identified *Acidobacteria* which comprised of 1-2% of the total identified microbial communities. Majority of these populations were from the order *Acidobacteriales* (comprising <1-2% of the total identified microbial communities). The remaining species were affiliated to *Solibacteres*, particularly of the order *Solibacterales* (<1% of the total identified microbial communities) (SFig. 25a and 26a). The *Acidobacteria* communities of five (reactors 1, 2, 3, 8, 12) out of twelve treatments were revealed to exhibit low β-diversities. *Acidobacteria* communities of reactor 2 and 8 clustered on the upper left quadrant of the PCoA plot (at order level; SFig. 25b and 26b), indicating similarity between the two local communities. *Deinococcus-Thermus*, another phylum, had the orders *Deinococcales* and *Thermales* species (SFig. 27a), the two belonging to the class *Deinococci*. The relative abundances of the two orders were ≤1% of the total identified microbial populations. We observed low β-divesities between the *Deinococci* species of reactor 11 and 12 while the diversity of its communites in reactors 3 and 6 and those of reactors 7 and 9 were found to be significantly similar (SFig. 27b). All the other local *Deinococcus-Thermus* communities expressed high β-diversities.

The identified *Verrucomicrobia* communities were classified into 3 classes that comprised 1-2% of the total identified microbial communities. Among the identified classes were *Opitutae, Verrucomicrobiae* and *Spartobacteria* (SFig. 28a). The affiliates of *Verrucomicrobiales* were the most abundant followed by those of the *Puniceicoccales* and *methylacidiphilales* species; all representing <1% of the total identified communities (SFig. 29b). Notably, we found out that all the *Opitutae* and *Spartobacteria* reads were unclassified at a lower taxonomic level. Only the *Verrucomicrobia* communities of reactors 3 and 7 were found to cluster partially on the lower right quadrant of the PCoA plot, (at the class level, SFig. 28b). In contrast, several local communities including the ones identified in reactors 1 and 10, and reactors 3, 6, 7 and 11 were found to cluster partially (indicative of low β-divesities) within the plots (at order level; SFig. 29b). However, their communities in reactor 5 and 8 were found to cluster significantly on the upper right quadrant of the PCoA plot (SFig. 29b), implying high similarities between the *Verrucomicrobia* species in the two treatments. Other treatments were found to reveal high β-diversities. Among the identified *Tenericute*’s, were *Acholeplasmatales*, *Mycoplasmatales* and *Entomoplasmatales*, all comprising of <1% of the total identified microbial communities and belonged to the class *Mollicutes* (SFig. 30a). Interesting, member species of this class were obsolutely absent in reactor 5 only, contrary to other phyla species, probably due to natural selection pressure. Our data also revealed significantly high similarities of the identified *Tenericute*’s species inhabiting reactors 4 and 9 and reactors 6 and 8 (SFig. 30b). The local communities of the other treatments were observed to be significantly dissimilar.

Our results also identified *Planctomycetacia* affiliates which comprised 1-3% of the total identified microbial communities. These affiliates were all assigned to the order *Plantomycetacetales* (*Planctomycetacia*). *Chlorobia* were identified in our study (comprising of 1-2% of the total identified communities). These belonged to the order *Chlorobiales*. *Spirochaetes* were also identified (<1-4% of the total identified microbial composition). All the *Spirochaetes* identified in this study belonged to the order *Spirochaetales*. *Thermotogae* of the order *Thermotogales* were also identified in the study. Other less abundant communities identified were *Synergistales* (*Synergistates*, *Synergistia*), *Lentisphaerales* (*Lentisphaerae*, *Lentisphaerae*), *Fusobacterales* (*FusoBacteria*, *FusoBacteria*), and *Aquificales* (*Aquificae*, *Aquificae*), all of which comprised of <1% of the total identified communities. *DeferriBacteriales*, *Chlamydiales*, *Chrysiogenales*, *Nitrospirales*, *Fibrobacteriales*, *Elusimicrobiales*, *Dictyoglomales*, *Gemmatimonadales* were other groups of microbes identified in this study. The *Candidatus poribacteria* affiliates were not accurately classified at the lower taxonomic ranks (Table 3).

### The identified *Archaeal* biomes

We analyzed the identified *Archaea* domain and established that their species comprised of 6-12% of the total microbial populations. Out of the identified communities, we obtained 5 phyla (SFig. 31a), 9 classes (SFig. 32a) and 16 orders (SFig. 33a). Among the communities, those identified in reactor 4 and 12 were found to cluster on the lower left quadrant of the PCoA plot, indicating their high similarity. We also revealed low β-diversities of the identified *Archaeal* communities, of reactor 3 and 6 and also those detected in reactor 7 and 9 (Phylum level; SFig. 31b). However, at a lower taxonomic level, only those identified in reactor 3 and 7 were observed to have low β-diversity (Class and order level; SFig. 32b and 33b). All the other reactor treatments were observed to contain highly dissimilar communities of *Archaea* at different taxonomic ranks.

A majority of *Archaea* species identified in this study were classified as *Euryarchaeota* and comprised of 5-11% of the total microbial communities. The *Euryarchaeota* communities were classified into 8 classes (SFig. 34a) and 10 orders (SFig. 35a). Interestingly, we observed significant dissimilarities (high β-diversity) of the *Euryarchaeotes* among the reactor treatments at the two taxonomic levels of class and order (SFig. 34b and 35b). The class *Methanomicrobia* was the most abundant *Euryarchaeota* phylotype and comprised of 4-9% of the total identified microbial populations. Further, we established that the majority of the members of the class were affiliates of *Methanomicrobiales* order (2-4% of the identified total communities), followed by those assigned to *Methanosarcinales* and *Methanocellales* (SFig. 36a), which comprised 1-3% and ≤1% of the identified total microbial communities, respectively. The local composition of *Methanomicrobia* species were highly dissimilar among the reactor treatments except for those from reactor 1 and 7 that were found to cluster. The communities of the two treatments revealed significant similarities (significant threshold, *P≤ 0.05*; SFig. 36b). Other identified methanogenic communities were assigned to *Methanobacterales*, the class *Methanobacteria*; *Methanococcales* belonging to *Methanococci* class; and *Methanopyrales* the affiliates of *Methanopyri* phylotype. All these species were revealed to comprise of <1% of the detected microbial communities.

Other non-methanogenic *Euryarchaeotes* identified in our study were the affiliates of *Thermococcales* (*Thermococci*), *Archaeglobales* (*Archaeglobus*), *Halobacterales* (*Halobacteria*), and *Thermoplasmatales* (*Thermoplasmata*). Among these phylotypes of *Euryarchaeotes*, affiliates of *Thermococcales’* were dominant while the *Thermoplasmatales* were the least abundant. *Crenarchaeota* and *Thaurmarchaeota* phyla were among the other identified non-methanogenic species. Between the two phyla, *Crenarchaeotes* were the most diverse and were classified into 4 orders (SFig. 37a). Within the identified *Crenarchaeota,* the following orders were identified; *Desulfurococcales*, *Thermoproteales*, *Sulfolobales* and *Acidilobales* (in order of abundance). All these orders are members of *Thermoprotei* class which comprised of <1% of the identified microbial communities. We noted that half of the studied treatments (reactor 6 and 9, reactor 7 and 10 and reactor 2 and 11) had *Crenarchaeota* communities that had low β-diversities, while the rest of the communities, in the other six treatments had significantly high β-diversities (significant threshold, *P≤0.05*; SFig. 37b). *Thaurmarchaeota*, another non-methanogenic phylum was classified into *Nitrosopumilales* and *Cenarchaeales* orders (SFig. 38a) both of which exhibited high β-diversities (SFig. 38b). Other identified phyla were *Korarchaeota* and *Nanoarchaeota*. The classification of these two phyla at the lower taxonomic ranks was inaccurate and the sequence reads from the two were therefore treated as unclassified.

### The identified *fungal* biomes

Our study also identified the presence of *fungal* species. The fungal communities were classified into 5 phyla (SFig. 39a), 14 classes (SFig. 40a) and 23 orders (SFig. 41a). Their relative abundances were however low and they comprised of <1% of the total microbial populations among the reactor treatments. Significantly high β-diversities were noted for the identified local communities of *fungi* at the phyla and class levels (SFig. 39b; SFig. 40b). However, at the lower taxonomic levels, low β-diversities were a common feature particularly for the communities identified in reactor 2 and 10, reactor 3 and 6, reactor 4 and 9, and reactor 8 and 11 (SFig. 41b).

The affiliates of *Ascomycota* were the most abundant guild (<1 of the total communities) within the *Eukaryotes* and were classified into 7 *Ascomycota* classes (SFig. 42a) that included *Eurotiomycetes*, *Saccharomycetes* and *Sordariomycetes,* respectively. Other identified classes were *Schizosaccharomycetes*, *Dothideomycetes*, *Leotiomycetes* and *Pezizomycetes* all of which comprised <1% of the total microbial communities in the respective treatments. We also further classified the identified *Ascomycota* species into 11 orders (SFig. 43a). Interestingly we observed significantly low β-diversity between *Ascomycota* communities in reactor 7 and 8. Other identified *Ascomycota* communities were observed to exhibit high β-diversities at the lower taxonomic levels (class and order level; SFig. 42b and 43b). Among the classes, *Sordariomycetes* were the richest within the phylum and their species were classified into 4 orders; the *Hypocreales*, *Sordariales*, *Magnaporthales* and *Phyllachorales* (SFig. 44a). The *Sordariomycetes*’ communities were found to exhibit significantly high β-diversities (Statistical threshold, *P≤0.05*) (SFig. 44b) among the studied treatments. Despite their higher abundance compared to *Sordariomycetes* phylotypes, the *Eurotiomycetes*’ another class of *Ascomycota* were classified into two orders; *Eurotiales* and *Onygenales* (SFig.45a). The two *Eurotiomycetes* orders were shown to comprise <1% of the identified total microbial populations. Like other fungal phylotypes, only the *Eurotiomycetes*’ communities of reactor 1 and 3 were found to cluster on the PCoA plot, which was indicative of high community similarity. The other reactor communities were revealed to have significantly dissimilar *Eurotiomycetes* populations (significant threshold, *P≤0.05*; SFig. 45b). Other identified fungal species were the affiliates of *Saccharomycetales*, *Schizosaccharomycetales*, *Pleosporales*, *Helotiales* and *Pezizales* phylotypes, all comprising <1% of the total microbial communities.

Among the identified fungi, *Basidiomycota* species were the second most abundant (<1% of the total communties). The species were classified into 4 classes and 5 orders (SFig. 46a and 47a, respectively). Among the classes, *Agaricomycetes* dominated the treatments, followed by *Tremellomycetes*, *Ustilaginomycetes*, and *Exobasidiomycetes* species (SFig. 46a) respectively. The affiliates of the class *Agaricomycetes* were further classified as *Agaricules* and *Polyporales* species (SFig. 48a). Other *Basidiomycota* species were affiliated to the following orders; *Tremellales* (*Tremellomycetes*), *Ustilaginales* (*Ustilaginomycetes*) and *Malasseziales* (*Exobasidiomycetes*). We observed two treatments’ (reactor 11 and 12) that contained *Agaricomycete*’s communities with low β-diversities (SFig. 48b) while the local *Basidiomycota* communities in the other reactor treatments had significantly high β-diversities (SFig. 46b and 47b).

The *Chytridiomycotes* were among the identified rare species. This was classified into *Chytridiomycetes* (more abundant) and *Monoblepharidomycetes* (SFig. 49). Among the *Chytridiomycetes* were *Spizellomycetales* (identified in reactor 9 and 10), *Rhizophydiales* and *Cladochytriales*, the latter two dectected only in reactor 9 (SFig. 50). All the affiliates of *Monoblepharidomycetes* were classified as *Monoblepharidales* species and were also detected in reactor 9 only. Other rare phylotypes identified in our study include *Blastocladiales*, (*Blastocladiomycetes*, *Blastocladiomycota*, detected in reactor 9), *Entomophthorales*, *Mortierellales*, and *Mucorales*, all derived from unclassified fungal reads (also identified in reactor 9, 10 and 11 respectively, SFig. 51). *Microsporidia* species were not accurately classisfied at the lower taxonomic levels and were regarded as unclassisfied though they comprised <1% of the total microbial populations.

### The microbiomes’ α-diversities and biogas production

Since environmental and technical variables can outweigh the biological variations when handling metagenomic datasets, prior to analysis, we asked ourselves, whether we had attained the maximum microbial richness and if our sampling was a representive of this maximum. To answer these questions, we conducted a rarefaction analysis through statistical resampling and plotted the total number of distinct species annotations against the total number of sequences sampled (Fig. 4). As a result, a steep slope, that leveled off towards an asymptote was obtained for each sample and later combined (Fig. 4). This finding indicated that a reasonable number of individual species were sampled and more intensive sampling of the same treatments would yield probably few additional species, if not none. Further the α-diversity indices revealed 947-1116 genera across the treatments, comfirming our maximum sampling effort was attained for further diversity studies. The study also revealed variabilities of reactor performance i.e biogas productivity (Table 3) among treatments, the phenotype that we linked to the observed α-diversity variability, among the treatments.

**Fig. 4:**
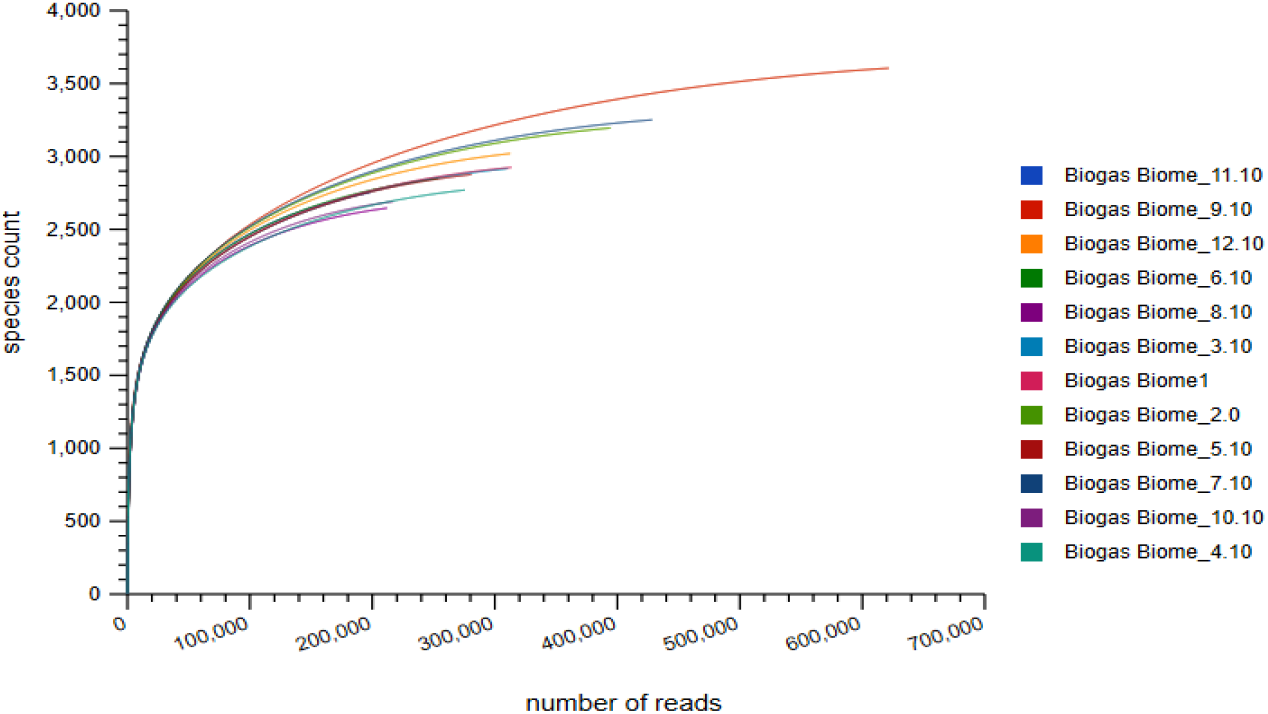
Rarefaction plot showing a curve of annotated species richness. The curve is a plot of the total number of distinct species annotations as a function of the number of sequences sampled.

### The microbiomes’ β-diversities and biogas production

To understand the influence of environmental variations and other variables on the local microbial communities and thus biogas production, we conducted comparative community diversity studies on the twelve biogas reactor treatments. As a result, we revealed high β-diversity among the local communities in the majority of the treatments (at the domain level). The only observed exceptions were the local communities of reactor 4 and 11 that clustered on the lower left quadrant of the plot (SFig. 52a), revealing significant similarity between the organisms in the two treatments. Similarly, at a lower taxa level (phylum, class and order), only the communities of reactor 2 and 10 were found to be similar, and were located in either the lower right or upper left quadrants of the PCoA plot (SFig. 52b and 52c). We also observed low β-diversity between the communities of reactor 4 and 9 at the lower taxonomic levels (phylum, class and order) with species clustering partially on the lower left and upper right quadrants of the PCoA (SFig. 52b and 52c), respectively. Other treatments that revealed low β-diversity between their communities were reactor 8 and 11 and reactor 3 and 6. Communities from these reactors were found to cluster partially in the lower right and lower left quadrants of the plot (Fig. 5). Similarly, we also noted significant variability of biogas production among the treatments (Table 3), an indication of the observed β-diversities at different taxonomic levels and within the taxa (stated above).

**Fig. 5:**
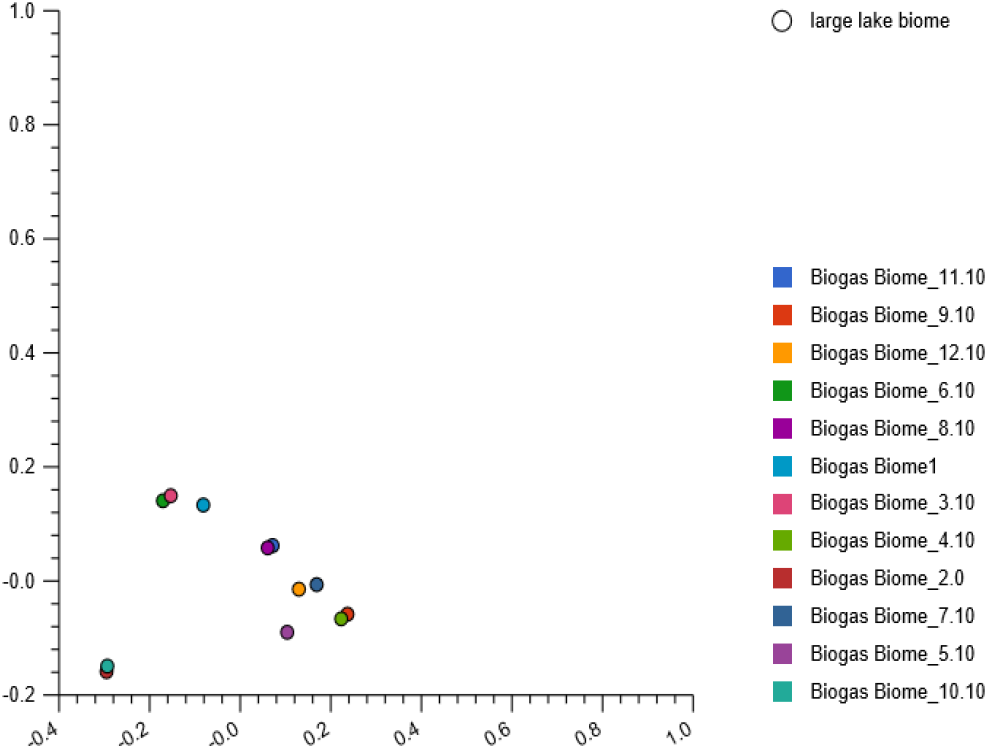
The PCoA analysis revealing β-diversity of the twelve treatments at the order level: The reads compositions in reactor 2 and 10 were partially similar, located in the lower left quadrant, those in reactor 8 and 11 located in the upper right quadrant, reactor 4 and 9 in the lower right quadrant, while those in reactor 3 and 6 were located in the lower left quadrant of the plot.

## Discussion

### Annotations against the 16 biological databases

The aim of our study was to use next generation sequencing technologies and different of bioinformatics tools to investigate the biogas microbiomes from environmental samples obtained from a wide geographic coverage. We combined marker genes and other genes on the assembled scaffolds to generate maximum taxonomic assignments. Among the sixteen utilized databases, SEED subsystems, yielded the highest reads hits annotations, the findings that corroborated with the previous reports [22]. Unlike other databases, SEED integrated several evidences that provided annotation benefits over the other 15 utilized databases. PATRIC database had fewer annotations compared to the IMG database, due to its high speciality of storing only pathosysstems information. The annotation of low protein sequences by the SWISS-PROT database was due to its manual curation and annotation systems. Contrasting the findings were the annotations of TrEMBL and Refseq databases that both utilized automated schemes, which superseded the annotation of SWISS-PROT database. However, in spite of its low resolution, majorly due to lack of regular updates, the outputs of SWISS-PROT are more accurate compared to the automated databases. Furthermore, orthologous groups databases such as COG, eggNOG etc were utilized to distigush orthologs from paralogs. In this case, we coupled gene classification schemes to the taxonomic assignements in order to increase detection resolution. Among these five databases (Fig. 1), eggNOG had the highest hits annotation due to its high phylogenetic resolution, automated annotation scheme, and coverage of more genes and genomes than its counterpart databases. In contrast, COG and NOG databases were among the databases that had the lowest hits annotations because of their manual curation and annotation schemes and fewer regular updates due to manual labor requirements. KEGG database was used to reconstruct metabolic pathways of the identified taxonomic assignements. Other utilized nucleotide databases were Genbank that ensured uniform and comprehensive annotation against the worldwide nucleotide sequences, Ribosomal Database Project (RDP) database that covered small rRNA gene subunit (SSU) of *Archaea* and *Bacteria* domains, LSU and SSU databases that archived large and small rRNA subunit of *Archaea*, *Bacteria* and *Eukarya* sequences respectively. The latter two; provided a complete species detection advantage over the RDP database. The two databases archived nucleotide sequences together with their postulated secondary structures. The SSU database stored more community sequences than its counterpart; the LSU database and it used prokaryotic and eukaryotic models for secondary structure alignments. The extra feature of the SSU database explained why the database, had the highest annotation resolution compared to other amplicon sequences databases. However, its worth-noting that the three rRNA databases archived complete or near-complete rRNA sequences, although partial fragments of 70% of the estimated chain length of the combined molecule were also included. These characteristics of the amplicon sequence databases limited the annotation of our Miseq generated short reads. The ENVO ontologies analysis, which revealed high resolution of large lake biomes, suggested that majority of the cultured and sequenced species were derived from large water masses.

### Annotations against the archived taxonomic profiles

Our taxonomic annotations approach generated more detection, compared to the other annotations conducted elsewhere [23]. While other studies revealed 0.042% to 0.132% rRNA genes, we revealed 280 (0.00%) to 1,032 (0.00%) rRNA genes. These findings implied that the proportion of the rRNA genes to the total shotgun generated genes were negligible. The observations were highly attributed to our sampling and sequencing approaches as previously explained by Knight *et al*. [24]. Furthermore, while most of the studies used 19-35% of the total generated sequences, [23], our approach annotated and used 39.75-46.70% of our total generated sequences. All these findings indicate that more than 53% of the biogas generating populations is still unknown. Briefly, our annotations were assigned to *Bacteria*, *Archaea*, *Eukaryotes* and *Viruses*, which corroborated the findings of other studies [25]. Among the domains, *Bacterial* species dominated in abundance, comprising 87-93% of the total microbial populations, followed by *Archaea* and *Eukaryotes*, while *Viruses* comprised the least abundant population. The observed high *Bacterial* abundance, suggest their crucial metabolic roles in biomass conversion and other reactions within the reactor systems [25]. As expected, we found low abundance among the *Eukaryotes* (*fungi*) which was indicative of the presence of low recalcitract biomasses concentration. Generally, the *fungal* species are involved in several biochemical reactions of biogas production including hydrolysis, fermentation and syntrophic reactions. The identified affiliates of *Archaea* were majorly consumers of smaller substrates that were generated by the *Bacterial* and *fungal* species. The *Archaea* species are able to use different methanogenic routes to convert the substrates into CH_4(g)_. Nevertheless, the main roles of the identified and less abundant group of the virus were unclear, though the species could have been active in degrading other microbial cells in the AD systems. However, their detection was largely attributed to the viral pathogenicity on both *Bacteria* and *Archaea* species. High β-diversity among the local communities (at the domain level) was an indication of environmental influences at the different ecological location.

### The identified *Bacterial* biomes

More than 90 representative orders of the *Bacteria* domain were revealed in our datasets. The first step of the AD process involves hydrolytic reactions that convert large macromolecules into smaller substrates [26]. Several *Bacterial* communities are capable of hydrolysis, even on high lignocellulosic plant biomasses. The identified species that participate in these reactions were annotated to the *Firmicutes*, *Bacteroidetes*, *Chloroflexi* to mention but a few and their species are known to harbor cellulosomes, that secrete extracellular enzymes, to degrade the substrate. Previous reports have indicated that species belonging to *Bacteroidales*, *Bifidobacteriales, Clostridiales Lactobacillales, Thermotogales* and many other identified *Bacterial* species catalyze the first step of AD [26]. The step involves rate limiting reactions, largely regulated by the substrate composition. The nature of the chemical composition of the substrate, generally depends on the environmental variables (Table 2 and 3), which influence microbial species’ abundances and β-diversity differently. All these affects reactor stability and performance. As expected, the class *Clostridia,* that comprised *Clostridiales*, *Thermoanaerobacteriale*s and *Halanaerobiale*s orders, was the most dominant among the identified phylotypes. The guild converts large macromolecules into organic acids, alcohols, CO_2(g)_ and H_2(g)_ [27–33]. In this case, the species use a substrate-level phosphorylation mechanism to conserve the energy. Acetogenic and homoacetogenic species belonging to the same guild such as *Moorella thermoacetica* and *Thermoanaerobacter kivui* are able to use the reductive acetyl-CoA pathway to reduce CO_2(g)_ into CO_(g)_, concommitantly generating reducing equivalents [34, 35] and vice versa. As a consequence, the acetogens and homoacetogens use proton or sodium dependent mechanisms to conserve their energy [35].

Above all, the *Clostridia* guilds are able to catalyze other biochemical reactions such as reductive dechlorination of various compounds that leads to production of sulfide from either thiosulfate or sulfite. The reactions consume pyruvate and lactate as the carbon source. In this case, the member species are able to use [NiFe]-hydrogenases to generate H_2(g)_ [33]. In our systems, we also identified *Halanaerobiales* species that have a capacity to tolerate high salt concentrations [36]. Generally the reactor treatments comprised high salts concentrations and the *Halanaerobiales* must have regulated the toxic cations from the ionization of the salts. All these roles of *Clostridia* suggest their metabolic versatility, and these features were used to explain the observed low richness of *Firmicutes* compared to *Proteobacteria* species. Similarly, the observed high β-diversities among the treatments were linked to the versatile roles of *Clostridia*. However, the observed low β-diversities (SFig. 13b) of communities in reactor 7, 11 and 12 are probably attributed to similar treatment perturbations and the identified environmental variables (Table 2 and 3).

The *Bacillales* and *Lactobacillales* species identified in our study, all belonged to the class *Bacilli*. These species are able to use mixed fermentation routes to metabolize carbohydrates and other biological macromolecules [37, 38]. Formate dehydrogenase complex activities are the main mechanisms of energy conservation [38] in these microbial guilds. Generally, *Bacilli* co-exist in a symbiotic relationship with *Clostridia* species in lactate and acetate metabolism [39], a mechanism that is termed as lactate cross-feeding. High β-diversities among the observed treatments (SFig. 14b) were solely attributed to cow-dung’s substrate composition and environmental variables (Table 2 and 3). However, the low β-diversities observed between or among the treatments (SFig. 14b) that formed the clusters were largely attributed to similarities of reactor’s operation conditions. All these implied that the respective organisms were in the same succession stage at the time of sampling. The *Erysipelotrichi* another identified *Firmicutes* class, utilize the syntrophic acetogenesis route to metabolize lipids into acetate, H_2(g)_ and CO_2(g)_ [23]. Its species, together with other *Firmicutes* species like *Smithella propionica* use a non-randomizing pathway [40] to metabolize propionate into acetate. Among other identified *Firmicutes*, were *Syntrophomonas wolfei* (*Clostridiales*, *Clostridia*) that use the β-oxidation pathway to oxidize C_4_ to C_7_ fatty acids in order to produce either acetate and H_2(g)_ or propionate, acetate and H_2(g)_ [41, 42]. Notably, these syntrophic species are consumers of 30% of the generated electrons, which bypasses the rate limiting steps of the volatile fatty acids accumulation. All these, contributes to reactor stabilities. In general, we associated the observed low *Firmicutes* abundances and diversities (richness, SFig. 11a and 12a) with genetic versitilities that conferred a capacity to encode an assortment of proteins; which metabolize a variety of biomasses into biogas. Furthermore, the observed high β-diversities (SFig. 11b and 12b) of the *Firmicutes* support our concepts on the genetic divergence among the species and we postulate that the trait was mainly influenced by the sizes of the reactors and the variability of the environmental conditions (Table 2 and 3).

The *Synergistete*’s species identified in this study have been established to ferment amino acids into lactate, acetate, H_2(g)_ and CO_2(g)_ [43, 44]; though they have no ability to metabolize carbohydrates and fatty acids. One such identified example is *Dethiosulfovibrio peptidovorans*. The affiliates of genus *Anaerobaculum* which belong to *Synergistete*s catabolize peptides into acetate and short-chain fatty acids while their close relatives, *Aminobacterium*, are capable of fermenting amino acids into acetate, propionate and H_2(g)_ [43, 44]. Other amino acid metabolizers identified in this study were the affiliates of *Fusobacteria* [45] which are able to catabolize substrates into acetate, propionate, and butyrates, concomitantly releasing H_2(g)_, CO_2(g)_ and NH_3(g)_. The member species of the two phyla are known to use the substrate-level phosphorylation to conserve their energy [46] from amino acids metabolism. Our findings contradict the metabolism of the amino acid by the *Clostridia* species, whose energy conservation mechanisms remain unclear and have been postulated to occur through chemiosmosis [30]. However, the two phyla described here have been found to generate energy through syntrophic association with methanogens; otherwise the generated protons would increase the systems’ alkalinity and lead to inhibited steady-state of the biochemical reactions. Based on the low abundance of the two phyla and richness among the treatments, we postulate low amino acid affinities among their identified species, which would suggest another selection pressure in the AD systems.

The identified affiliates of *Bacteroidales*, *Flavobacterales*, *Cytophagales* and *Sphigobacteriales*, all belong to *Bacteroidetes*, and are majorly known to ferment carbohydrates and proteins into mixed products (acetate, lactate, ethanol, and volatile organic acids), concomitantly releasing H_2(g)_ [47]. Moreover, their species are also capable of metabolizing aliphatic, aromatic and chloronated compounds [48] and were observed to co-exist with methanogens possibly to increase energy extraction from indigestible plant materials. The member species were highly varied among the treatments and we hypothesize that the cattle breeds and the operating conditions in the reactors were the key drivers of the observed β-diversities. Interestingly, we also observed consistency in the relative abundance (2% of total reads) of the *Cytophagale*’s guild among the different treatments, which may be attribitable to the long periods in conditions of low nutrient limitation [49]. Although, the metabolic activity of the identified *Chlorobi* overlap with that of *Bacteroidetes* [50], their species use reverse tri-carboxylic acid (rTCA) cycle to fix CO_2(g)_ and N_2(g)_ in anaerobic conditions. We hypothesize that these rTCA reactions may have been utilized by the *Chlorobi* to oxidize reduced sulfur compounds [51, 52]. In our results, the *Bacteroidetes* species however revealed a higher selection advantage over *Chlorobi* due to the observed low *Chlorobi* abudunces and richness (i.e with only a single order, the *Chlorobale*).

The identified *Proteobacteria* guilds consisted of metabolically versatile species and were revealed to comprise the highest number of species (richness) compared to other identified phyla in the treatments. The guilds comprised of species with capacity to metabolize proteins [53] carbohydrates and lipids [54] and we associated their low β-diversity (SFig. 4b and 5b) with low cow-dug’s substrate composition heterogeneity and differences in environmental conditions (Table 2 and 3). The *γ-Proteobacteria* was revealed in our study as the most diverse class of the *Proteobacteria* and the class comprised of 14 orders (SFig. 7). The *Enterobacteriales* included *Escherichia coli*, which are facultative anaerobes able to ferment carbohydrates into lactate, succinate, ethanol, acetate, H_2(g)_ and CO_2(g)_) [55]. Unlike other syntrophic species, the affiliates of *Enterobacteriales* have been found to use the methyl-citrate cycle [56] to metabolize propionate into pyruvate, a precursor for butanol, isopropanol and other mixed acid fermentation products. *Burkholderiales*, another *γ-Proteobacteria* phylotype identified in this study, produces succinate from carbohydrates concomitantly oxidizing H_2(g)_, S^0^ and Fe^0^ into their respective oxidized forms to conserve energy [57, 58]. We suggest that the high abundance and β-diversities of the *γ-Proteobacteria* communities (SFig. 7, Table 3) could be due to low environmental temperature.

In contrast, the affiliates of genus *Sutterella*, among other species of *β-Proteobacteria* identified in this study are known saccharolytic and nitrate reducers [58]. The species also consume propionate, butyrate and acetate syntrophically [54, 58]. We suggest that the high abundance and diversity of *β-Proteobacteria* (SFig. 8) in the treatments was favoured by the agroecological zones and prevailing environmental temperatures (Table 2 and 3) of the reactors. The *Caulobacterales*, an order of *α-Proteobacteria* were among the identified low abundant species in our study. These species have previously been revealed to survive in low nutrient environments [59]. *Rhodospirillales* (*α-Proteobacteria*), such as *Rhodospirillum rubrum*, have been previously determined to participate in homoacetogenesis and convert CO_(g)_ and formate into acetyl-CoA, CO_2(g)_ and H_2(g)_ [60]. The produced acetyl-CoA is further converted into acetate, a precursor for methanogenesis. Like members of *β-Proteobacteria*, the *α-Proteobacteria* species identified in this study were also influenced by the agroeclogical zones (Table 2 and 3) and we postulate that the cow-dug’s substrate composition was the main cause of the observed β-diversities and variations (SFig. 9) among the treatments. The *Campylobacterales* and *Nautiliales* species which are affiliates of *ε-Proteobacteria* were also identified in our study. These species metabolize proteins in the AD systems and have also been revealed to express an additional role of antibiotic resisistance in biofilms and granules [54].

Among the identified *δ-Proteobacteria* species, *Pseudomonadales*, the gemmules producing anaerobes [61] were the most abundant and have been found to co-exist with algae [61]. The reported syntrophic traits of the *Pseudomonadales* order were revealed to enable the species to withstand unfavourable environmental conditions, which could have offered the species the selection pressure advantage over other *δ-Proteobacteria* species. Generally, the *δ-Proteobacteria* species are syntrophic oxidizers [62–64], with the ability to convert volatile fatty acids into acetate, formate, CO_2(g)_ and H_2(g)_. The *Syntrophobacteriales* species identified in this study e.g *Syntrophus aciditrophus* are able to use the randomized methyl-malonyl-CoA pathway to oxidize butyrate, propionate and long-chain fatty acids into acetate, formate, CO_2(g)_ and H_2(g)_ syntropically [62, 63] while *Syntrophobacter wolinii* [64] another member species, is known to use the β-oxidation pathway to metabolize propionate into acetate. Other species belonging to the genus *Syntrophaceticus* identified in this study have been reported to oxidize the produced acetate into CO_2(g)_ and H_2(g)_ [65]. Interestingly, for these species to effeciently oxidize the respective macromolecules, they have been revealed to use an electron transport mechanism that creates a positive redox potential by reversing electrons that were generated in syntrophic reactions. In contrast, the identified affiliates of *Desulfarculale*’s, *Desulfobacteriale*’s and *Desulfovibrionale*’s phylotypes are sulfate reducers, although in the absence of sulfate (electron acceptor), the species are able to oxidize organic compounds, (e.g lactate, acetate and ethanol) into CO_2(g)_ and H_2(g)_ syntrophically [66]. The *Desulfuromonadale*’s species have been revealed to use Fe^3+^ compounds to oxidize ethanol into CO_2(g)_ and H_2(g)_, but in strict co-operation with methanogens [54]. Lastly, the *Myxococcales* and *Chlamydiae* species have been identified as fermenters of carbohydrates, [67] and proteins [68], which are ultimately converted into organic acids. All these *δ-Proteobacterial* roles contribute to the systems’ stability by providing electrons, which act as reducing equivalents in the AD systems. We also postulate that differences in the cow-dug’s substrate composition were the main drivers of the observed variations in the *δ-Proteobacteria* abundances and β-diversities among the treatments (SFig. 6b). The other low abundance species identified in our study were assigned to the *Mariprofundale*’s order, the affiliates of *Z*-*Proteobacteria* class. The affiliates co-exist with *Nitrosopumilales* to oxidize Fe^0^ [69] into Fe^2+^ or Fe^3+^ ions. Moreover, the two phylotype species have previously been revealed to convert nitrogen substrates into NO_3_^−^_(aq)_ in their syntrophy [70, 71]. In this study, through their syntrophy, we hypothesize a completion of redox reactions in the treatments through electrons release. In summary, all affiliates of *Proteobacteria* identified in this study are capable of reversing oxidative damage to methionine [54], a trait that explains why their species are more (richness) compared to other phyla species in our treatments. Moreover, *Proteobacteria* species are able to express shorter diversification time-scales over other microbes hence leading to large population sizes, high growth rates, and strong selective regimes of the species. All these factors facilitate rapid *Proteobacteria* adaptation through either mutation or recombination events.

We also identified *Solibacterales* and *Acidobacterales* species that were affiliated to the phyla *Acidobacteria*. Members of these species ferment polysaccharides and monosacharides into acetate and propionate [72]. The species are also able to fix CO_2(g)_ anaplerotically [73], ultimately replenishing carbon atoms in the AD systems. Furthermore, the affiliates are also able to catabolize inorganic and organic nitrogen substrates as their N-sources [74]. The detection of these species in our study is attributed to the high amount of NH_4_^+^_(g)_ in our AD systems. We postulate that reactor perturbations rather than environmental variables were the main influencers of the genetic composition of the *Acidobacteria* in our study (Table 3 and SFig. 25b and 26b).

Generally, the *Mollicute*’s species (*Tenericutes* phylum) detected in our study are facultative anaerobes. These were all absent in reactor 5, probably due to the nature of the substrates or feeds fed to the cattle breeds. The species are fermenters of sugars, and they convert the substrates into organic acids such as lactic acid, saturated fatty acids and acetate [75, 76]. All the identified species are affiliates of *Acholeplasmatales*, *Mycoplasmatales* and *Entomoplasmatales* which have previously been shown to exhibit reduced respiratory systems, with incomplete TCA cycle, that lack quinones and cytochromes. The metabolisms of *Tenericutes* generally yield low amount of ATP and large quantities of acidic metabolic end products [75]. It is probable that the produced acidic metabolites negatively affected the performance of other microbes in the systems in our study; which led to poor reactor performance. We hypothesize that the occurrence of these species in our systems was driven by the agroecological zones and soil types, which directly affected the nature of the cow-dug’s substrates fed into our treatments (Table 2 and 3 and SFig. 30).

Like other *Bacterial* species, the affiliates of *Actinobacteria* (SFig. 17a) identified in this study, are able to metabolize a wide variety of substrates [77, 78, 79] to produce acetic acid, lactic acid, H_2(g)_ and other products. Among the *Actinobacteria*, *Coriobacteriales* metabolize formate and H_2(g)_, and concomitantly reduce NO_3_^−^_(aq)_ into NH_3(g)_ [77]. *Slackia Heliotrinireducens* was one of the *Coriobacteriales* species identified in this study. Species of the *Bifidobacteriale* were also identified in our study. These species are able to metabolize oligosaccharides and release lactic acid [78]. Other species belonging to *Actinomycetales* syntrophically metabolize propionate into acetate [79]. The metabolism of NH_4(aq)_ by *Actinobacteria* is known to protect methanogenic *Archaea* from other oxidizing agents. However, their species have been reported to be more sensitive to nutrients perturbations [79] and due to this, reactor perturbations particularly the feeding regimes (SFig. 17b) may have contributed to the observed species abundances and diversities among our treatments in this study. The low abundance of *Plactomycete*s in this study has also been observed elsewhere [80]. *Plactomycete*s have been found to metabolize sulphate-containing polysaccharides [80], recalcitrant hydrocarbons and other organic compounds [81]. However, when in syntrophic association with *Chloroflexi* (SFig. 18a, 19a, 20a and 21a), the species are also able to oxidize butyrate [82] into smaller substrates.

The low abundance of communities of *Thermotogae* in this study is similar to findings of previous studies [83]. The carbohydrate metabolism roles of the *Thermotogae* were previously determined based on the gene content of *Defluviitoga tunisiensis* [84]. The species have been reported to convert the macromolecule into acetate, ethanol, CO_2(g)_, and H_2(g)_. In most cases, the species utilize the Embden-Meyerhof Parnas (EMP) pathway to generate pyruvate, which is either converted into acetyl-CoA, a precursor for acetate and ATP formation or reduced to ethanol, while at the same time oxidizing electron carriers. The H_2(g)_ produced in these reactions is coupled to the oxidation of NADH and reduction of ferredoxin [84]. The reactions have been found to favour proton production which reduces the *P*_*H2*_ in the AD systems. However, for these reactions to proceed favourably *Thermotogales* have been revealed to depend on *Clostridiales* species [85] which implies that the abundance and richness of *Thermotogale*’*s* is regulated by the *Clostridiales* species. The *Spirochaete*s were detected in low abundance in our study as was also found out by Wang and his co-wokers [86]. The identified *Spirochaete*s in this study are homoacetogens and acetogens which have previously been revealed to convert H_2(g)_, CO_2(g)_ and CO_(g)_ into acetate [86, 87]. These species have also been revealed to co-exist with methanogens to provide an alternative metabolic route, termed as the butyrate oxidation pathway for H_2(g)_ consumption [87]. Similarly, the *Spirochaetes* species have also been found to co-exist with the *Thermotogale*’*s* species and to jointly ferment tetrasaccharides and cellobiose into organic acids [88]. These syntrophies protect *Clostridiale*’s species from substrate inhibition mechanisms [85]. Based on our findings and those reported elsewhere, we suggest that numerous microorganisms identified in our AD systems co-operate through the use of an assortment of mechanisms to metabolize the substrates. Due to this co-operation, we further postulate the existence of a holobiontic mechanism in our treatments that is used to regulate symbionts and other syntrophic species through ecological selection, via co-evolution to minimize conflict between or among the symbiotic species.

The *Verrucomicrobia* communities identified in this study were affiliated to *Methylacidiphiles*, *Puniceicoccales* and *Verrucomicrobiales* orders (SFig. 28a and 29a). These communities have been established to metabolize cellulose and other C1-containing compounds [89]. The affiliates of *Cyanobacteria* were classified as *Chroococcales*, *Oscillatoriales* and *Nostocales* species (SFig. 23a and 24a) which are mainly involved in H_2(g)_ and N_2(g)_ metabolism [90–92] in addition to carbohydrate metabolism. Research has shown that the conversion of N-containing substrates into NH_3(g)_ is an endergonic reaction that requires ATP [90–92]. The reaction has also been found to oxidize H_2(g)_ into obligate H^+^_(aq)_ ions, circumventing high *P*_*H2*_ in the AD systems [90–92]. Therefore, the recycling of H_2(g)_ through the oxyhydrogen route, also termed as the Knallgas reaction by the *Cyanobacteria* species has been revealed to provide ATP and protect strict anaerobes and other enzymes from oxidizing agents, by providing reducing equivalents and removing toxic O_2(g)_ species [93, 94]. In our case, we attributed the variation in abundance of *Cyanobacteria* species among the treatments to the variability of H_2(g)_ concentration among the treatments.

Another identified phylotype in our study was the *Candidatus poribacterial* species. These species ferment carbohydrate and proteins, including urea [95] into organic acids. The species have been revealed to use the Wood-Ljungdahl pathway also known as CO_2(g)_ reductive pathway to fix CO_2(g)_ generated from carbohydrates and other macromolecules. The species have also been previously reported to utilize the oxidative deamination pathway [96] to ferment amino acids, and the randomized methyl-malonyl-CoA pathway to metabolize propionate. All these reactions produce H_2(g)_ and we postulate commensal symbiontism between *C. poribacteria* and acetogenic species in our treatments. Like other *Bacteria* species, *Fibrobacter* members such as *Fibrobacter succinogenes* identified in our study were likely to have been involved in the metabolism of cellulose [97] and other macromolecules in our AD systems.

Although it’s not a common tradition to explain the existence of metal-reducing organisms in the AD systems, it is worth noting that the electron sink ability of any system depends on these organisms. One of such phylotypes identified in our study were the affiliates of *Gloeobacterales* order (SFig. 22a and 23a). The member species have previously been reported to transfer electrons directly from methanogens to syntrophic *Bacteria* [98]. These kinds of reactions are thought to enhance clean biogas production through the inhibition of sulfate reducing bacteria. Other identified metal reducing species in our study were the members of *Clostridiales* that have previously been reported to utilize Fe^3+^ Co^3+^ and Cr^6+^ as electron acceptor [98]. Other identified *Clostridiales*’ species have also been reported to use the acrylate pathway to reduce arsenate and thiosulfate concomitantly fermenting organic molecules [99]. The affiliates of *Nitrospirales* (*Nitrospira* phylum) identified in this study, have been reported to reduce sulfate using H_2(g)_ and thereby promoting acetate oxidation [100, 101]. We hypothesize that all these metal reducing reactions contributed to the reactor stability through the provision of reducing equivalents in the systems.

### The identified *Archaeal* biomes

The generated volatile organic acids, CO_2(g)_ and H_2(g)_ by acidogens, acetogens, and homoacetogens are further metabolized by *Archaea*, particularly the species that generate CH_4(g)_ [102]. The *Archaea* domain constituted of six (6) methanogenic and ten (10) non-methanogenic guilds identified in our study, with the two groups playing equally important roles in the AD systems. The methanogenic species are consumers of *Bacterial* products and convert them into CH_4(g)_, the biogas. The methanogenic and non-methanogenic species identified in this study are affiliates of *Euryarchaeotes*, which utilize hydrogenotrophic, acetoclastic, methylotrophic and methyl-reducing pathways to metabolize the substrates of *Bacterial* species into CH_4(g)_. Generally, the species of *Eurychaeota* are known to indirectly stabilize the AD systems by consuming H_2(g)_ [103] and we suggest that the varied H_2(g)_ concentrations among the treatments, contributed to the observed variances in *Euryarchaeota*’s abundance and β-diversity among the local communities. However, environmental (Table 1 and 2) and reactor perturbations may also have influenced the community abundance. Among the methanogens, *Methanosarcinales*, the affiliates of *Methanomicrobia* class, were the second most abundant group, consisting of 1-3% of the total communities identified. The affiliates of *Methanosarcinales* use three metabolic routes singly or in combination to convert methanol and methylamines, acetate or CO_2(g)_/H_2(g)_ and other potential substrates into CH_4(g)_. The produced H_2(g)_ from the reactions is also consumed by the same species to maintain *P*_*H2*_ [104]. However, some species within the guild, are known to lower the accumulated *P*_*H2*_ through the reassimilation of acetate, a process that transfers H_2(g)_ to acetogens. Under certain conditions, the species of *Methanosarcinales* (e.g *Methanosarcina acetivorans C2A* and *Methanosarcina barkeri*) and *Methanobacteriales* (*Methanothermobacter thermoautotrophicus*) have been reported to express metabolic features similar to acetogenic organisms. Previously, these species have been reported to use the acetyl-CoA pathway to metabolize CO_(g)_ into acetate and formate, under different *P*_*CO(g)*_ [105]. These reactions provide an alternative route for ATP generation from CO_2(g)_ reduction by the generated H_2(g)_. Normally, the *Methanosarcinales* couple methanogenesis to ion transport, also termed as chemiosmotic coupling or chemiosmosis, to establish an electrochemical gradient across the cell membrane [105] because the CO_2(g)_ reduction reaction is normally unfavourable. The *Methanosarcinales* have however been reported to express significant metabolic plasticity.

Among the *Archaea* in our study, *Methanomicrobiale*’s, affiliates of the *Methanomicrobia* class (SFig. 36a) were the most abundant within the class, comprising of 2-4% of the identified total microbial populations. The affiliates of *Methanomicrobiale*’s use two methanogenesis pathways; hydrogenotrophic and methylotrophic routes, to convert formate, H_2(g)_/CO_2(g)_, 2-propanol/CO2_(g)_ and secondary alcohols [106] into CH_4(g)_. The *Methanomicrobiales* consume isopropanol to yield acetone and the reducing equivalent, NADPH [107]. However, the species are known to lack the CO_2(g)_ reductive pathway, and hence uncouple methanogenesis from CO_2(g)_ fixation, which implies that the species uses electron birfucation to conserve energy. Due to these characteristics, the *Methanomicrobiales* species are able to express the high acetate requirement phenotype for carbon assimilation [108]. In costrast, the identified *Methanocellale*’s species, which are affiliates of the same class, solely use the acetoclastic route to convert CO_2(g)_ into CH_4(g)_ [109]. Majority of the *Methanocellale*’s species have been reported to reduce CO_2(g)_ with H_2(g)_, except in rare cases where formate and secondary alcohols are used as alternative electron donors. Similarly, all the identified *Methanobacteriales*’ species, the affiliates of *Methanobacteria* class use the CO_2(g)_ reductive route to convert CO_2(g)_ into CH_4(g)_ with formate and H_2(g)_ acting as electron donors. We are prompted to postulate that formate metabolism particularly in acetoclastic methanogenesis was triggered by the low *P*_*H2*_ and that through electron bifurcation the species were able to conserve their energy. Other detected methanogenic species were the affiliates of *Methanopyrales*, belonging to *Methanopyri* class and *Methanococcales* species that were assigned to *Methanococc*i class (SFig. 34a and 35a). The members of the two classes have previously been reported to metabolize H_2(g)_ and CO_2(g)_ into CH_4(g)_ under thermophilic and saline conditions, [110, 111], which may indicate that our treatments had high amounts of salts and that the temperatures were high enough to support the species.

In contrast, the non-methanogenic group of *Archaea* generally reduce metal elements and their compounds [98, 99], to provide reducing equivalents in the AD systems. The affiliates of *Halobacteriales* (*Halobacteria*, *Euryarchaeota)* were among the detected non-methanogenic species in this study. The member species of this group of non-methanogens use the arginine deiminase pathway [112, 113] or other metabolic routes to catabolize amino acid substrates. The *Halobacteriales* conserve energy when the intermediate metabolites; ornithine and carbamoyl-phosphate, are further converted into CO_2(g)_ and NH_3(g)_. The significant differences in relative abundances of the *Halobacteriales* among the treatments can be attributed to the concentration of the alternative electron acceptors and salts concentrations in the systems. However, other deterministic factors such as competition and niche differentiation could also have led to the observed variations in *Halobacteriales* abundance among the treatments.

Our study also identified *Thermoplasmatales*, the affiliates of *Thermoplasmata* class. These groups of *Archaea* utilize sulfur for energy consevation [114]. However, their close relatives, *Methanomassiliicoccales*, formerly *Methanoplasmatales* have been found to use the methyl-reducing pathway to convert methanol into CH_4(g)_. These relatives conserve energy through electron bifurcation [115] and we postulate that the affiliates of *Thermoplasmatales* are also involved in CH_4(g)_ production although few or none of the biochemical assays have previously been conducted to test their biogas production abilities. We also identified *Thermococcales* species, of the class *Thermococci*. The affiliates of this class of *Archaea* have been reported to have short generation time and to withstoond nutrient stress for longer durations [116]. Like for *Bacterial* species, and in the absence of sulfur, *Thermococcales*, have been reported to be involved in both acidogenesis and acetogenesis reactions. Species of this group metabolize proteins and carbohydrates into organic acids, CO_2(g)_, and H_2(g)_ [117, 118]. Other affiliates of *Thermococcales*’ are involved in homoacetogenic reactions through which they oxidize CO_(g)_ into CO_2(g)_, and catabolize pyruvate into H_2(g)_ and other products [119, 120]. We also identified *Archaeoglobales*, the affiliates of *Archaeoglobus* class which use sulfate, as the reducing agent, to oxidize lactate, long fatty acids and acetate into CO_2(g)_ [121]. Some species e.g *Geoglobus ahangari*, however, reduce Fe^3+^ compounds with H_2(g)_ [122]. The identified *Archaeoglobales* and other metal reducing species identified in our study indicate the importance of these microbes in redox balancing reactions which contribute to reactor stability.

The *Sulfolobales* and *Desulfurococcales*, all belonging to *Thermoprotei* and *Crenarchaeota* phylotypes were also identified in this study (SFig. 37a). Species of these groups metabolize carbohydrates and amino acids [123, 124] into pyruvate and CO_2(g)_. They also ferment pyruvate into lactate and CO_2(g)_. The CO_2(g)_ generated from the above reactions either enters into the CO_2(g)_ reductive pathway, the route that is catalyzed by acetogens/homoacetogens to produce CH_4(g)_ or is fixed through the 3-hydroxypropionate/4-hydroxybutyrate or dicarboxylate/4-hydroxybutyrate cycles by the identified *Thermoprotei* species to generate the energy [125]. Among other *Thermoprotei* affiliates identified in this study were *Thermoproteales* that use sulfur and its compounds as electron acceptors, to ferment carbohydrates, organic acids and alcohols [126]. In these reactions, the species consume CO_2(g)_ and H_2(g)_. Though the species have been previously reported to grow at low rate, they also ferment and oxidize amino acids and propionate, respectively [127]. Other affiliates like *T. neutrophilus* are known to assimilate acetate and succinate in the presence of CO_2(g)_ and H_2(g)_ [128]. This species use H_2(g)_ to reduce sulfur. Unlike most of the other *Archaeal* species, *Thermoproteales* use oxidative and reductive TCA routes to conserve energy. *Acidilobales* another *Thermoprotei* order identified in this study uses S^0^ and thiosulfate to metabolize protein and carbohydrate substrates [129]. The species use either EMP or the Entner-Doudoroff (E-D) [130] pathways to convert the substrates into acetate and CO_2(g)_ [131]. The affiliates of *Acidilobales* are also able to oxidize triacylglycerides and long-chain fatty acids [131] into acetate and CO_2(g)_. Substrate-level and oxidative phosphorylation are the major routes of energy conservation in *Acidilobales*; and the two mechanisms occur when S^0^ and acetyl-CoA concentrations are high. Due to S^0^ respiration in *Acidilobales* species, we postulate that their member species close the anaerobic carbon cycle via complete mineralization of organic substrates and the variations of these substrates together with other deterministic and non-deterministic (Table 2 and 3) factors contributed to the observed variation in abundance among the treatments and the high β-diversity (SFig. 37b). We also hypothesize that soil type was another factor that exerted natural selection on the *Acidilobales* through the alteration of the cow-dung’s substrates chemical composition.

The identified *Thaurmarchaeota* communities were classified into *Cenarchaeles* [132] and *Nitrosopumilales* [133] phylotypes (SFig. 38). The member species of these groups use the modified 3-hydroxypropionate/4-hydroxybutyrate pathway to fix inorganic carbon to generate energy. The communities are also able to metabolize urea and its products into, NH_3(g)_ [134, 135]. The *Nanoarchaeota* (SFig. 31a) species exploit *Crenarchaeota* mechanisms [136, 137] to metabolize amino acids and complex sugars [138, 139] while the *Korarchaeotes* are sole amino acid fermenters. The energy metabolism in the last two phyla is still unclear and further biochemical studies are needed to unravel their full metabolic activities.

### The identified *fungal* biomes

Like the *Bacterial* species described in this study, the identified *fungi* (SFig. 39a, 40a and 41a) have the ability to convert macromolecular substrates into acetate, CO_2(g)_ and H_2(g)_ [140]. Among the identified fungal species, *Ascomycota* were the richest and most abundant guild, comprising of <1% of the identified communities in the treatments. Among the *Ascomycota* species, a majority were affiliates of *Saccharomycetales* (*Saccharomycetes*, *Ascomycota*), that generally use the EMP pathway to ferment glucose and other sugars into pyruvate. One of such identified species is *Saccharomyces cerevisiae* [141] that is known to ferment carbohydrates into mixed fermentation products. Other *Saccharomycetes* species e.g *Hansenula polymorpha* yeast ferment monosaccharides, fatty acids and amino acids into acetate, ethanol, CO_2(g)_ and H_2(g)_ [142]. Furthermore, the species are also able to oxidize methanol into formaldehyde and H_2_O_2(g)_ [143]. The two products act as the sources for formate and H_2(g)_, the precursors for methanogenesis. The observed variations in abundance of the *Saccharomycetes* species may be attributed to their metabolic plasticity, deterministic factors and other environmental variables. The other *Ascomycota* affiliates belonging to *Pezazales* (*Pezizomycetes*), *Pleosporales* (*Dothideomycetes*), and *Eurotilaes* (*Eurotiomycetes*) identified in this study use mixed fermentation pathways to ferment carbohydrates [144, 145]. The metabolisms of these *Ascomycota* species have however been observed to vary within and among the phylotypes, despite the presence of a common enzyme for the pyruvate metabolism [144, 145]. For example, among the *Eurotiale*’s order; the species *Paecilomyces variotii* expresses a low metabolic capacity on plant-derived compounds compared to other *Eurotiale*’s members [146, 147]. This seems to indicate the genetic diversity of *Eurotiomycetes*’ species, seen in the variations of abundance of the species among the treatments in this study. Our data also suggests that the communities of *Eurotiomycetes* were influenced by the environmental temperature and soil type (Table 2 and 3 and SFig. 45a). Like the identified *Fusobacteria* species and other identified amino acid metabolizers, *Dothideomycetes*, another amino acid consumer phylotype, metabolize lipids and proteins into volatile organic acids [147]. On the other hand, the affiliates of *Pezizomycetes* use sulfate as an electron sink to consume amino acids [148, 149]. We attribute the community variation observed within the above groups to the composition of the cow-dung’s substrates and natural selection pressure differences among the treatments.

The *Sordariomycetes*’ affiliates identified in this study (SFig. 44a) produce enzymes that express an assortment of degradative mechanisms and other fermetative roles. The *Hypocreales*’ affiliates are such an example that produces powerful hydrolytic enzymes [150] that catabolize complex biomass in their vicinity. *Fusarium* sp. a member of *Hypocreales*, metabolizes NH_3(g)_, lipids and proteins [151, 152] into the respective smaller molecules. The species has been found to use NAD(P)H to reduce NO_3_^−^_(aq)_ into NH_4_^+^_(aq)_. The production of NH_4_^+^_(aq)_ by the *Hypocreales* is normally coupled to the substrate-level phosphorylation and ethanol oxidation; the processes that produce acetate [151, 152]. The two mechanisms ameliorate the NH_4_^+^_(aq)_ toxicity to the micro-organisms. Other *Hypocreales* species such as *Hirsutella thompsonii* confer wide metabolic activities that involve chitinases, lipases, proteases and carbohydrate degrading enzymes [153]. *Monilia sp*. [154] and *Neurospora crassa* [155] were among the identified *Sordariomycetes* and they convert cellulose into ethanol. *Magnaporthales*’ such as *Pyricularia penniseti* convert isoamylalcohol into isovaleraldehyde while other *Magnaporthales*’ species convert fumaryl-acetoacetate into fumarate and acetoacetate [156, 157]. Other roles played by the *Sordariomycete*’s species include the metabolism of aromatic compounds through which, phenolics and aryl alcohols/aldehydes are converted into acetate, formate and ethanol [158, 159]. Our data suggests that the *Sordariomycete*’s community’s abundances were influenced by the cow-dug’s substrates and the environmental variables (Table 2 and 3) of the reactors.

In contrast to the *Ascomycota* species, affiliates of *Basidiomycota* establish susceptible hyphal networks [160] that may have been constrained by the reactor pertubartions in our study. This may explain why the *Ascomycota* species dominated over the *Basidiomycota*’s species in this study. Nonetheless, like *Ascomycota* species, the affiliates of *Malasseziales*, *Ustilaginales*, *Agaricules* and *Tremellales* species, all belonging to *Basidiomycota*, are able to metabolize carbohydrates [145, 161], urea [161] and NO_3_^−^_(aq)_ [161] in the systems. Other members like *Polyporales* metabolize woody substrates and their respective products [162]. All these functions clearly indicate that *Basidiomycota’s* species played diverse metabolic roles in our AD systems in this study. Interestingly, we also identified *Chytridiomycota* affiliates in reactor 9 and 10. These affiliates are capable of catabolizing pollen, chitin, keratin, and cellulose [163, 164], which may be indicative of the presence of these substrates in the two treatments. The detected *Spizellomycetale*s, *Cladochytriale*s and *Rhizophydiale*s species in the two treatments, further revealed the presence of highly decayed plant tissue with high cellulose content. The identified *Monoblepharidomycetes* members are close relatives of *Cladochytriales* and *Spizellomycetales* [165] species and were detected only in reactor 9 and we postulate that these groups perform similar roles in the AD systems.

Generally, *Chytridiomycota* species are carbohydrate metabolizers as previously revealed by the presence of glucose transporters and invertase activity in their systems [163, 164]. The species use electron bifurcation mechanisms to conserve their energy and the mechanisms occur in the hydrogenosome [166]. The *Blastocladiales*, the close relatives of *Chytridiomycotes* and the affiliates of *Blastocladomycota*, class *Blastocladomycetes* were only detected in reactor 9 in our study. Their member species metabolize cellulose into acetate, CO_2(g)_ and H_2(g)_. Interestingly, *Blastocladiales* have previously been reported to use the Malate fermentation pathway to generate and conserve their energy [167]. *Mortierellales* formerly classified under *Mucoromycotina* [168] and latter assigned to *Mortierellomycotina* phylum [169] are converters of organic compounds into poly-unsaturated fatty acids [170] while their relatives *Entomophthorales* and *Mucorales* metabolize decaying organic matter. However, the affiliates of the three orders were only detected in reactor 9, 10 and 11, respectively.

Like similar studies [171, 172], our data revealed that fungi co-occurred with the species of *Bacteria* and methanogens. In these syntrophy, we postulate that the fungal species catalyzed inter-species H_2(g)_ transfer and re-generated oxidized reducing equivalents; NAD^+^ and NADP^+^ in the systems. All these reactions increased metabolism of the dry matter [173] in the treatments. We also suggest that the fungal syntrophy is more complex than simple cross-feeding mechanisms, because the species shift more oxidized products into a more reduced form. Through these mechanisms, the precursors, lactate and ethanol are converted into acetate and formate depending on the prevailing conditions. The involved interactions between fungi and other domain’s species have been determined to be very crucial to an extent of influencing the isolation and culturing of some fungal species [173, 174]. The findings indicate why fungal species should be jointly considered with other domains’ species in the context of biogas production.

### The microbiomes’ α- and β-diversities and biogas production

To establish whether environmental conditions and perturbations influenced the microbial populations, we compared the collected phenotypic traits with the genotypic dataset. The phenotypic datasets revealed an insight on the cause of the observed communities’ β-diversities and biogas production variation. We attempted to use the principles of the four ecological mechanisms that include selection, drift, speciation, and dispersal to better understand the cause of the observed variance in microbial composition among the treatments and thus the cause of the biogas production variability among the sites. While selection is deterministic and ecological drift is stochastic, dispersal and speciation may have contributed to the two ecological parameters. One of the challenges of examining communities’ in any given niche is the difficulty of estimating real dispersal among the sites. Nonetheless, our treatments were fluidic, naturally encouraging rapid microbial dispersion. Hence, species dispersal was not the major limiting factor that influenced our local communities. Similarly, we also excluded speciation based on our sampling process and experimental design. Therefore, we suggest that selection and drift were the likely two main factors that contributed to variation of community abundances among the treatments and the observed high β-diversities in our study. The two factors were most likely influenced by the substrate inputs (Table 3) and disturbances from environmental variables (Table 2 and 3). Environmental conditions directly affect the chemical composition of the substrate fed to the cattle breeds. As a result, their wastes, the cowdung is fed to the reactor which ultimately influences the microbial composition and biogas productivity. Temperature, agroecological zones and soil types were among the environmental factors that were observed to drive the specific communities’ β-diversities and their relative abundances among the treatments. All these could have contributed to the observed variation of biogas production among the studied treatments. We postulate that environmental temperatures combined with other deterministic factors such as the species metabolic traits, largely contributed to the lack of identification of the *Chytridiomycota*, *Blastocladomycota* and other rare fungal species in the majority of the treatments. The findings increased the number of the total species in the few treatments, where the rare species were identified as opposed to where they were absent. Nonetheless, we never observed significant differences in reactor performances between the two categories of the treatments. Consequently, we linked the observations with the low abundances of the rare *fungal* species identified in the few respective reactors, which implies the importance of the species abundance in biogas production. The composition of the cow-dung’s substrates fed to these treatments could largely be influenced by cattle breeds and several environmental factors including agroecological zones, soil type, rainfall, environmental temperatures to mention but a few, and its chemical compositions could have stimulated the growth of specific species, favouring colonization and speciation in the respective treatments. Similarly the same phenomenon could have happened to other microbial communities. In this case, the available resources could have reduced competition among the species, weakening the niche selection abilities. Generally these ecological factors normally strengthen the ecological drift and prority effects [175] that result to unpredictable site to site variation of microbial abundances and high β-diversities, even under similar environmental conditions. We used the same arguments to explain our observations on the variations of the relative abundances among treatments and high β-diversities, particularly when the conditions were similar (Table 2 and 3). We also extended the same argument to the biogas production capacities among the treatments.

Besides the cow-dung’s substrate effects, we hypothesize that the intensity and duration of the disturbances [176], led to high rate of species mortality, low growth rates and species extinction. These kinds of disturbances caused decrease of stochasticity and to account for their effects on our findings, we adopted the concepts of neutral theory [176]. Generally their associated caveats reduce ecological drifts that result to low competition abilities among the low abundance species. Due to these facts, we postulate that the persistence and intensities of these disturbances in our specific treatments eliminated weak and less adopted microbial populations. One of such identified case was revealed in reactor 5 in which the *Tenericutes* affiliates were absolutely extinct. Similarly the absence of *Mortierellales, Entomophthorales*, *Mucorales* to mention but a few, in other treatments (SFig. 51) was an indication of unfavorable disturbances that could have originated from either cow-dung’s substrates or other reactor perturbations. All these may have led to more predictable site-to-site variations and we used these concepts to explain the observed low β-diversity among the treatments (e.g in reactor 2 and 10, reactor 3 and 6 etc). However, to understand better the links between the environmental conditions and the microbial communities in these AD systems, we suggest the utilization of the metacommunity theory that considers the world as a collection of patches that are connected to form a metacommunity through organisms’ dispersal. The model depends on the specific species traits, the variance of the patches in the environmental conditions as well as frequency and extent of species dispersal in the systems. Other ecological models lack these features. Therefore, the model can be used to ask and test whether dispersal, diversification, environmental selection and ecological drift were the vital elements in the biogas production process. In this case, a detailed understanding of the contribution of these factors on specific species abundances and community assemblage can be used to tailor the reactor management from an ecological point of view.

### Conclusions and Future prospects

Our results provide an insight into the biogas microbiomes composition, richness, abundances and variations within and among the taxa and the twelve reactor treatments. Interestingly, microbiome structure and richness were similar among the twelve treatments, except for the affiliates of *Rhizophydiales*, *Cladochytriales*, *Monoblepharidales*, *Blastocladiales* and *Entomophthorales*, which were detected only in reactor 9. Contrasting the findings were the affiliates of *Mortierellales* and *Mucorales* that were identified in reactor 10 and reactor 11, respectively, while the *Spizellomycetales* species were detected in both reactor 9 and 10. We also revealed that the affiliates of *Tenericutes*, were the only *Bacterial* guild that was absent in reactor 5. The Cow-dung’s substrates and metabolic capabilities of these species were the main elements that were considered to limit their dispersal to other treatments. Nonetheless, the identified phylotypes richness within and among the phyla were found to vary, except for the phyla, that comprised a single phylotype. All the identified species abundances among the treatments and the community’s β-diversities in the majority of the treatments significantly varied. All these contributed to our conclusion that environmental and spatial variables largely influenced biogas microbiome populations and thus the amount of biogas production. The discoveries highlight the importance of identifying the involved natural processes that give rise to such identified variations. Based on the results obtained in this study, we suggest further studies on species responses on the natural conditions for optimal biogas production and microbial community managements. We also suggest the application of metacommunity theory among other ecological theories in developing management strategies for the AD systems.

## Materials and Methods

### Ethics and experimental design

The sampled treatment sites (reactors) were installed in the premises of privately owned land (Table 2) and permission to sample was granted by the owners. Randomized complete block design was utilized as previously described [177] to assign the twelve cowdung digesting treatments to the four blocks, which operated at mesophilic conditions. Organic loading rate and the hydraulic retention time were largely influenced by the the treatment size and the farmers’ routine activities. Sampling was conducted following the previously described procedure of Boaro *et al*. [178]. Geographic co-ordinates (Table 2) and agroecological zones (Table 3) were generated by Geographic Positioning System (etrex, Summit^*HC*^, Garmin) and ArcGIS^®^ software (ver10.1) [179], respectively. A pre-tested questionnaire was utilized according to Oksenberg *et al*. [180] protocol to collect the phenotypic dataset. Small scale farming and zero grazing of Freshian breeds were the main activities among the sampled farmers. However, the owners of reactor 3 and 11 (Meru and Kiambu blocks, respectively), reared Jersey breeds of cattle. Samplings were conducted during the warm and cool climatic conditions.

### Sample preparation and Sequencing

The double stranded deoxyribonucleic acids (dsDNA) was extracted from the duplicate sludge samples using ZR soil microbe DNA kit (Zymo Research, CA., USA), according to manufacturers instructions. The quantity and quality of the extracted dsDNA were determined as previously described by Campanaro *et al*. [181]. The dsDNA was normalized and library preparation performed using Nextera™ DNA sample prep kit (Illumina, San Diego, CA, USA), according to manufacturers instructions. Library normalization was conducted using Illumina TruSeq DNA PCR-free sample preparation kit buffer (Illumina, USA). To prepare for cluster generation and sequencing, equal volumes of normalized libraries were pooled, diluted in hybridization buffer and denatured, according to Nextera™ protocol. *De novo* deep sequencing was conducted by an Illumina MiSeq system as previously described by Caporaso *et al*. [182] but adopting a v3 kit chemistry with 300 cycles (2×150) as previously described [181].

### Bioinformatics and Statistical analysis

All paired-end reads in FASTQ format were pre-processed with FastQC software (ver0.11.5) following the procedure described by Zhou *et al*. [183]. Trimmomatic software was utilized to trim and filter the adaptor according to [184]. *De novo* assembly of the paired-end contigs were peformed by MetaSpade software (ver3.10) as described previously [185]. To re-trim low-quality regions of the scaffolds, we utilized SolexaQA as described by Cox *et al*. [186] while DRISEE, metagenomic quality assessment software was used to remove duplicated scaffolds according to Keegan *et al*. [187]. We screened near-exact scaffolds that matched human, mouse and cow genomes following Langmead *et al*. procedure that used a sort read aligner, the Bowtie [188]. M5nr [189], a non-redundant tool, FragGeneScan [190] and sBLAT [191] tools were utilized to compute datasets against 16 reference biological databases as previously described by Wilke *et al.* [189]. Two ENVO ontologies: large lake and terrestrial biomes were utilized according to Field *et al*. [192] and Glass *et al*. [193]. Alignment similarity cutoff were set at single read abundance; ≥60% identity, 15 amino acid residues and maximum e-value of 1e^−5^. The annotated datasets were archived in CloVR [194]. Prior to statistical analysis, the rarefaction curves were reconstructed, datasets was log_2_ (x+1) transformed and normalized using an R package, DESeq software [195]. PCoA based on euclidean model [196] was set at *P*≤0.05, and conducted against the twelve treatment samples. Results were visualized by MG-RAST pipeline (ver4.01) [197] and Krona radial space filling display [198]. Optimal scripts and restful application programming interface were utilized in our analysis.

## Acknowledgement

We appreciate Vincent Njunge, Collin Mutai, Tina Kyalo, Joyce Njoki Njuguna, Eunice Machuka, Muga Fredrick Nganga and Philis Emilda Ochieng of BecA-ILRI hub and Donald Otieno and Linnet Gohole of University of Eldoret for their technical surport.

## Authors Contribution

**Conceived idea**: MSM

**Sourced funds:** MSM

**Designed experiments:** MSM, OSO

**Performed experiments:** MSM

**Performed Sequencing:** MSM, MLW

**Analyzed the data:** MSM

**Drafted manuscrispt:** MSM

**Reviewed the manuscript:** MSM, OSO, PR, WFN, OJM, NPM

## Supplementary Material

**STable 1:** The quality control statistics of the obtained scaffolds for the twelve treatments

**SFig. 1:** Bar charts showing unfiltered and filtered sequencing reads (a) and the known and unknown protein genes (b) in our samples.

**SFig. 2**: Stacked barchat showing some (21 phyla) Bacteria domain phyla, relative abundances (a) and the PCoA plot (phylum level), based on the Euclidean model (b).

**SFig. 3:** The PCoA plot showing the relative abundances variation at the class (a) and order level (b).

**SFig. 4:** The stacked barchart revealing six classes of *Proteobacteria* communities, relative abundances (a) and their PCoA plots based on the Euclidean model (b).

**SFig. 5:** The PCoA plot based on the Euclidean model for all the identified *Proteobacteria* orders.

**SFig. 6:** Stacked barchat showing seven *δ-Proteobacteria* orders, relative abundances (a) and their PCoA plot revealing nucleotide composition variations, based on the Euclidean model (b).

**SFig.7:** Stacked barchart showing fourteen *γ-Proteobacteria* orders (a) and their PCoA plot based on the Eucleaden model (b).

**SFig. 8:** Stacked barchat showing seven *β-proteobacteria* orders, relative abundances (a) and their PCoA plot revealing nucleotide composition (dis)similarities among the reactors (b).

**SFig. 9:** Stacked barchat showing the seven *α-Proteobacteria* orders, relative abundances (a) and their PCoA plots, revealing the nucleotide composition variation among the reactors based on the Euclidean model (b).

**SFig. 10:** Stacked barchat showing two *ε-Proteobacteria* orders, relative abundances (a) and their PCoA plot based on Euclidean model (b).

**SFig. 11:** The stacked barchat revealing four *Firmicutes* classes, the relative abundances (a) and their PCoA plots, revealing variance among the twelve reactors based on the Euclidean model (b).

**SFig. 12:** Stacked barchat showing eight *Firmicute’*s orders, relative abundances (a) and their PCoA plot based on the Euclidean model (b).

**SFig. 13:** Stacked barchat showing the four *Clostridia* orders (a) and their PCoA plots, revealing nucleotide composition variations among the twelve reactors based on the Euclidean model (b).

**SFig. 14:** Stacked barchat showing two *Bacilli* orders, relative abundances (a) and their PCoA plot based on the Euclidean model (b).

**SFig. 15:** The stacked barchat revealing four *Bacteroidete*s classes, relative abundances (a) and their PCoA plots revealing nucleotide composition variation, based on the Euclidean model (b).

**SFig. 16:** Stacked barchat showing the four *Bacteroidete’*s orders, the relative abundances (a) and their PCoA plot based on the Euclidean model (b).

**SFig. 17:** Stacked barchat showing six *Actinobacteria* orders, the relative abundance (a) and their PCoA plot based on the Euclidean model (b).

**SFig. 18:** The stacked barchat showing four *Chloroflexi* classes, relative abundances (a) and PCoA plot revealing variance among the twelve reactors, based on the Euclidean model (b).

**SFig. 19:** Stacked barchat showing five *Chloroflexi* orders, relative abundances (a) and their PCoA plot based on the Euclidean model (b).

**SFig. 20:** Stacked barchat showing two *Chloroflexi* class orders, relative abundances (a) and their PCoA plot based on their Euclidean model (b).

**SFig. 21:** Stacked barchat showing the two *Thermomicrobia* orders, relative abundances (a) and their PCoA plot based on the Euclidean model (b).

**SFig. 22:** Stacked barchat showing *Cyanobacteria* class, relative abundances (a) and their PCoA plot based on the Euclidean model (b).

**SFig. 23:** Stacked barchat showing the five *Cyanobacteria* orders, relative abundances (a) and their PCoA plot based on the Euclidean model (b).

**SFig. 24:** Stacked barchat showing the four affiliates of the Unclassified *Cyanobacteria* nucleotide reads, relative abundances (a) and their PCoA plots based on the Euclidean model (b).

**SFig. 25:** The stacked barchat showing the two *Acidobacteria* classes, relative abundances (a) and their PCoA plots for their nucleotide composition based on the Euclidean model (b).

**SFig. 26:** The stacked barchat showing the two *Acidobacteria* orders, relative abundances (a) and their PCoA plots for their nucleotide composition based on Euclidean model (b).

**SFig. 27:** Stacked barchat showing two *Deinococcus-Thermus* orders, the relative abundances (a) and the PCoA plots based on the Euclidean model (b).

**SFig. 28:** Stacked barchat showing *Verrucomicrobia* classes, relative abundances (a) and their PCoA plot based on the Euclidean model (b).

**SFig. 29:** Stacked barchat showing three *Verrucomicrobia* orders, relative abundances (a) and their PCoA plot based Euclidean model (b).

**SFig. 30:** Stacked barchat showing three *Terneicutes* orders, relative abundances (a) and their PCoA plot based on the Euclidean model (b).

**SFig. 31:** The stacked barchart showing the four *Archaea* phyla, relative abundances (a) and their PCoA plot based on the Euclidean model (b).

**SFig. 32:** Stacked barchat showing nine *Archaea* classes, and the affiliates of the unclassified reads, relative abundances (a) and their PCoA plot based on the Euclidean model (b).

**SFig. 33:** Stacked barchat showing sixteen *Archaea* orders, relative abundances (a) and their PCoA plot based on the Euclidean model (b).

**SFig. 34:** The stacked barchat showing the eight *Eurychaeota* classes, the relative abundances (a) and their PCoA plots revealing dissimilarities of the nucleotide composition based on the Euclidean model (b).

**SFig. 35:** Stacked barchat showing ten *Euryarchaeota* orders, relative abundances (a) and their PCoA plot based Euclidean model (b).

**SFig. 36:** The stacked barchat showing three *Methanomicrobia* orders, the relative abundances (a) and their PCoA plot based on the Euclidean model (b).

**SFig. 37:** Stacked barchat showing four *Thermoprotei* orders, relative abundances and their PCoA plot based on the Euclidean model (b).

**SFig. 38:** Stacked barchat showing two *Thaurmarchaeota* order, the relative abundances (a) and their PCoA plot based on the Euclidean model (b).

**SFig. 39:** Stacked barchat showing five fungal phyla, the relative abundances (a) and their PCoA plots based on Euclidean model (b).

**SFig. 40:** Stacked barchat showing thirteen fungal classes, relative abundances (a) and their PCoA plot based on the Euclidean model (b).

**SFig. 41:** Stacked barchat showing 23 fungal orders, relative abundances (a) and their PCoA plot based on the Euclidean model (b).

**SFig. 42:** The stacked barchat showing the five *Ascomycota* classes, relative abaundances (a) and their PCoA plots based on the Euclidean model (b).

**SFig. 43:** The stacked barchat revealing eleven *Ascomycota* orders, relative abundances (a) and their PCoA plot based on the Euclidean model (b).

**SFig. 44:** Stacked barchat showing three *Sordariomycetes* orders, relative abundances (a) and their PCoA plot based on the Euclidean model (b).

**SFig. 45:** Stacked barchat showing the two *Eurotiomycetes* orders, the relative abundances (a) and their PCoA plot based on the Euclidean model (b).

**SFig. 46:** Stacked barchat showing four *Basidiomycota* classes, the relative abundances (a) and their PCoA plot based on the Euclidean model (b).

**SFig. 47:** Stacked barchat showing four *Basidiomycota* orders, relative abundances (a) and their PCoA plot based on the Euclidean model (b).

**SFig. 48:** Stacked barchart showing the proportion of the *Agaricomycetes* order, relative abundances (a) and their PCoA plot based on the Euclidean model (b).

**SFig. 49 :** Stacked barchat showing the two *Chytridiomycota* classes, and their relative abundances in the two eactors.

**SFig. 50:** Stacked barchat showing the four *Chytridiomycota* orders, and their relative abundances in the two reactors.

**SFig. 51:** Stacked barchat showing the three orders affiliated to unclassified fungal nucleotide reads.

**SFig. 52:** The PCoA analysis revealing β-diversity of the twelve reactors at the three taxa level: a) Domain, b) Phylum, and c) Class level.

