## Supplementary Fig. 1 for "Metagenomics survey unravels diversities of biogas’ microbiomes with potential to enhance its’ productivity in Kenya"

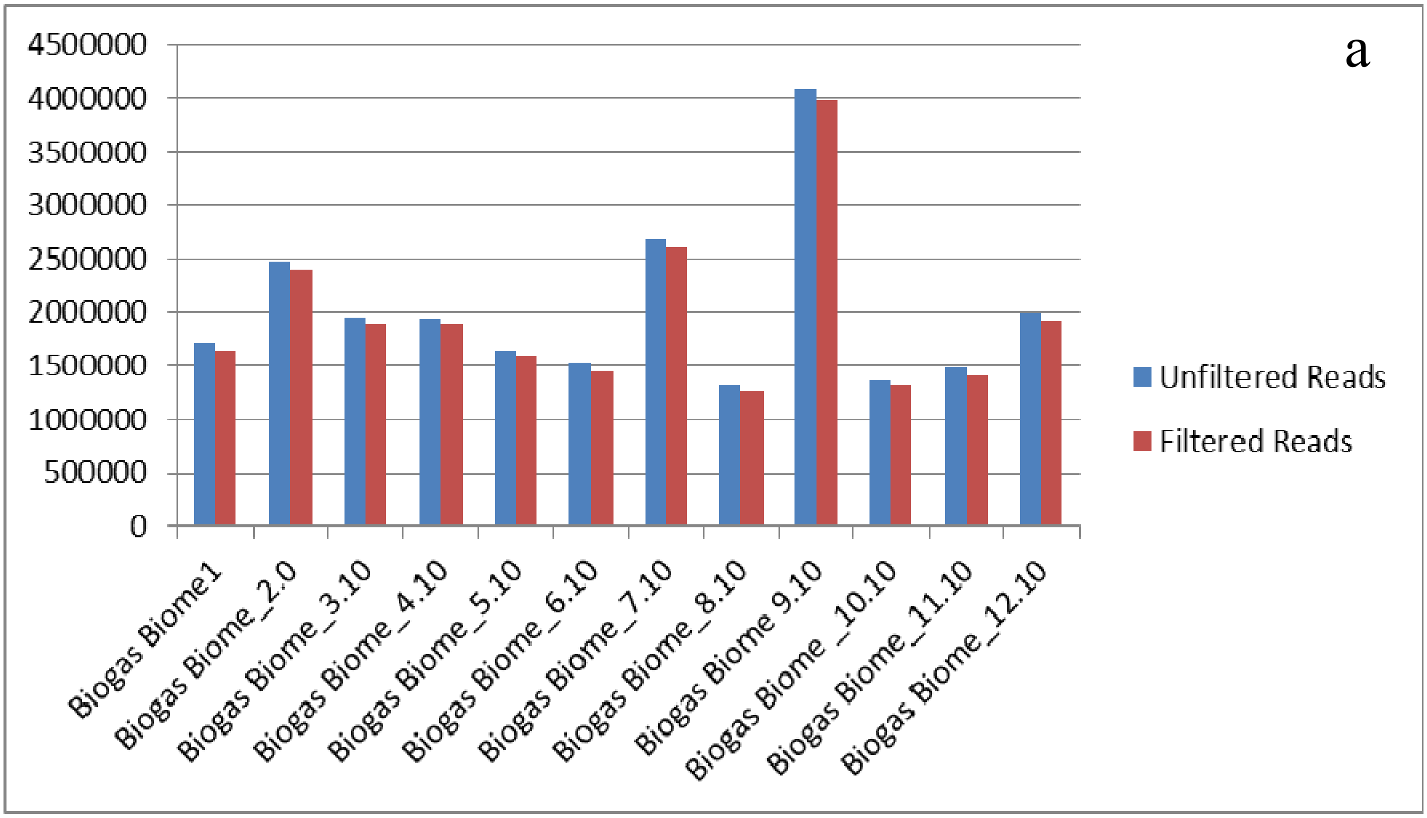


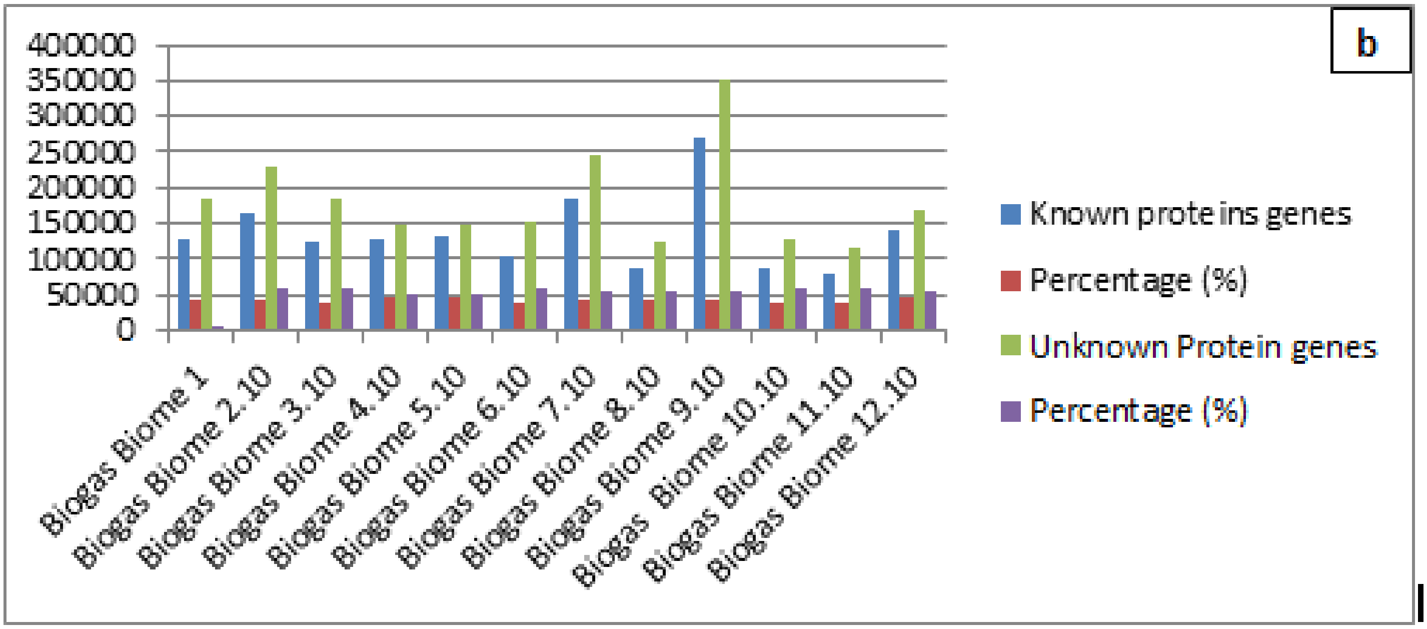


SFig 1: Bar charts showing a). Unfiltered and filtered sequencing reads and b). The known and unknown protein genes in our samples. More than 53.07% of the filtered nucleotide reads in our samples contained unknown proteins
