## Supplementary Fig,2 for "Metagenomics survey unravels diversities of biogas’ microbiomes with potential to enhance its’ productivity in Kenya"

a

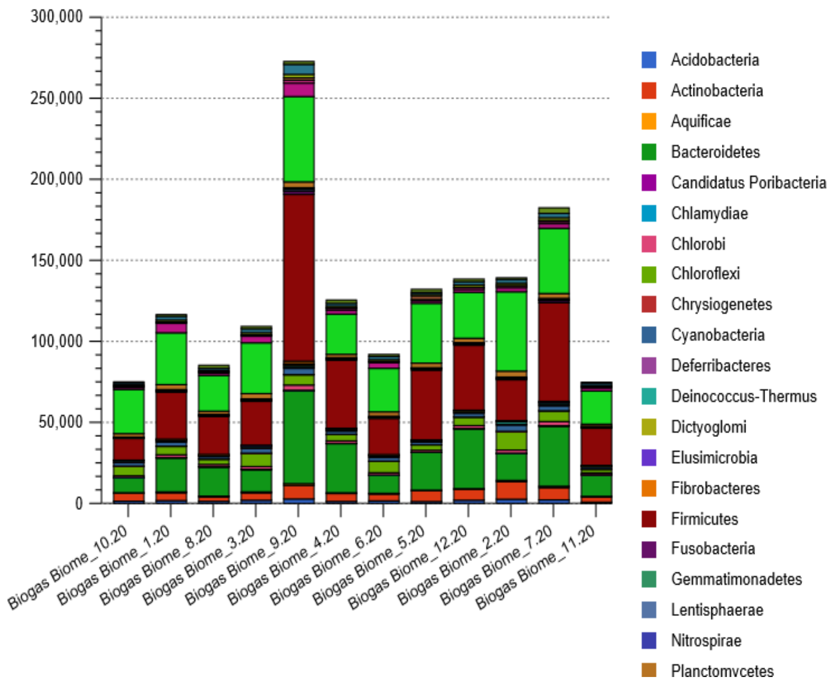


b

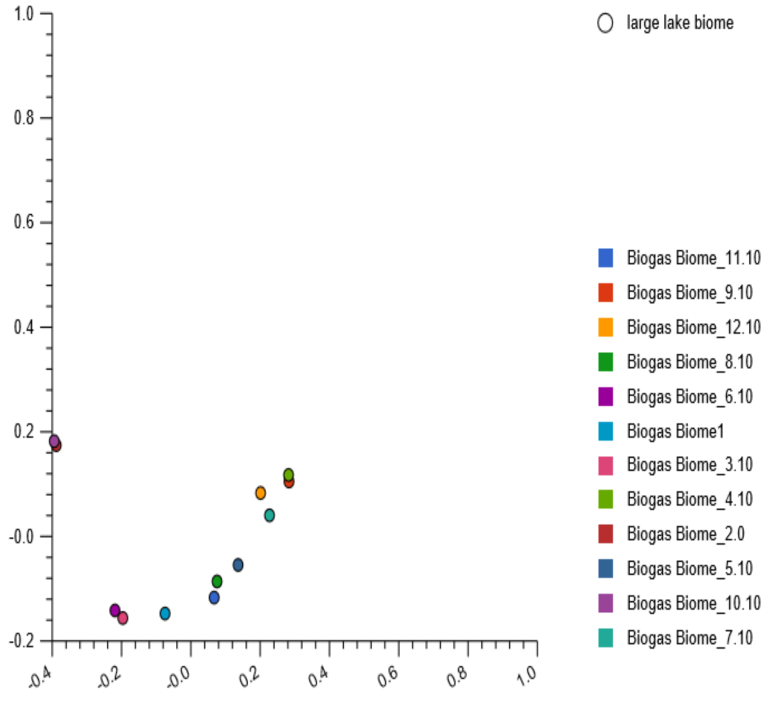


**SFig. 2**: Stacked barchat (a) showing some (21 phyla) bacteria domain phyla, relative abundances and the PCoA plot (phylum level) (b), based on the Euclidean model. The nucleotide composition in reactor 2 and 10 and those in reactor 4 and 9 partially clustered in
