## Supplementary Fig. 3 for "Metagenomics survey unravels diversities of biogas’ microbiomes with potential to enhance its’ productivity in Kenya"

a

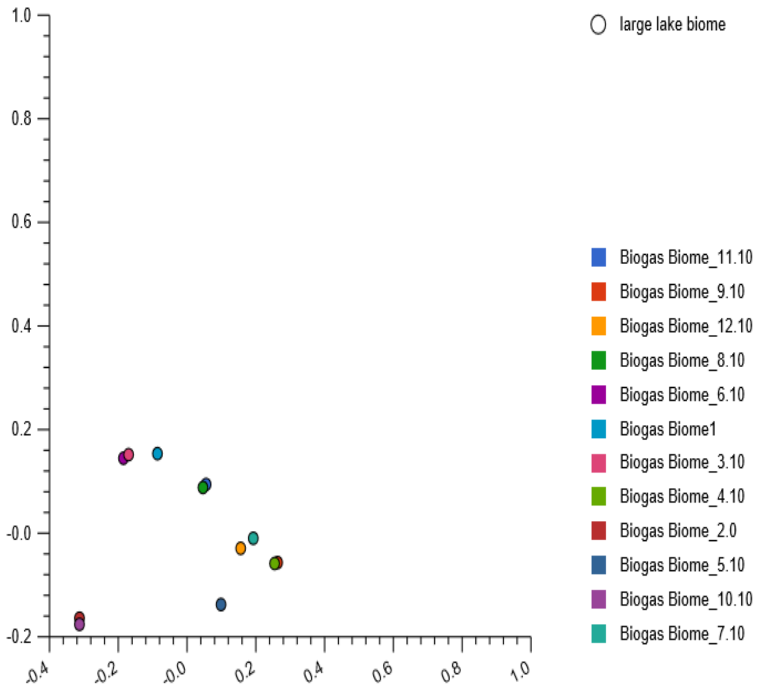


b

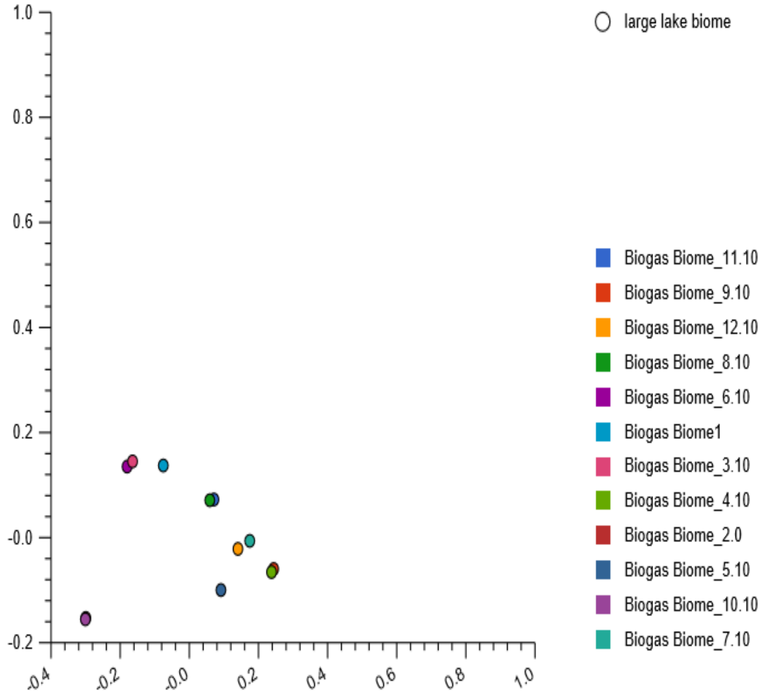


**SFig. 3:** The PCoA plot showing the (a) the nucleotide composition variation at the class and (b) the nucleotide composition variation at order level. At the class and order level, the nucleotide composition in reactor 2 and 10, located lower left quadrant, reactor 3 and 6, upper left quadrant, and reactor 8 and 11, upper right quadrant and those in reactor 4 and 9, lower right quandrant of the plot partially clustered.
