## Supplementary Fig. 4 for "Metagenomics survey unravels diversities of biogas’ microbiomes with potential to enhance its’ productivity in Kenya"

a

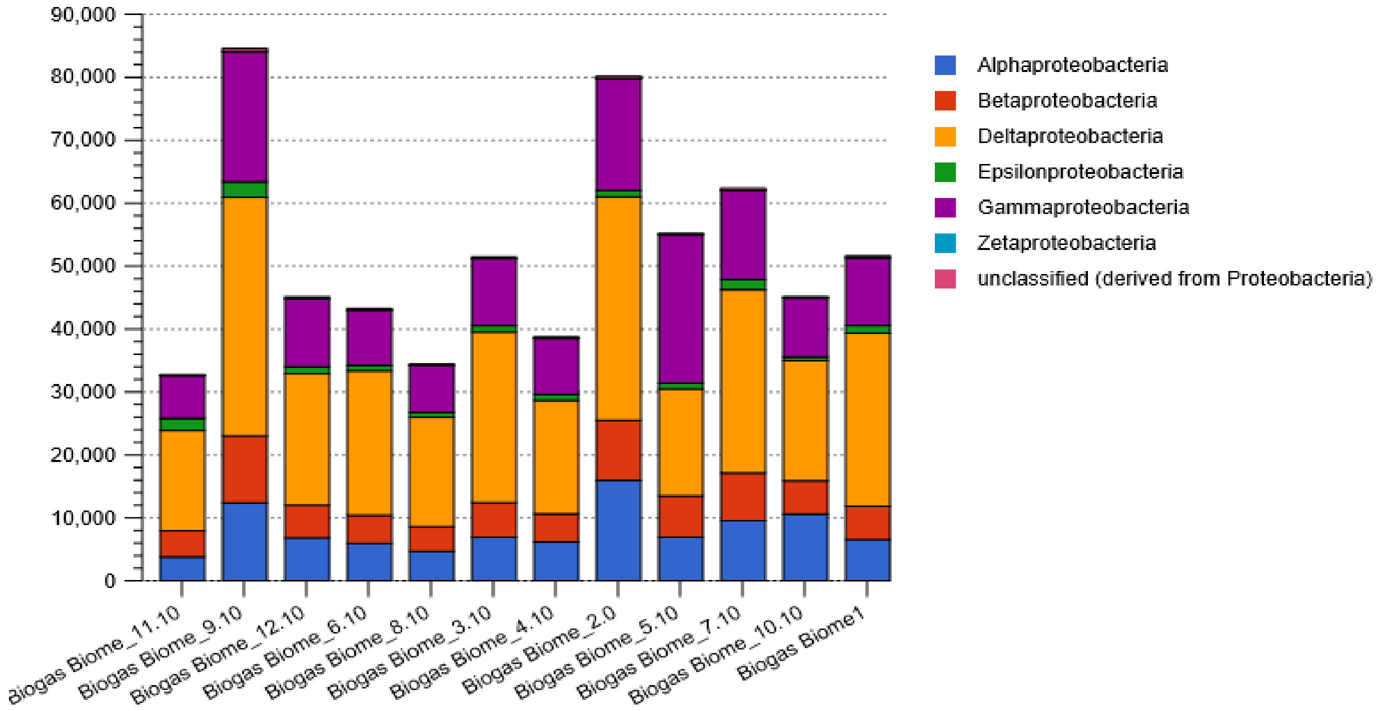


b

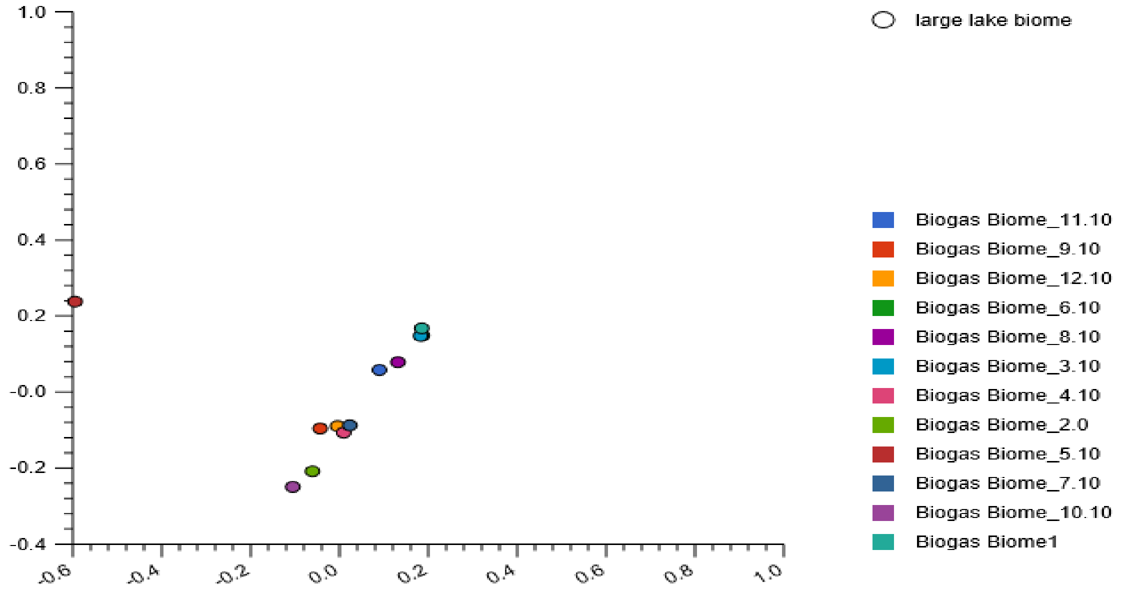


**SFig. 4:** The stacked barchart revealing six Proteobacteria classes, the proportions of their relative abundances (a) and their PCoA plots (b) based on the Euclidean model. The nucleotide composition for reactor 1, 3 and 6 clustered on the upper right quadrant of the plot. Similarly, the nucleotide composition detected in reactor 4, 7 and 12 partially clustered while those in reactor 5
