## Supplementary material for "Metagenomics survey unravels diversities of biogas’ microbiomes with potential to enhance its’ productivity in Kenya": Supplemenatry Fig. 5

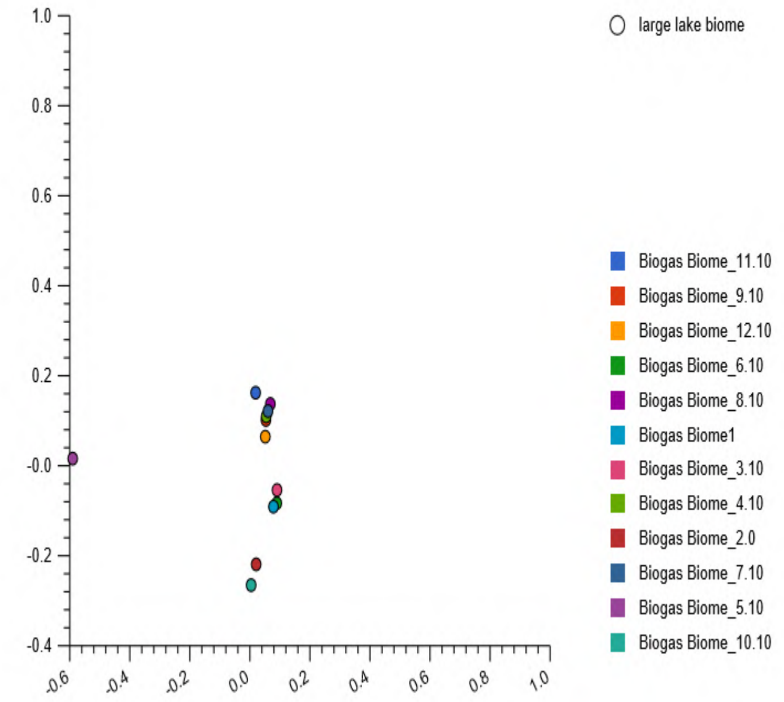


**SFig. 5**The PCoA plot based on the Euclidean model for *Proteobacteria* order. The PCoA plot indicated partial similarities of the nucleotide composition in half of the studied reactors, including reactor 1, 3 and 6, upper right quadrant; reactor 4, 7 and 12, lower right quadrant of the plot at the class level. Similar observations were made at the order level, except in a few reactors (reactor 4, 7, 8 and 9, that formed cluster.
