## Supplementary Fig. 6 for "Metagenomics survey unravels diversities of biogas’ microbiomes with potential to enhance its’ productivity in Kenya"

a

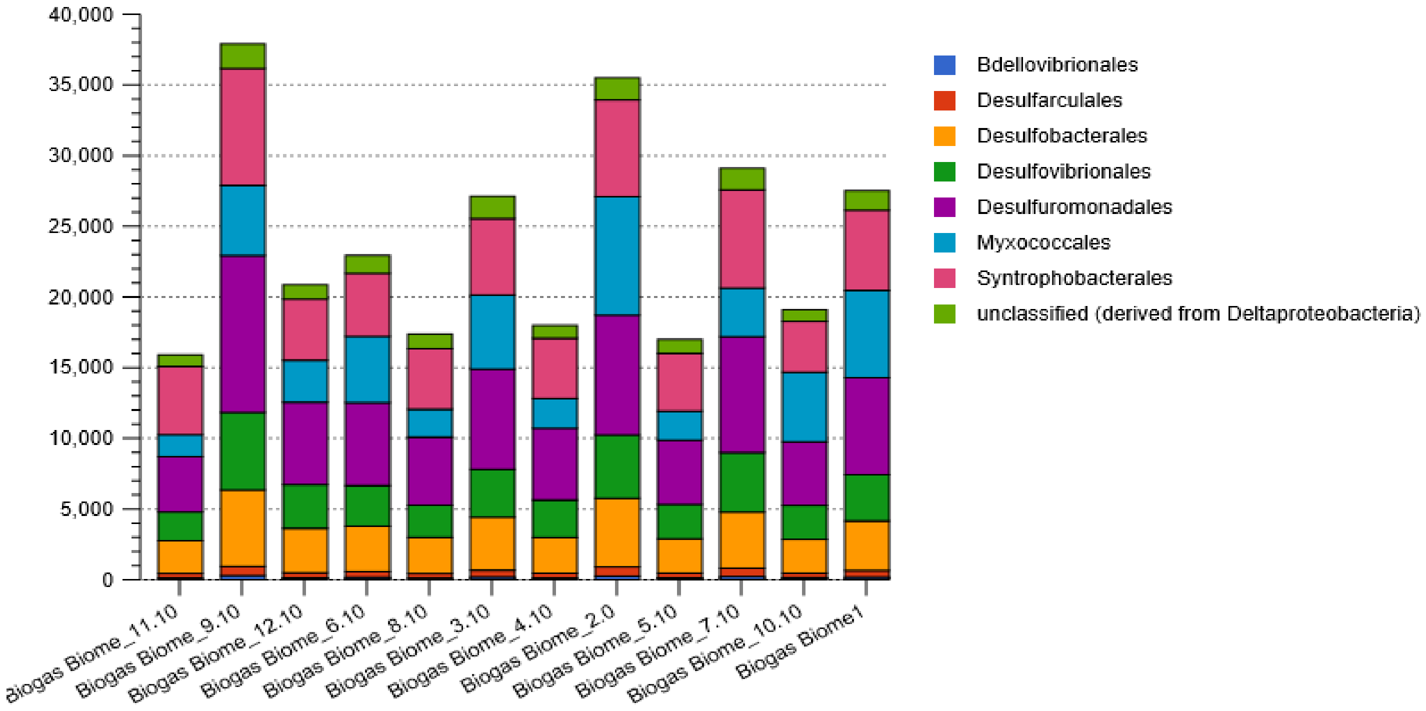


b

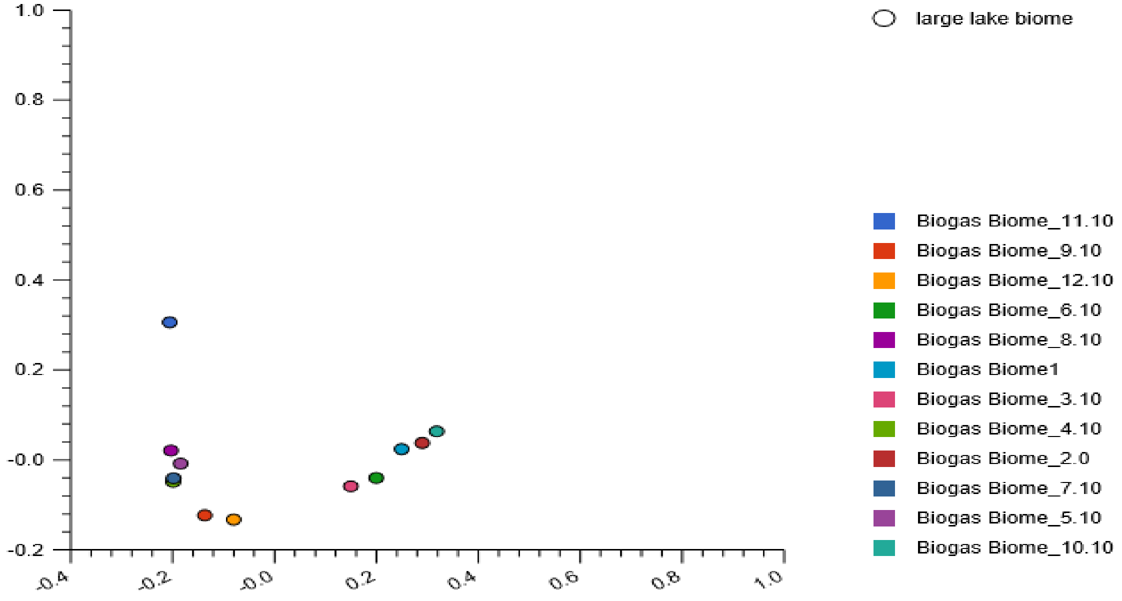


**SFig. 6:** Stacked barchat showing seven δ-Proteobacteria orders, relative abundances (a) and their PCoA plot revealing nucleotide composition variations, based on the Euclidean model. The plot revealed dissimilar nucleotide composition among the reactors except those in reactor 4 and 7 that clustered in the lower left quadrant of the plot.
