## Supplementary Fig. 7 for "Metagenomics survey unravels diversities of biogas’ microbiomes with potential to enhance its’ productivity in Kenya"

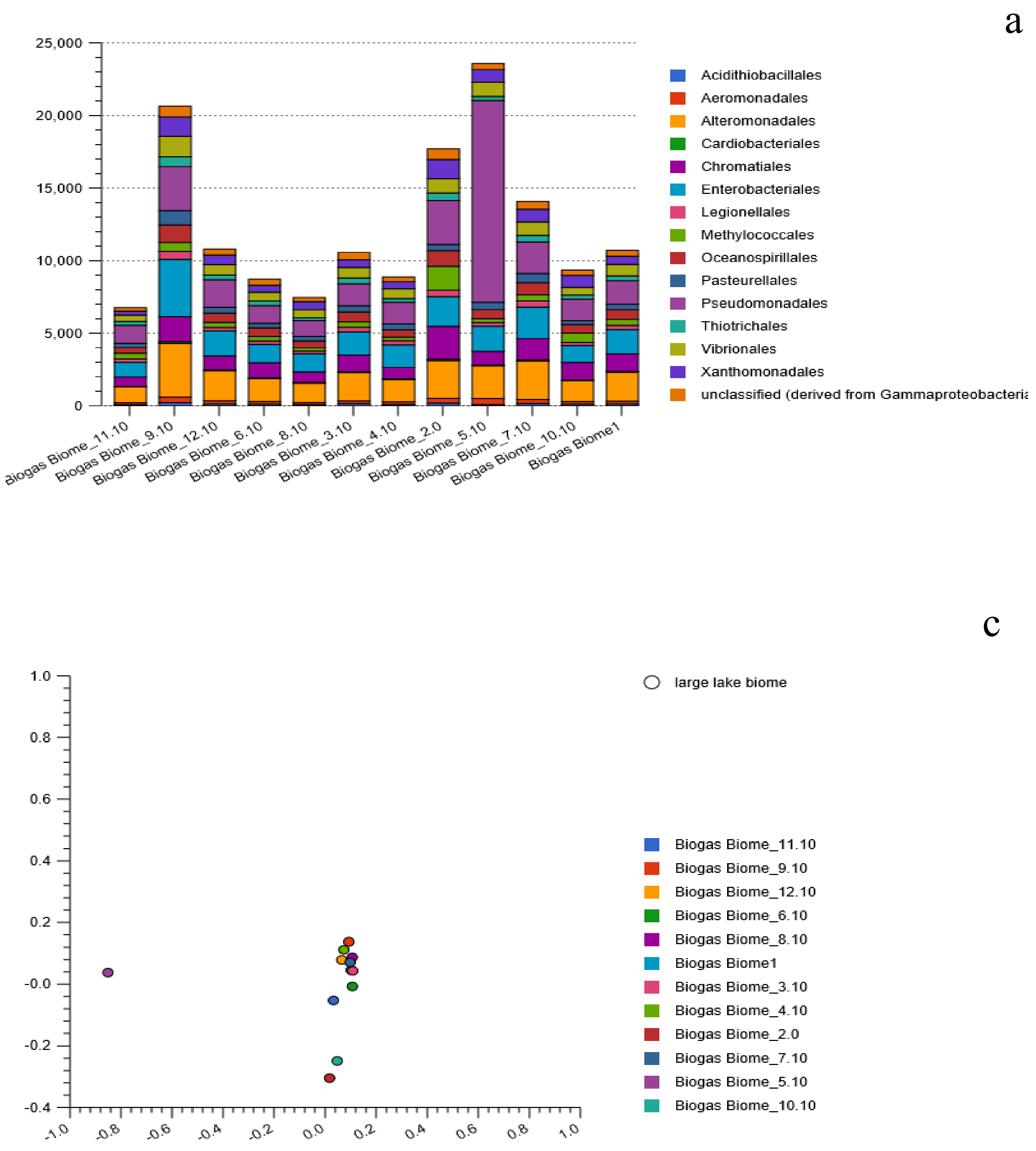


**SFig. 7:** Stacked barchart showing fourteen γ-Proteobacteria orders (a) and their PCoA plot based on the Eucleaden model. The plot revealed that the nucleotide composition in reactor 1 and 3 clustered in close proximity with reactor 7 that partially clustered with reactor 8 and 12. The nucleotide composition in reactor 4 and 9 were in close proximity. All the clustered reactors were located in the upper right quadrant of the plot. However, the nucleotide composition for reactor 5 were singly in the upper left quadrant of the plot.
