## Supplementary Fig. 8 for "Metagenomics survey unravels diversities of biogas’ microbiomes with potential to enhance its’ productivity in Kenya"

a

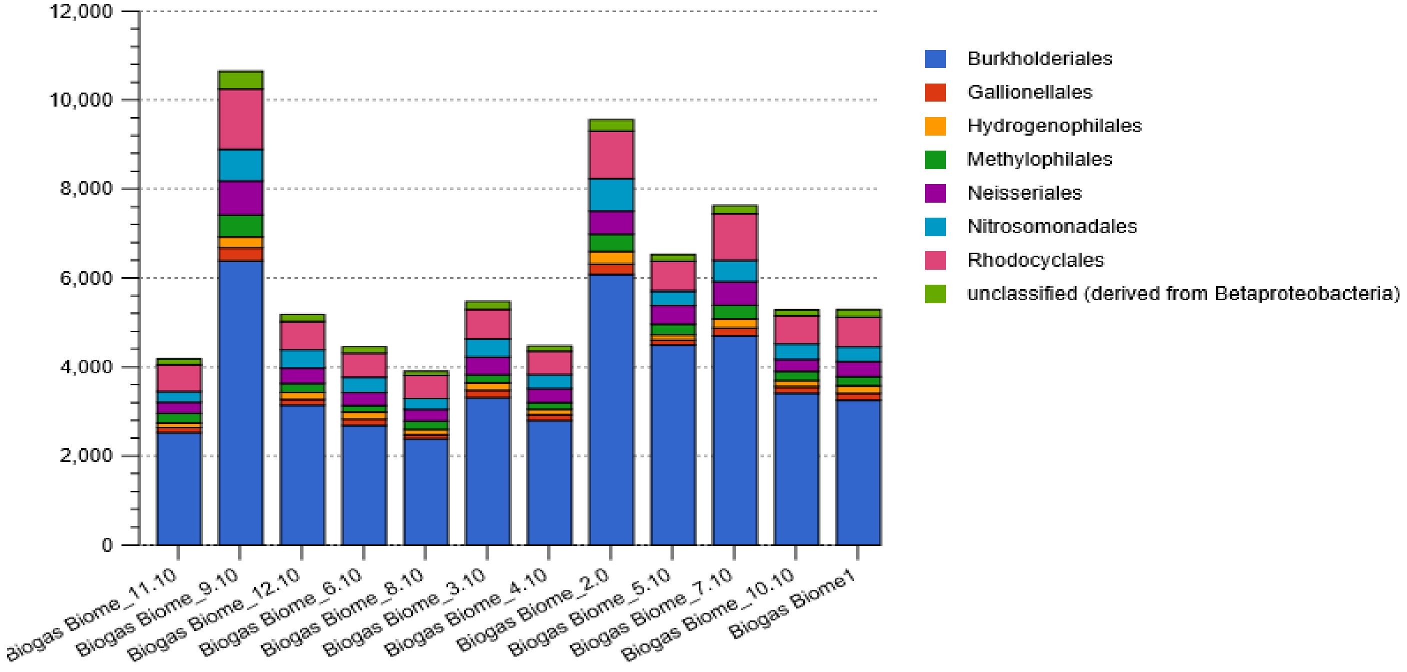


b

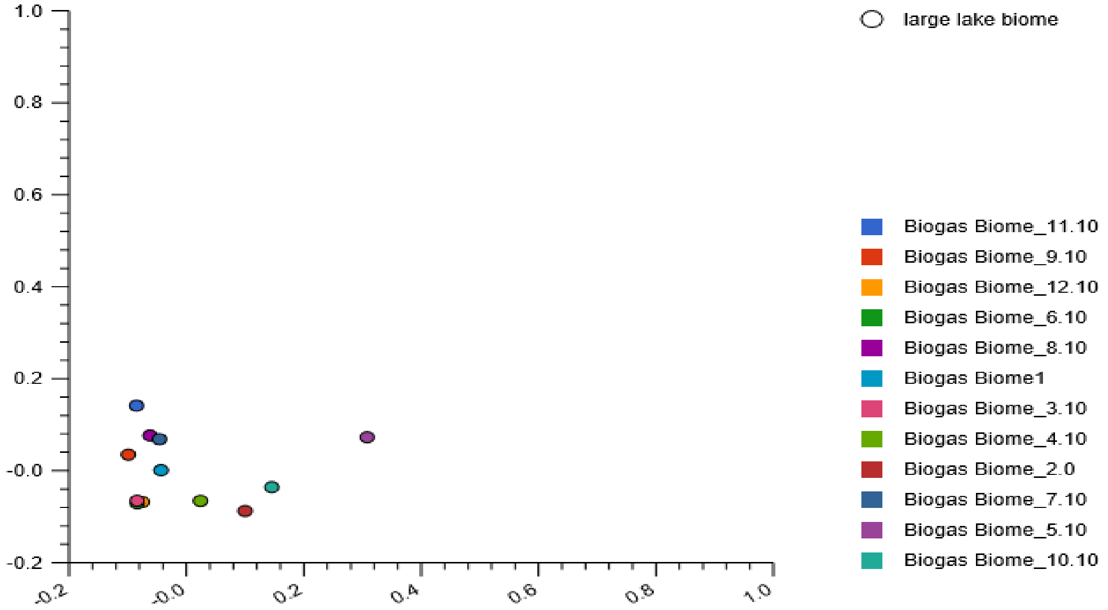


**SFig. 8:** Stacked barchat showing seven β-proteobacteria orders, relative abundances (a) and their PCoA plot (b) revealing nucleotide composition (dis)similarities among the reactors. The plots were based on the Euclidean model. The plots revealed that the nucleotide composition in reactor 7 and 8 clustered partially while the nucleotide composition in reactor 3, 6 and 12 clustered in the lower left quadrant of the plot. The majority of the reactors comprised dissimilar nucleotide composition of the detected β-
