## Supplementary Fig. 9 for "Metagenomics survey unravels diversities of biogas’ microbiomes with potential to enhance its’ productivity in Kenya"

a

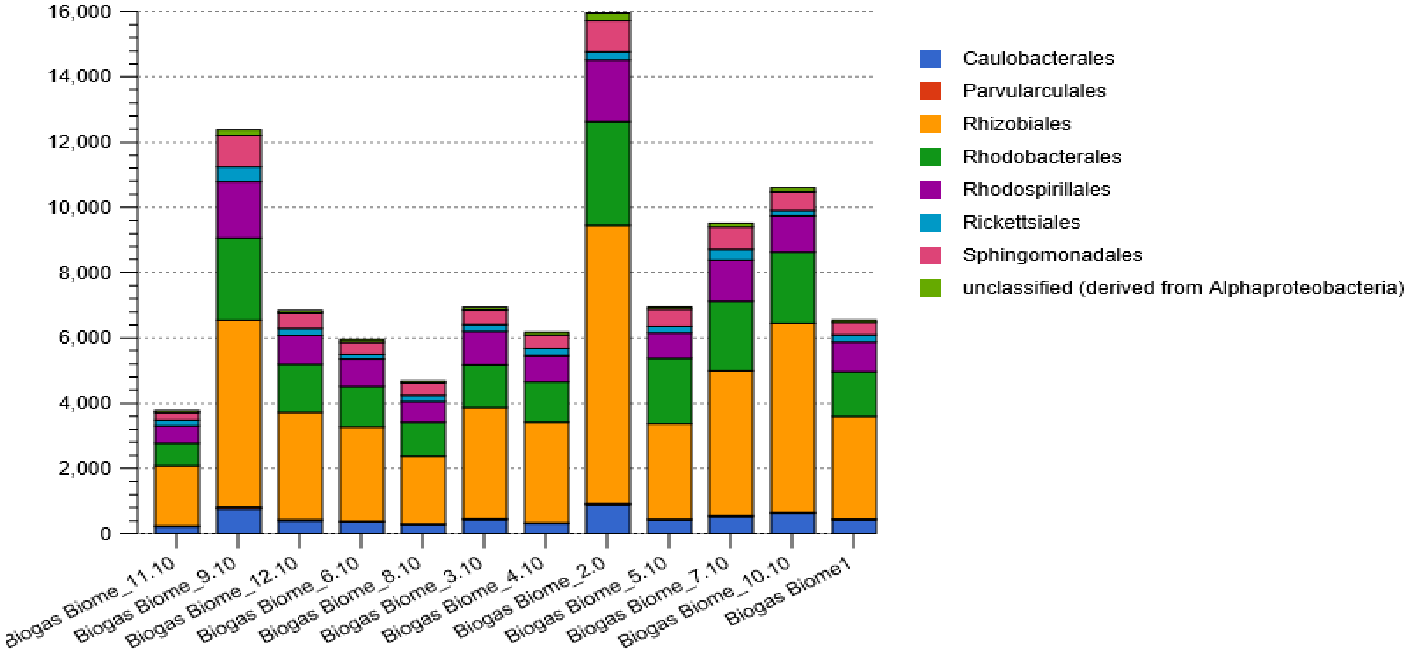


b

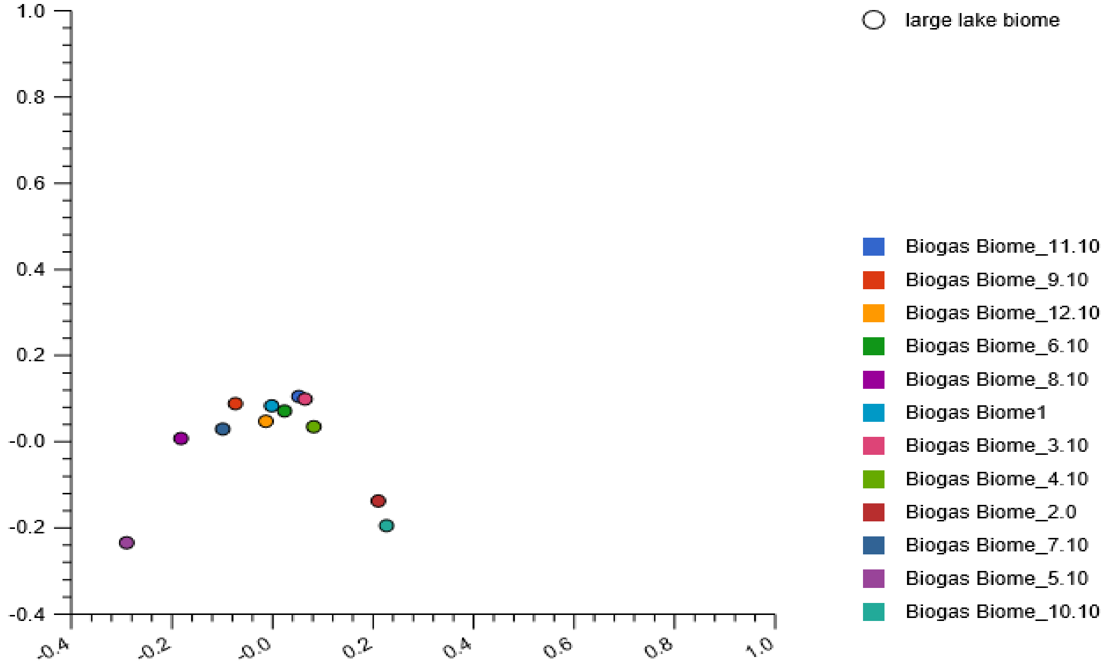


**SFig. 9:** Stacked barchat (a) showing the seven *α-Proteobacteria* orders, relative abundances and their PCoA plots (b), revealing the nucleotide composition variation among the reactors based on the Euclidean model. The plots revealed that the nucleotide composition in reactor 1 and 6 were in close proximity, located on the y-axis while those in reactor 3 and 11 partially clustered, located on the upper right quadrant of the plot. However, the nucleotide composition in the rest of the reactors were dissimilar.
