## Supplementary Fig. 10 for "Metagenomics survey unravels diversities of biogas’ microbiomes with potential to enhance its’ productivity in Kenya"

a

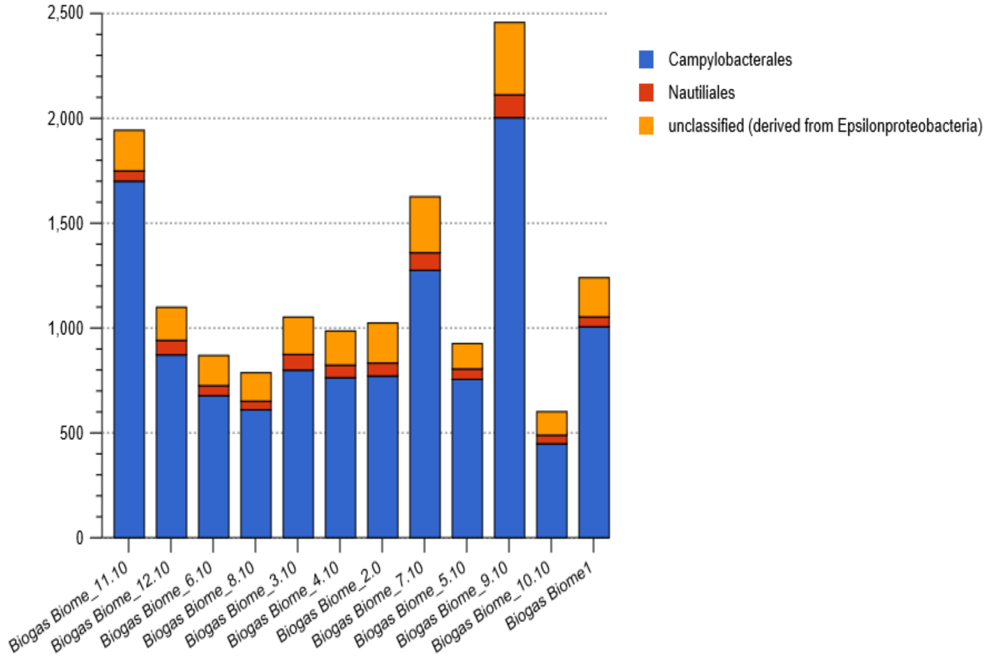


b

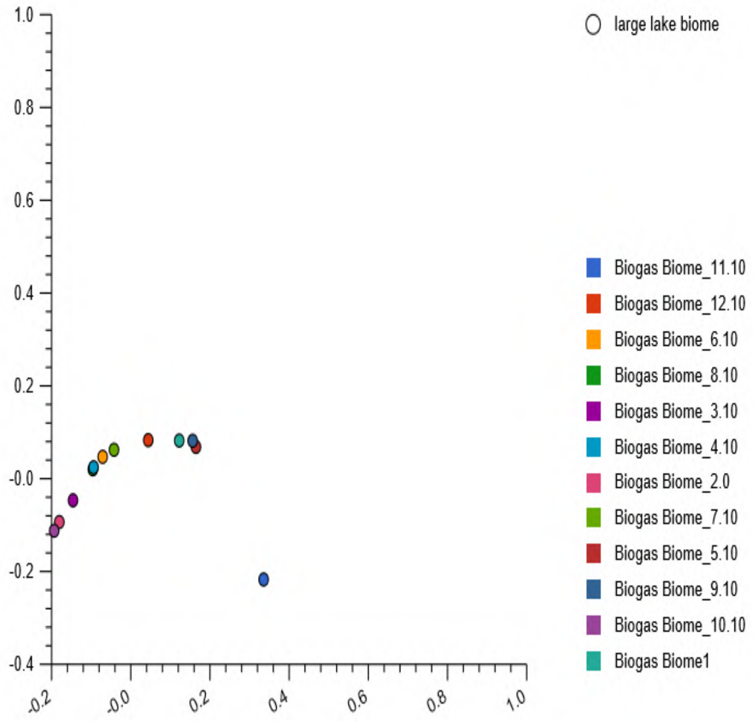


**SFig. 10:** Stacked barchat (a) showing two ε-Proteobacteria orders, relative abundances and their PCoA plot (b) based on Euclidean model. The PCoA plot for nucleotide composition in reactor 4 and 8 clustered in the upper left quadrant and those in reactor 5 and 9 partially clustered in the upper right quadrant of the plot.
