## Supplementary Fig. 11 for "Metagenomics survey unravels diversities of biogas’ microbiomes with potential to enhance its’ productivity in Kenya"

a

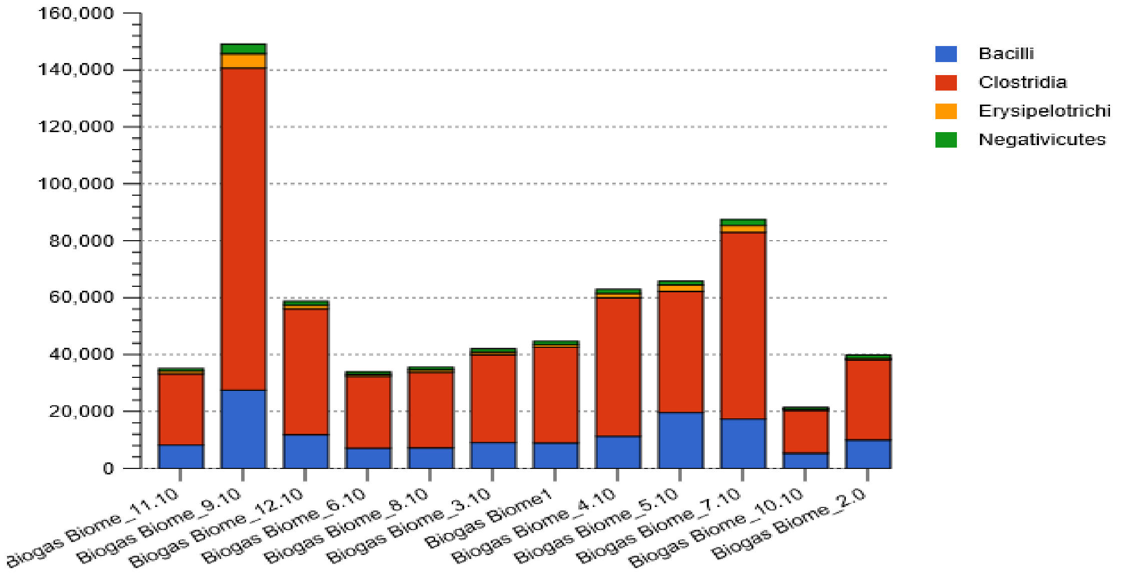


b

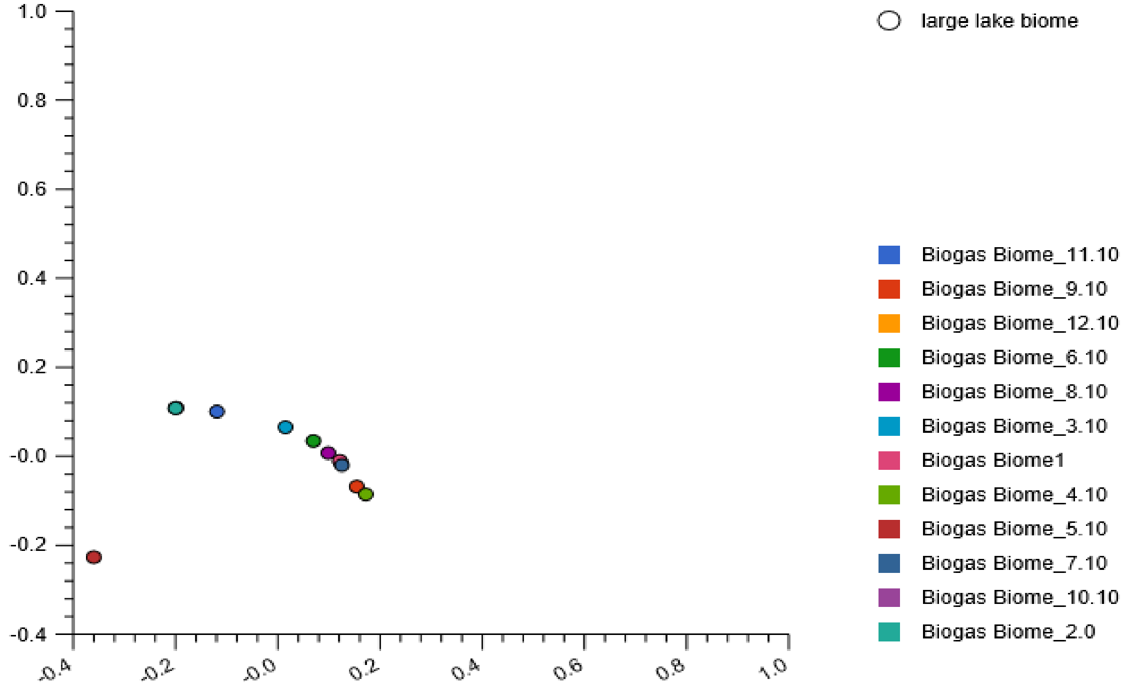


**SFig. 11:** The stacked barchat (a) revealing four *Firmicutes* classes, the relative abundances and their PCoA plots (b), revealing variance among the twelve reactors based on the Euclidean model. The PCoA plots revealed that reactor 4 and 9 were in close proximity, reactor 1 and 7 partially clustered while reactor 1 and 8 were in close proximity. The nucleotide composition in reactor 2, 10 and 12 clustered in lower left quandrant of the plot. The nucleotide composition in the other reactors were dissimilar with reactor 5 singly located on the upper left quadrant of the plot.
