## Supplementary Fig. 12 for "Metagenomics survey unravels diversities of biogas’ microbiomes with potential to enhance its’ productivity in Kenya"

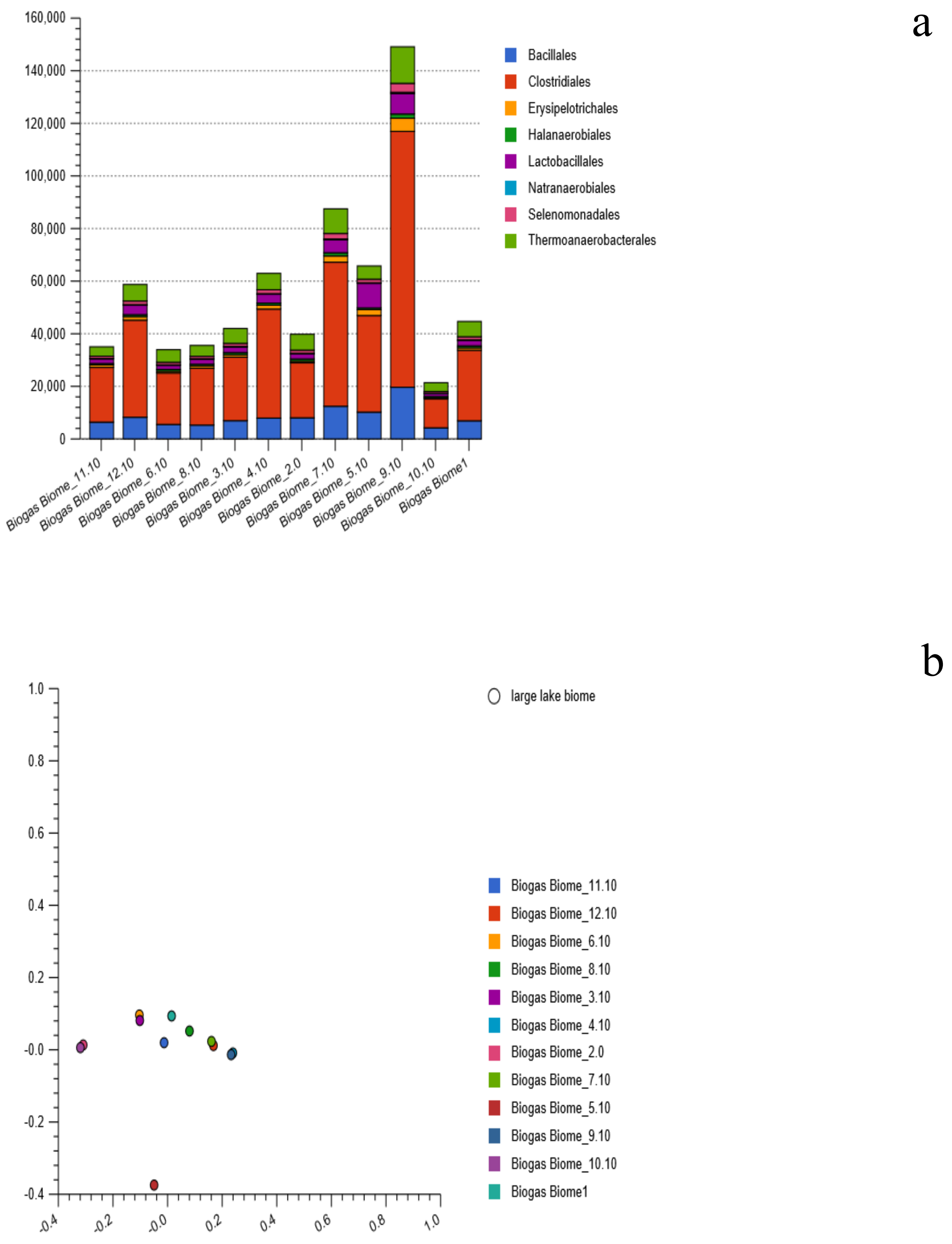


**SFig. 12:** Stacked barchat (a) showing eight *Firmicute*'s orders, relative abundances and their PCoA plot (b) based on the Euclidean model. The plot revealed partial nucleotide reads similarities between reactor 2 and 10, reactor 3 and 6, and reactor 7
