## Supplementary Fig. 13 for "Metagenomics survey unravels diversities of biogas’ microbiomes with potential to enhance its’ productivity in Kenya"

a

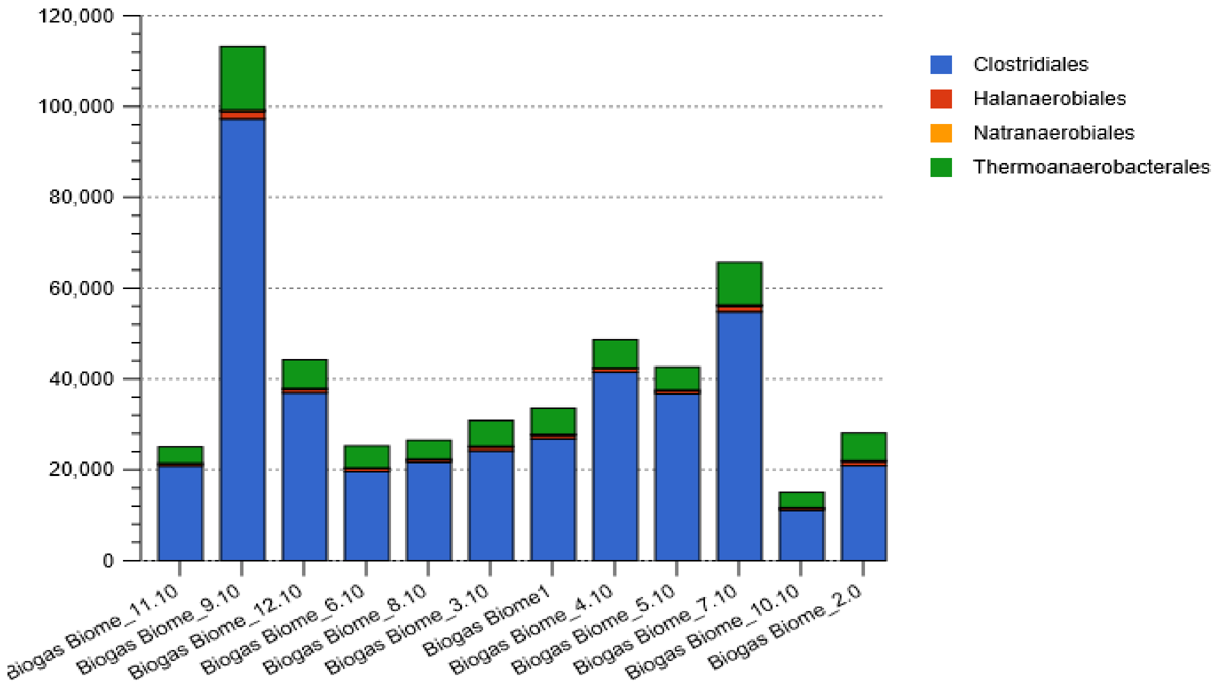


b

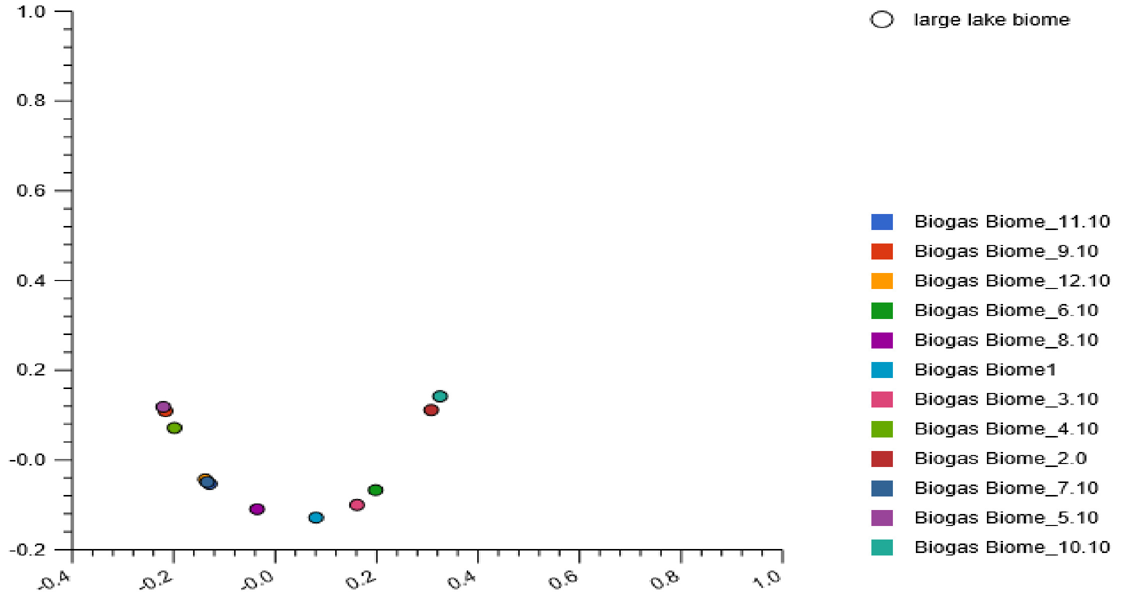


**SFig. 13:** Stacked barchat (a) showing the four *Clostridia* orders and their PCoA plots (b), revealing nucleotide composition variations among the twelve reactors based on the Euclidean model. The plots revealed nucleotide composition dissimilarities among the majority of the reactors. However, the nucleotide composition in reactor 5 and 9 partially clustered in the upper left quadrant of the plot while the nucleotide composition in reactor 7,11 and12 clustered in the lower left quadrant of the plot.
