## Supplementary Fig. 14 for "Metagenomics survey unravels diversities of biogas’ microbiomes with potential to enhance its’ productivity in Kenya"

a

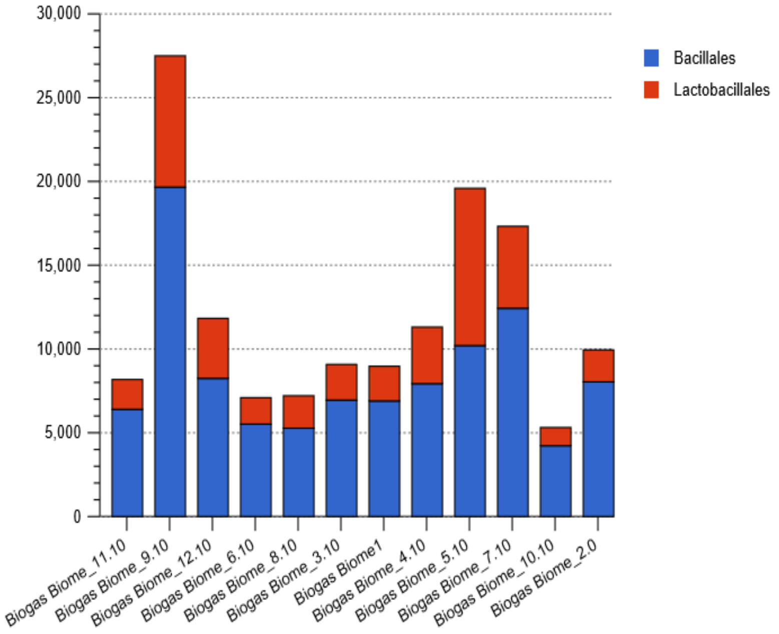


b

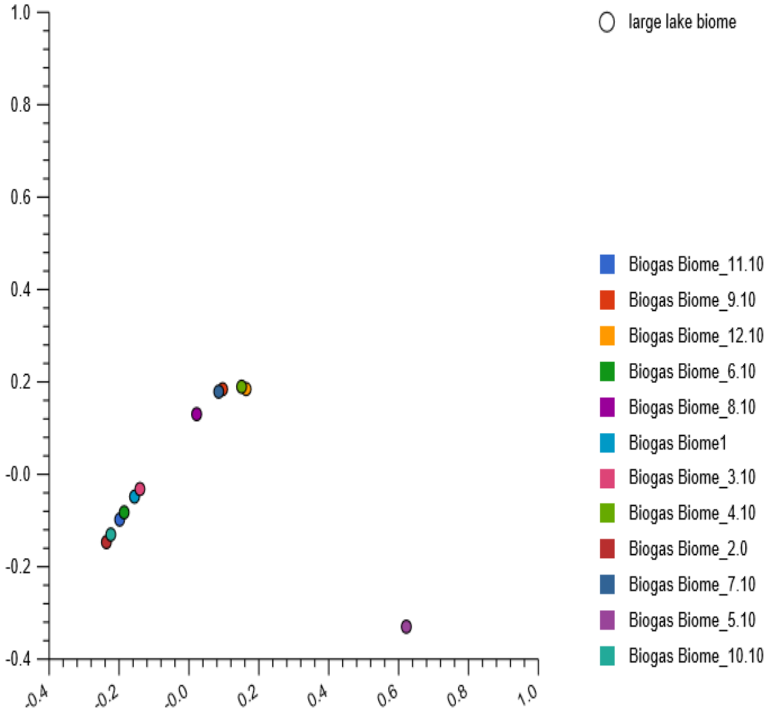


**SFig. 14:** Stacked barchat (a) showing two Bacilli orders, relative abundances and their PCoA plot (b) based on the Euclidean model. The nucleotide composition varied among the reactors. However, the partial clustering of the nucleotide occurred in pair-wise and only nucleotide of two reactors (reactor 5 and 8) were distinctively positioned within the plot.
