## Supplementary Fig. 15 for "Metagenomics survey unravels diversities of biogas’ microbiomes with potential to enhance its’ productivity in Kenya"

a

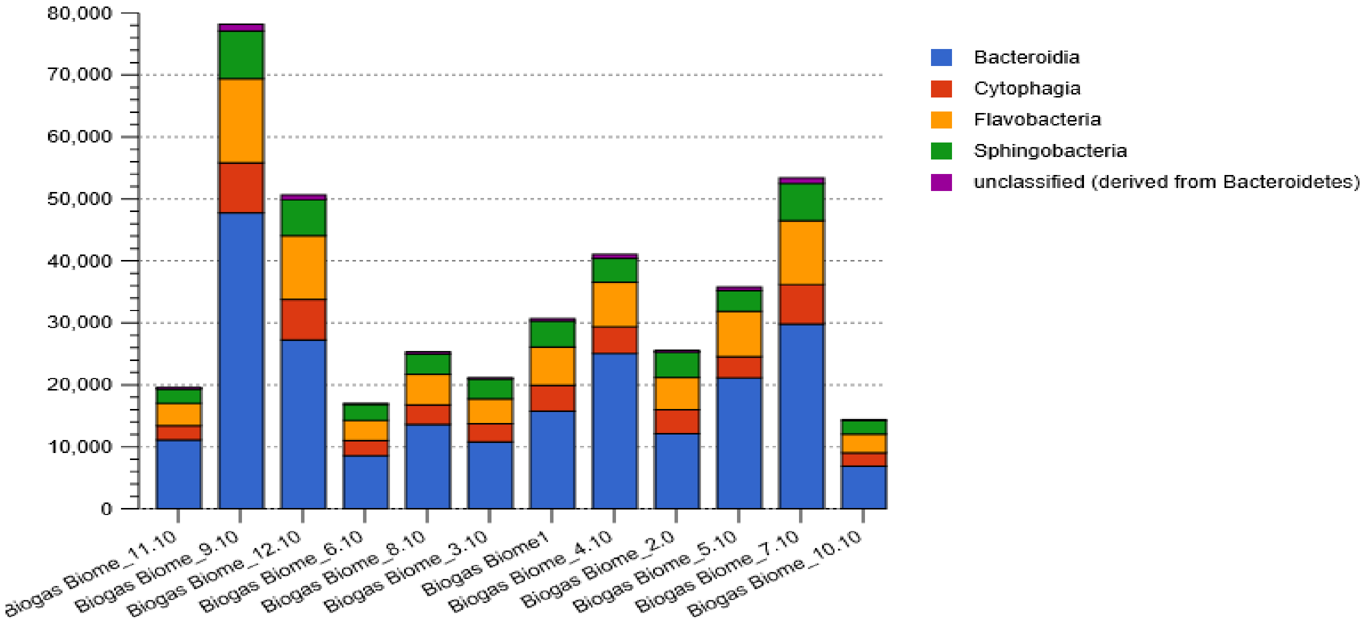


b

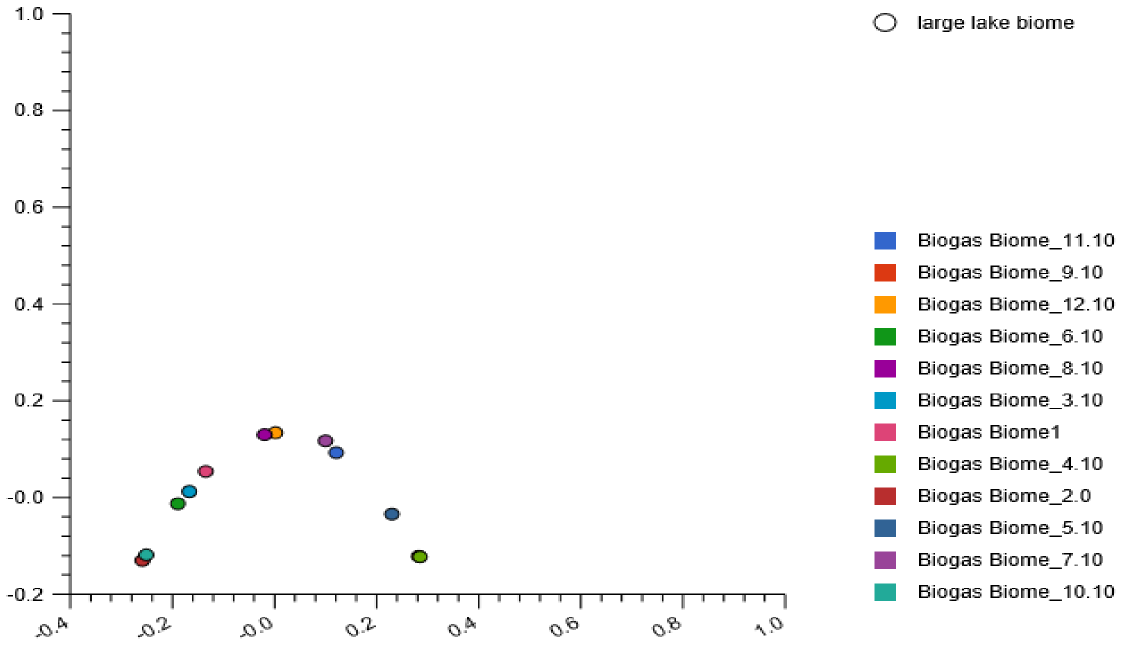


**SFig. 15:** The stacked barchat (a) revealing four *Bacteroidete*s classes, relative abundances and their PCoA plots (b) revealing nucleotide composition variation, based on the Euclidean model. The plots on the nucleotide composition variance among the reactors revealed that reactor 2 and 10 partially clustered on the negative and those in reactor 8 and 12 partially clustered in the lower left quadrant of the plot while those detected in reactor 4 and 9 clustered in the lower right quadrant of the plot.
