## Supplementary Fig. 16 for "Metagenomics survey unravels diversities of biogas’ microbiomes with potential to enhance its’ productivity in Kenya"

a

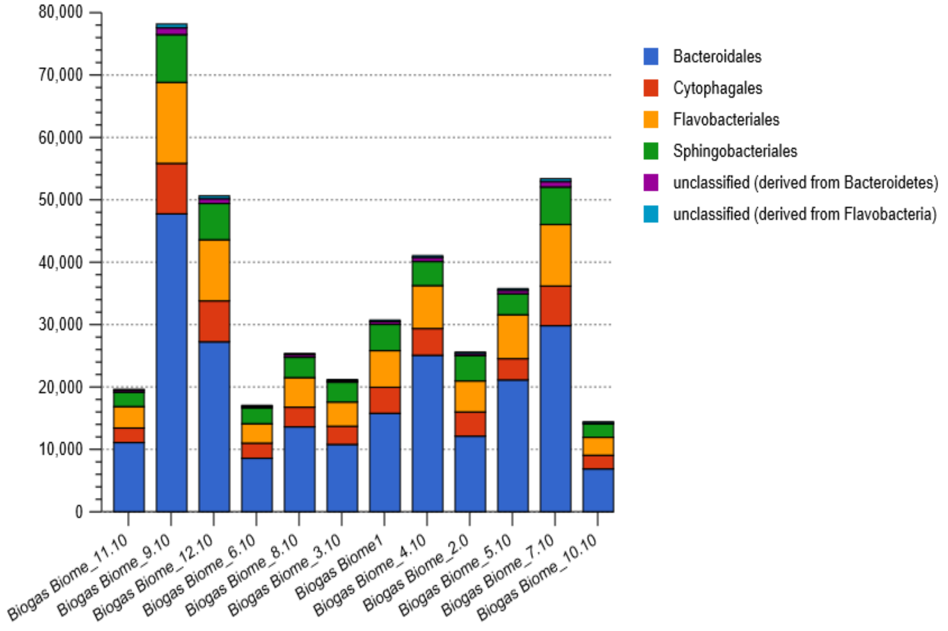


b

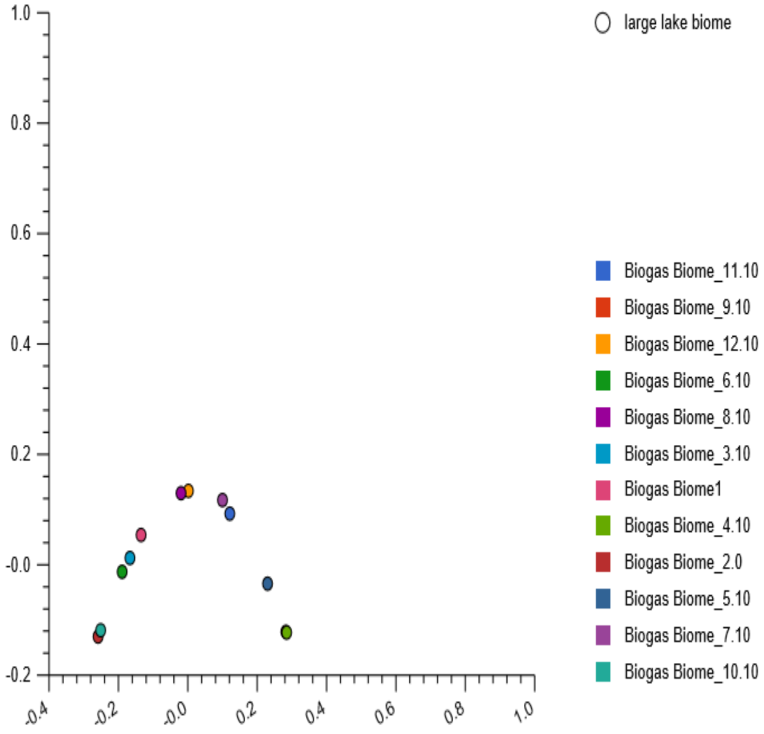


**SFig. 16:** Stacked barchat (a) showing the four *Bacteroidete*'s orders, the relative abundances and their PCoA plot (b) based on the Euclidean model. The nucleotide composition for reactor 2 and 10 and those in reactor 8 and 12 partially clustered in the lower left quadrant of the plot while those detected in reactor 4 and 9 clustered in the lower right quadrant of the plot.
