## Supplementary Fig. 17 for "Metagenomics survey unravels diversities of biogas’ microbiomes with potential to enhance its’ productivity in Kenya"

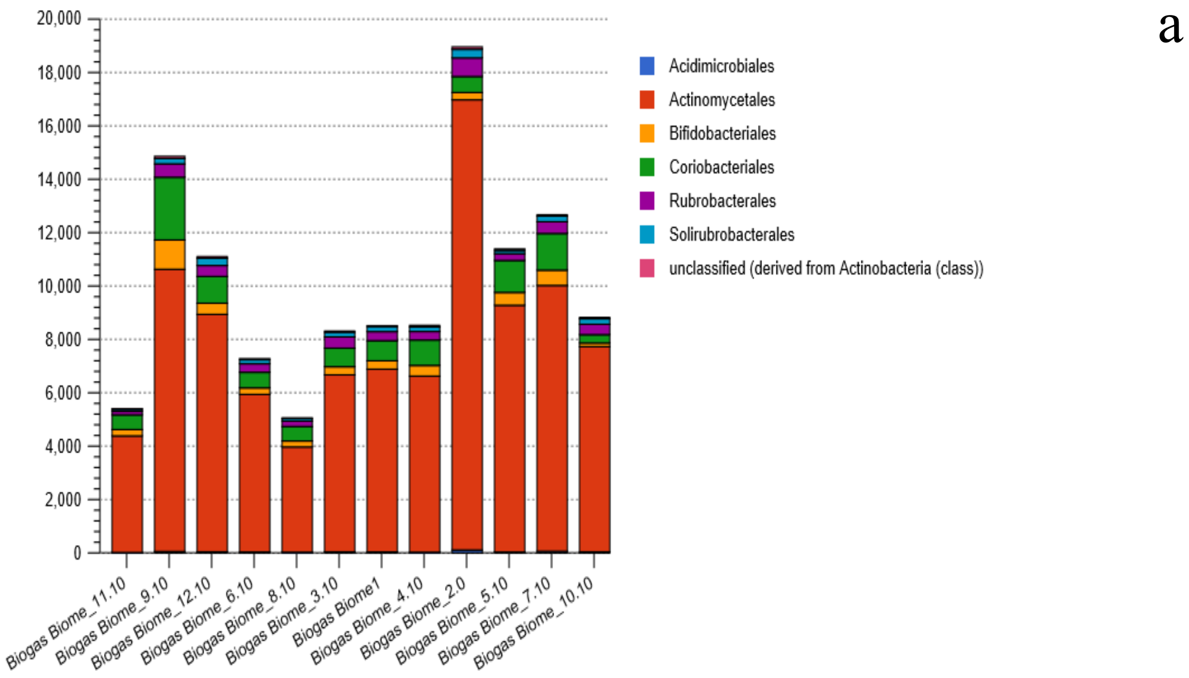


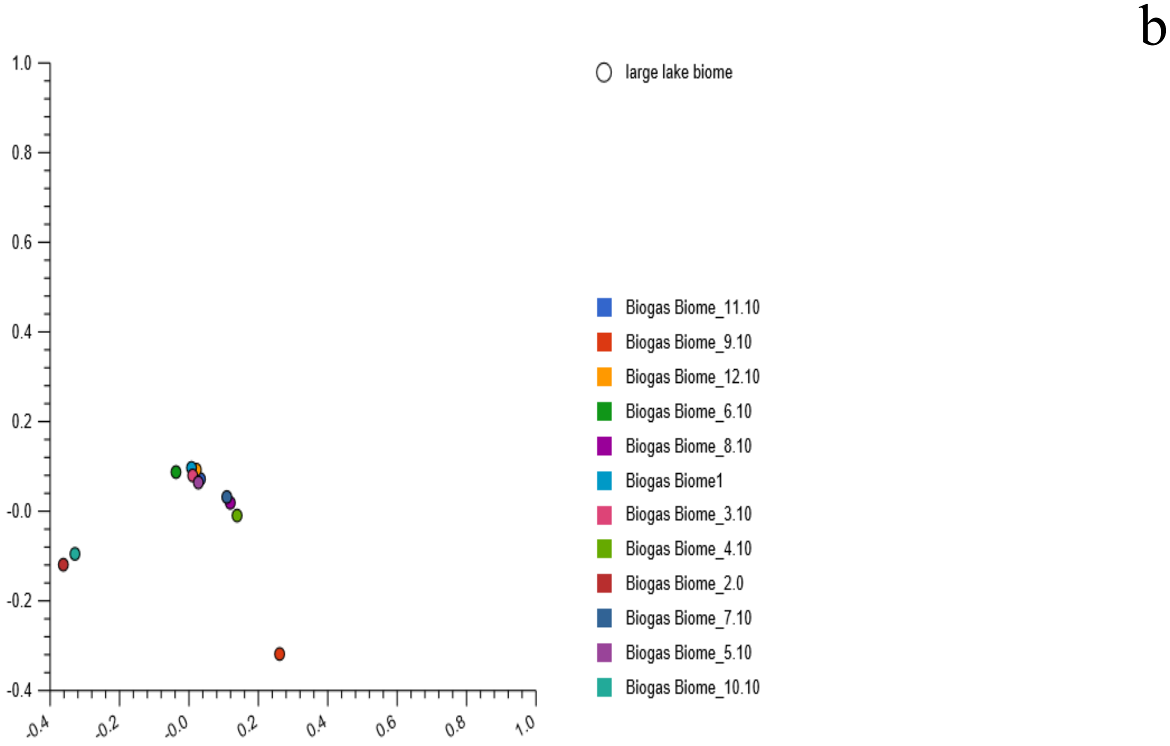


**SFig. 17:** Stacked barchat (a) showing six actinobacteria orders, the relative abundance and their PCoA plot (b) based on the Euclidean model. The nucleotide composition in reactor 7 and 8 in the upper right quadrant of the plot partially clustered, while the nucleotide in reactor 1, 3, 5, 11 and 12 formed a cluster along the y-axis of the plot.
