## Supplementary Fig. 18 for "Metagenomics survey unravels diversities of biogas’ microbiomes with potential to enhance its’ productivity in Kenya"

a

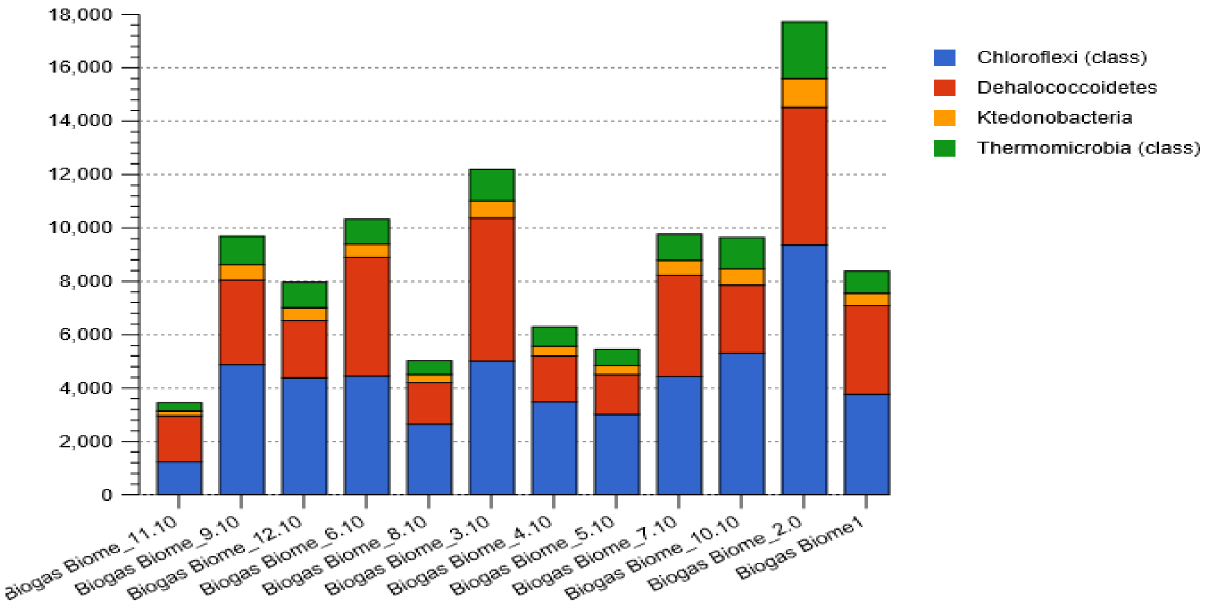


b

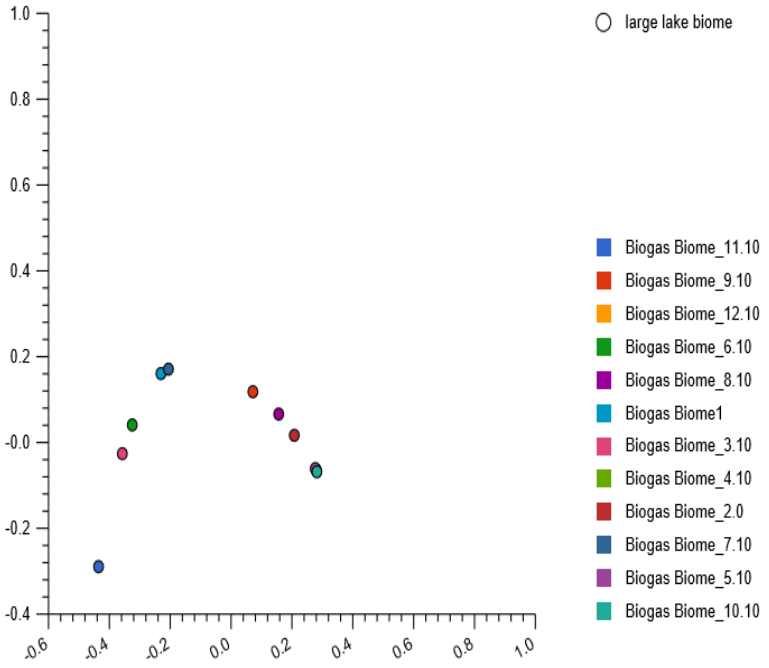


**SFig. 18:** The stacked barchat (a) showing four *Chloroflexi* classes, relative abundances and PCoA plot (b) revealing variance among the twelve reactors, based on the Euclidean model. The plots revealed that reactor 1 and 7 were in close proximity located on the upper left quandrant of the plot. The nucleotide composition in reactor 4, 5, 10 and 12 clustered in the lower right quandrant of the plot.
