## Supplementary Fig. 19 for "Metagenomics survey unravels diversities of biogas’ microbiomes with potential to enhance its’ productivity in Kenya"

a

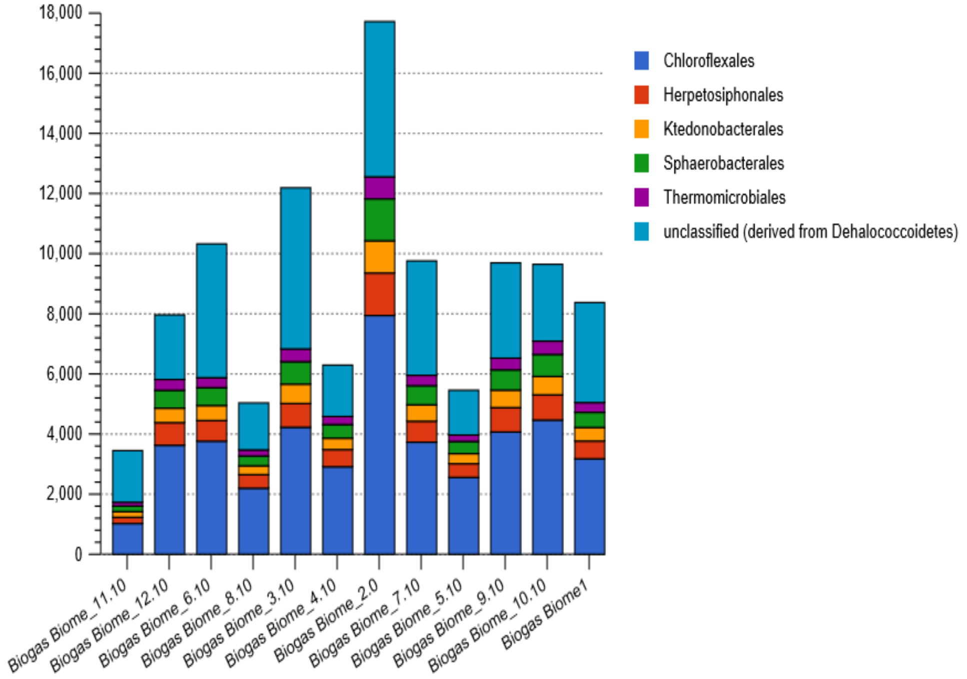


b

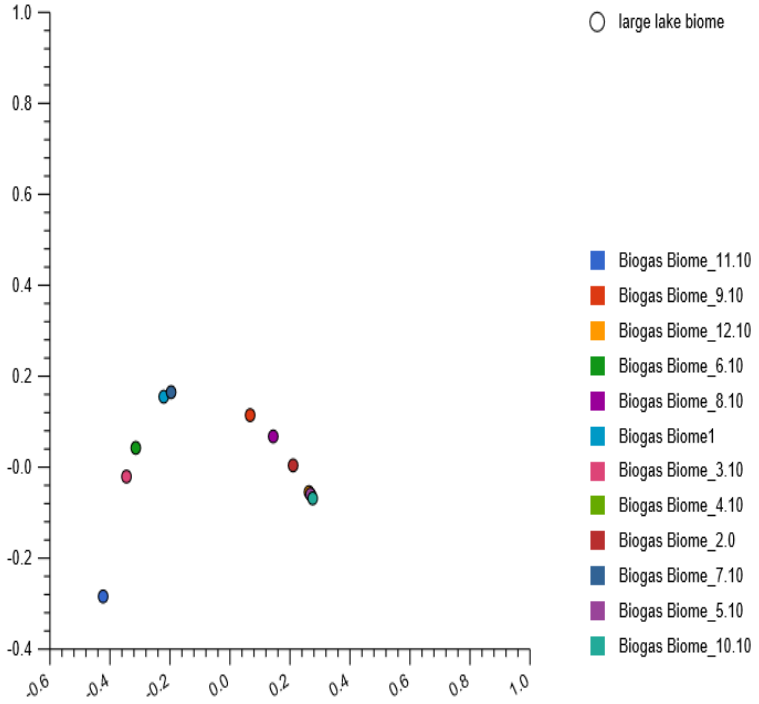


**SFig. 19**: Stacked barchat showing five Chloroflexi orders, relative abundances and their PCoA plot based on the Euclidean model.
