## Supplementary Fig. 20 for "Metagenomics survey unravels diversities of biogas’ microbiomes with potential to enhance its’ productivity in Kenya"

a

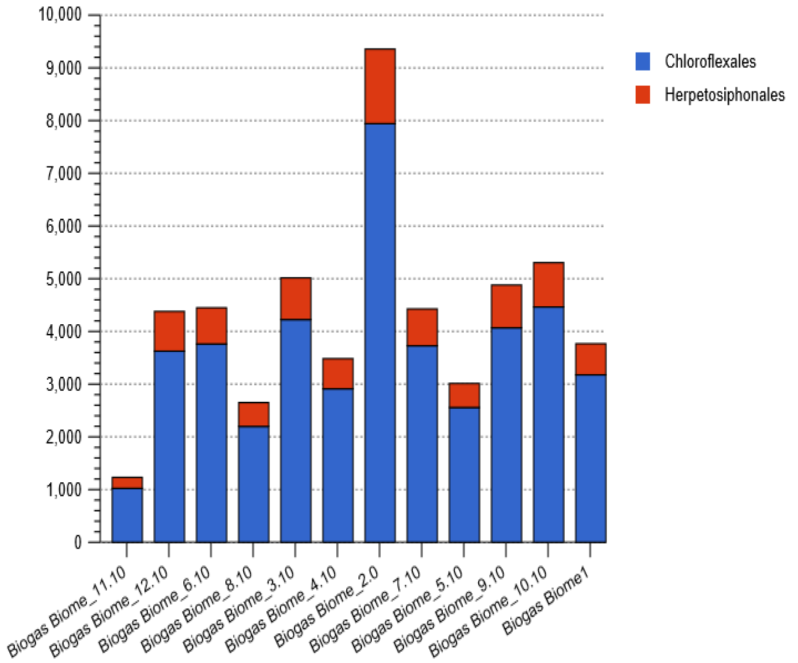


b

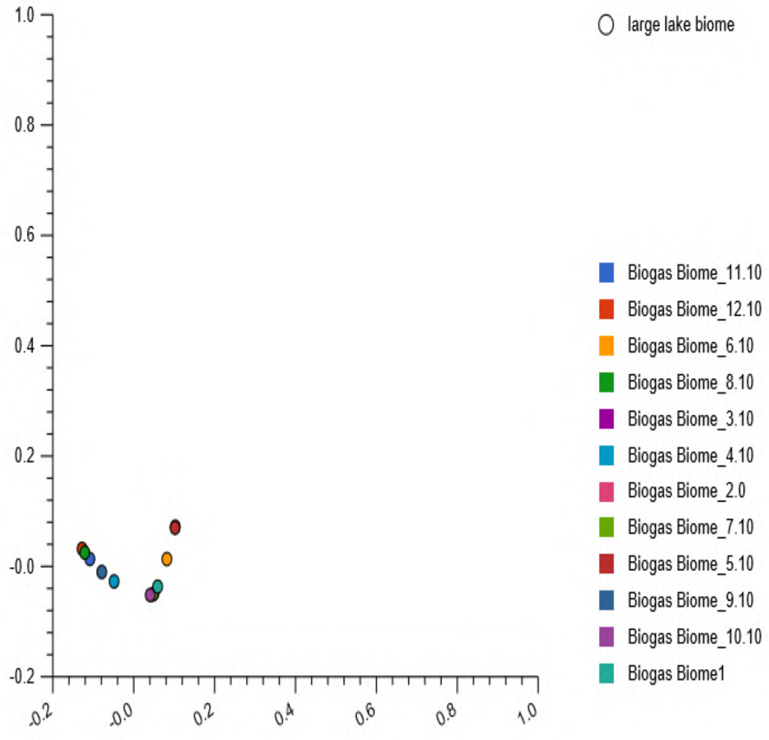


**SFig 20** Stacked barchat (a) showing two Chloroflexi class orders, relative abundances and their PCoA plot (b) based on their Euclidean model. The nucleotide composition in reactor 1 and 5 (clustered; upper right quadrant) and those in reactor 3 and 7 (clustered; lower right quadrant) were similar. Notably, the nucleotide reads in reactor 1 and 10 partially clustered with those detected in reactor 3 and 7. Similarly the nucleotide reads in reactor 8, 11 and 12 partially clustered in the upper left of the quadrant of the plot.
