## Supplementary Fig. 21 for "Metagenomics survey unravels diversities of biogas’ microbiomes with potential to enhance its’ productivity in Kenya"

a

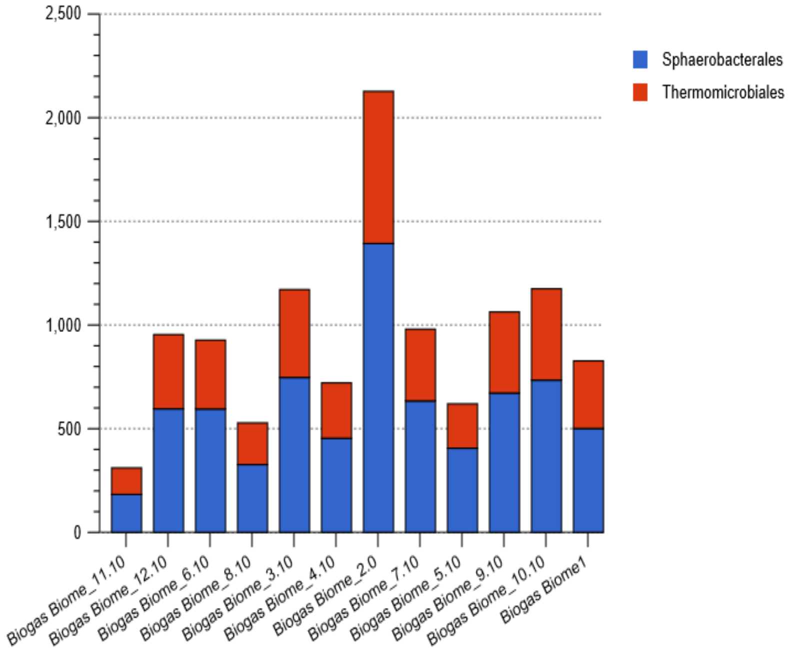


b

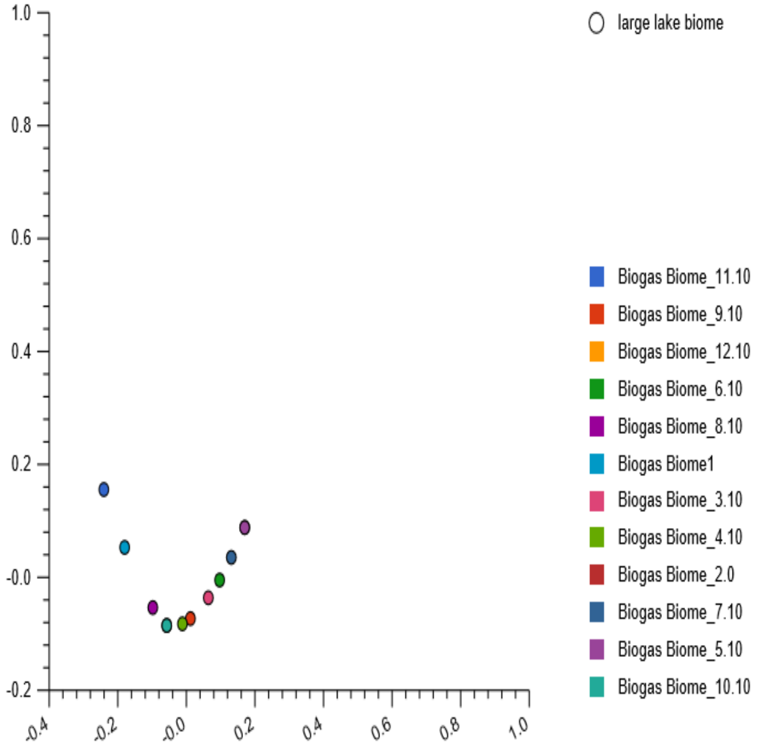


**SFig. 21:** Stacked barchat (a) showing the two *Thermomicrobia* orders, relative abundances and their PCoA plot (b) based on the Euclidean model. The PCoA plot revealed nucleotide composition dissimilarities among the twelve studied reactors.
