## Supplementary Fig. 22 for "Metagenomics survey unravels diversities of biogas’ microbiomes with potential to enhance its’ productivity in Kenya"

a

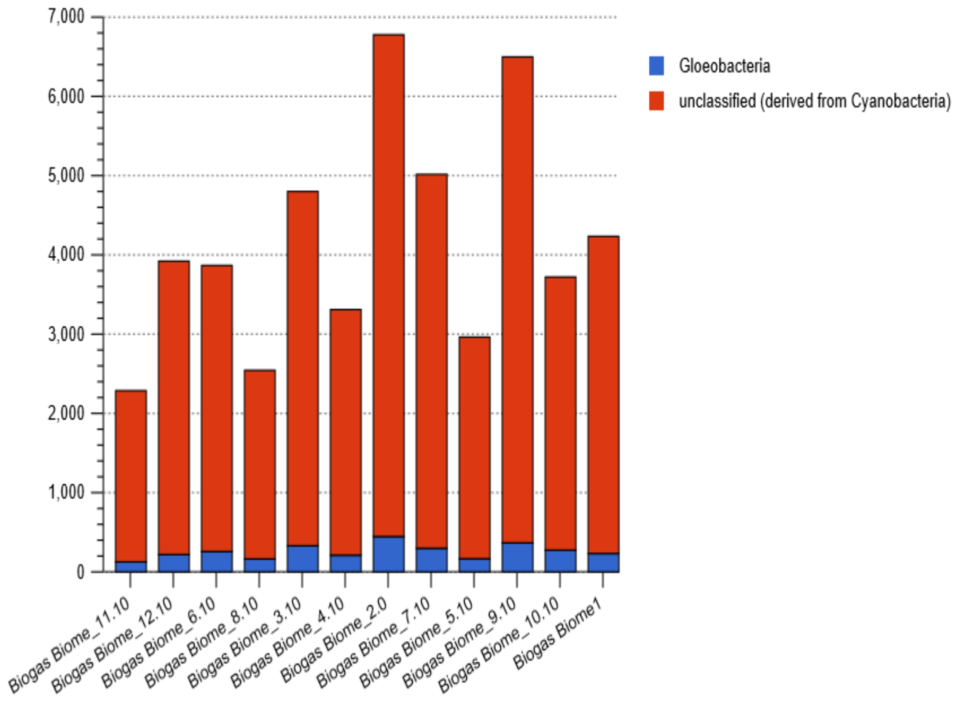


b

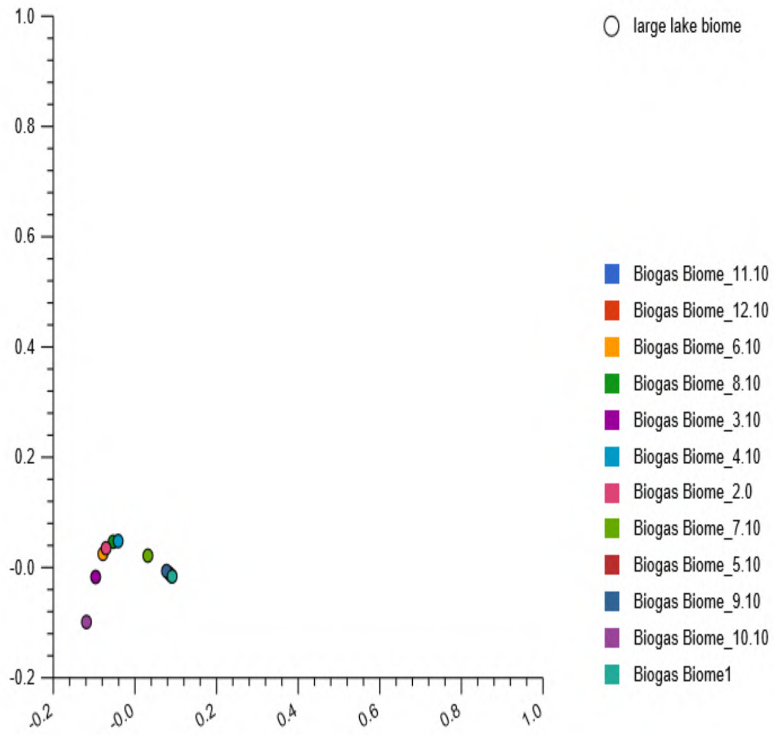


**SFig 22:** Stacked barchat (a) showing *Cyanobacteria* class, relative abundances and their PCoA plot (b) based on the Euclidean model. The plot reveal dissimilarities among the reactors and only the nucleotides composition in the three reactors (reactor 3, 7 and 10) that were distinctively positioned within the plot.
