## Supplementary Fig. 23 for "Metagenomics survey unravels diversities of biogas’ microbiomes with potential to enhance its’ productivity in Kenya"

a

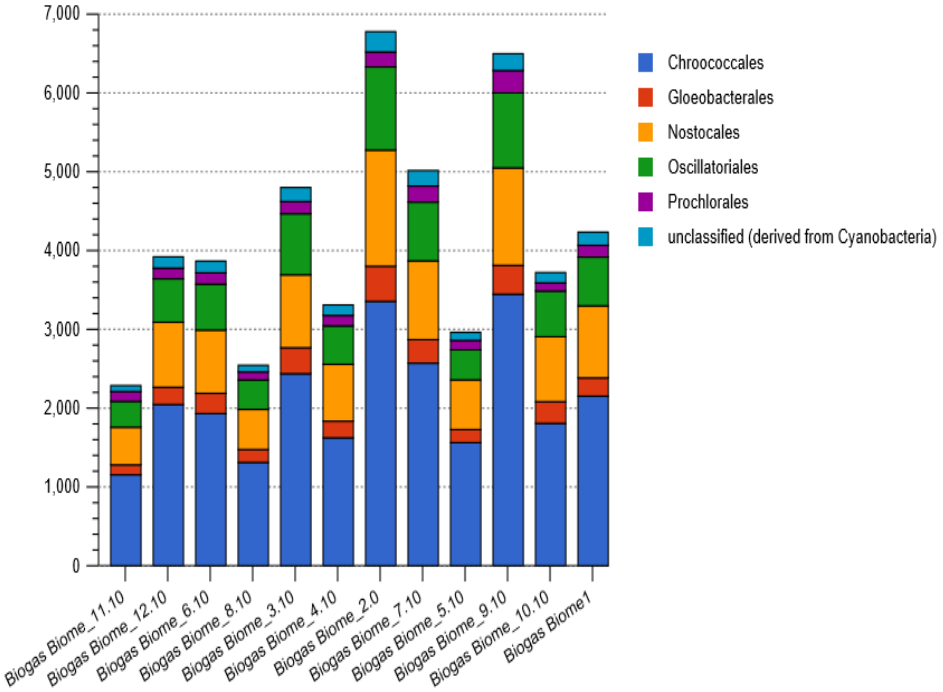


b

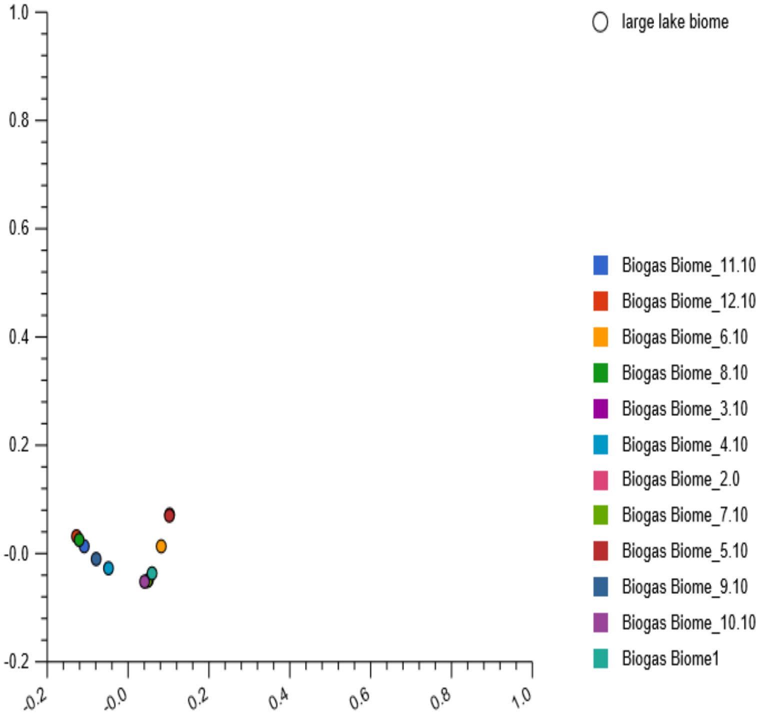


**SFig. 23** Stacked barchat (a) showing the five *Cyanobacteria* orders, relative abundances and their PCoA plot (b) based on the Euclidean model. The PCoA plot revealed four out of twelve reactors that were distinctively located in plots. Other reactors formed
