## Supplementary Fig. 24 for "Metagenomics survey unravels diversities of biogas’ microbiomes with potential to enhance its’ productivity in Kenya"

a

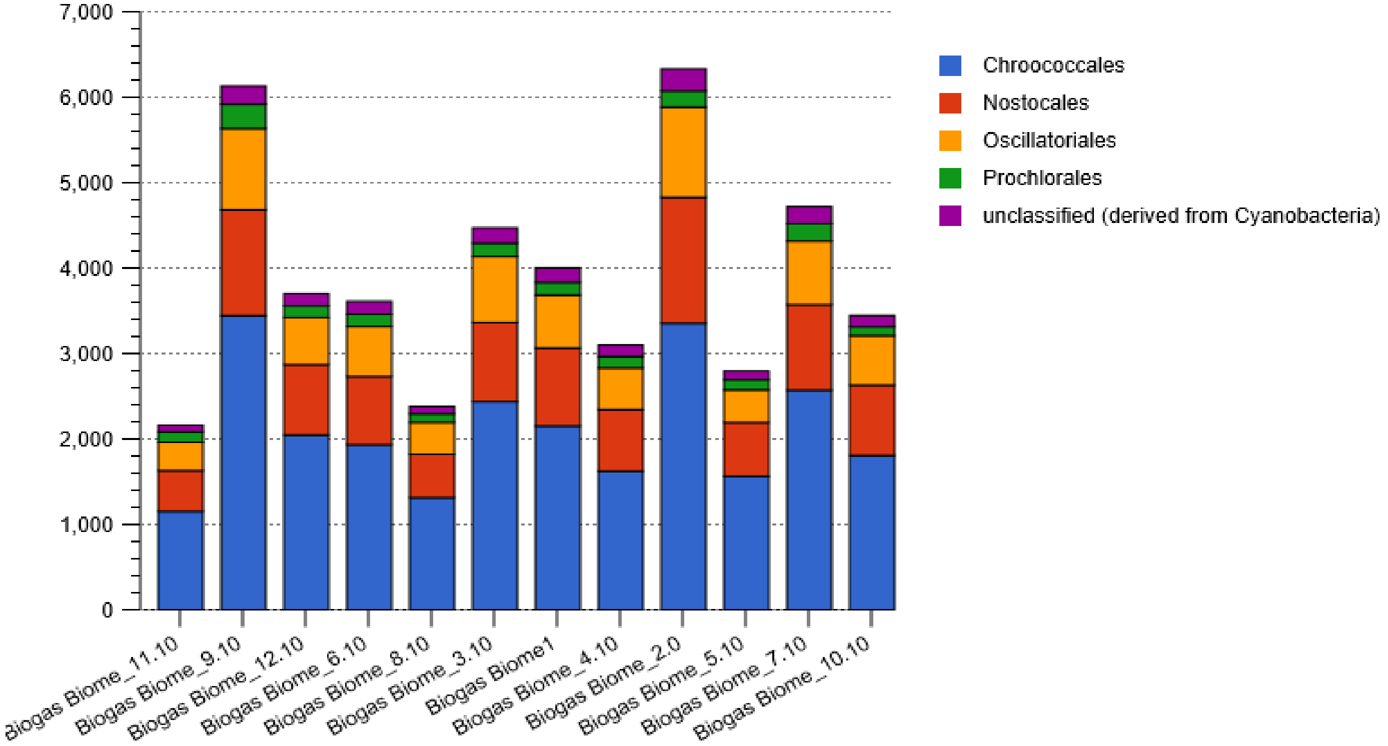


b

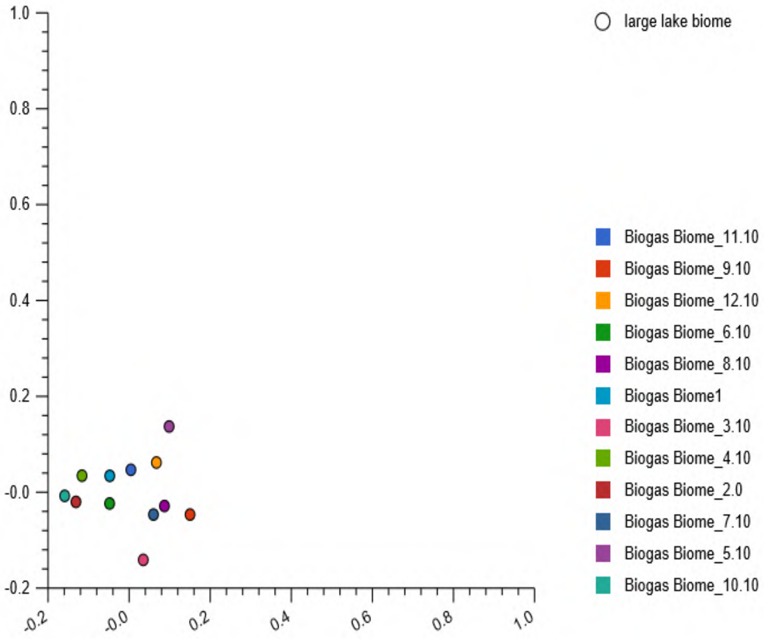


**SFig. 24:** Stacked barchat (a) showing the four affiliates of the Unclassified *Cyanobacteria* nucleotide reads, relative abundances and their PCoA plots (b), based on the Euclidean model. The plot revealed dissimilarities among the reactors, distributed in the four plot quadrant.
