## Supplementary Fig. 26 for "Metagenomics survey unravels diversities of biogas’ microbiomes with potential to enhance its’ productivity in Kenya"

a

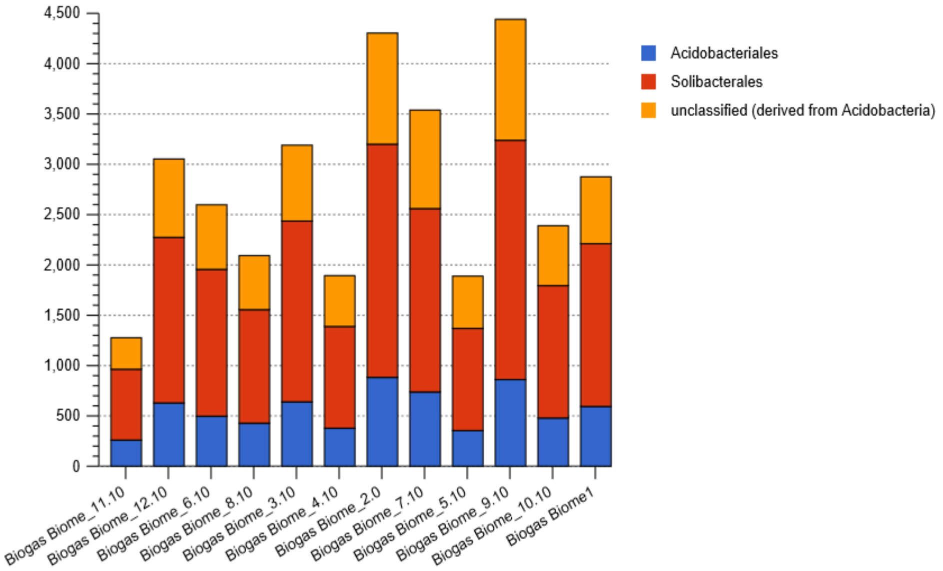


b

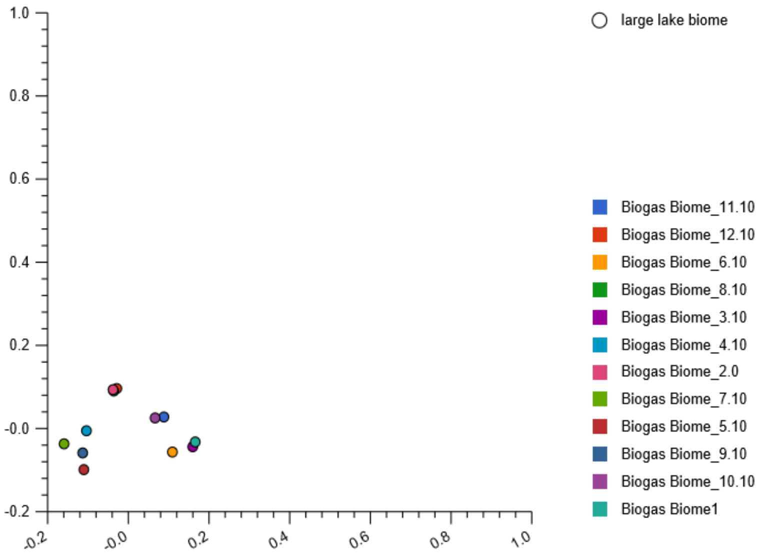


**SFig. 26:** The stacked barchat (a) showing the two *Acidobacteria* orders, relative abundances and their PCoA plots (b) for their nucleotide composition. The nucleotide composition in reactor 1 and 3 partially clustered in the lower right quandrant of the plot, those in reactor 10 and 11 were in close proximity located in the upper right quadrant of the plot while the nucleotide composition in reactor 2, 8 and 12 clustered in the upper right quadrant of the plot.
