## Supplementary Fig. 27 for "Metagenomics survey unravels diversities of biogas’ microbiomes with potential to enhance its’ productivity in Kenya"

a

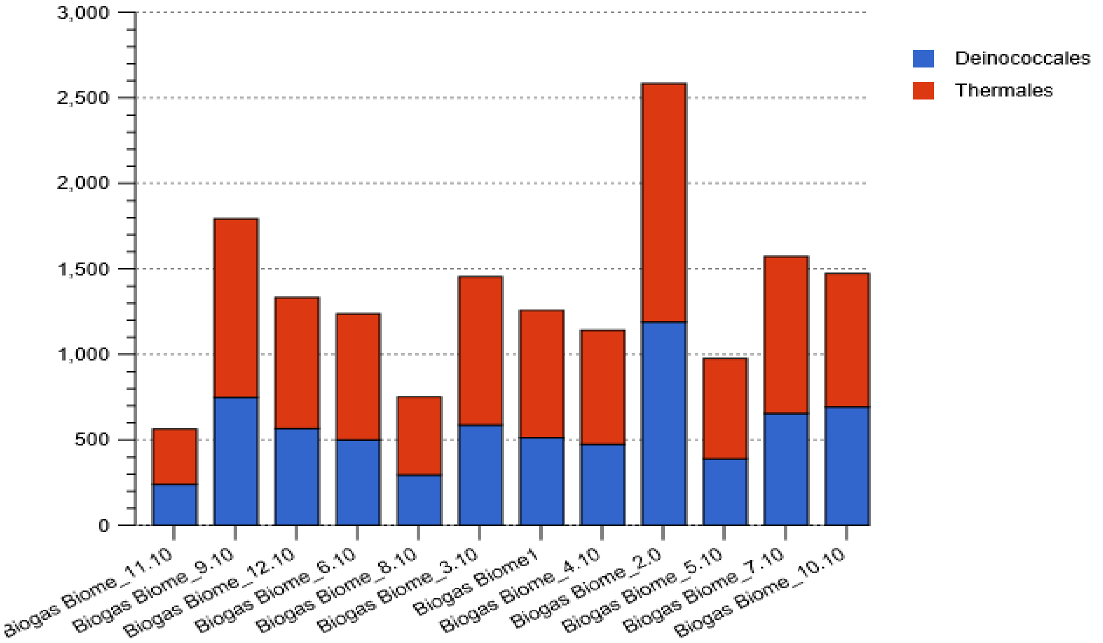


b

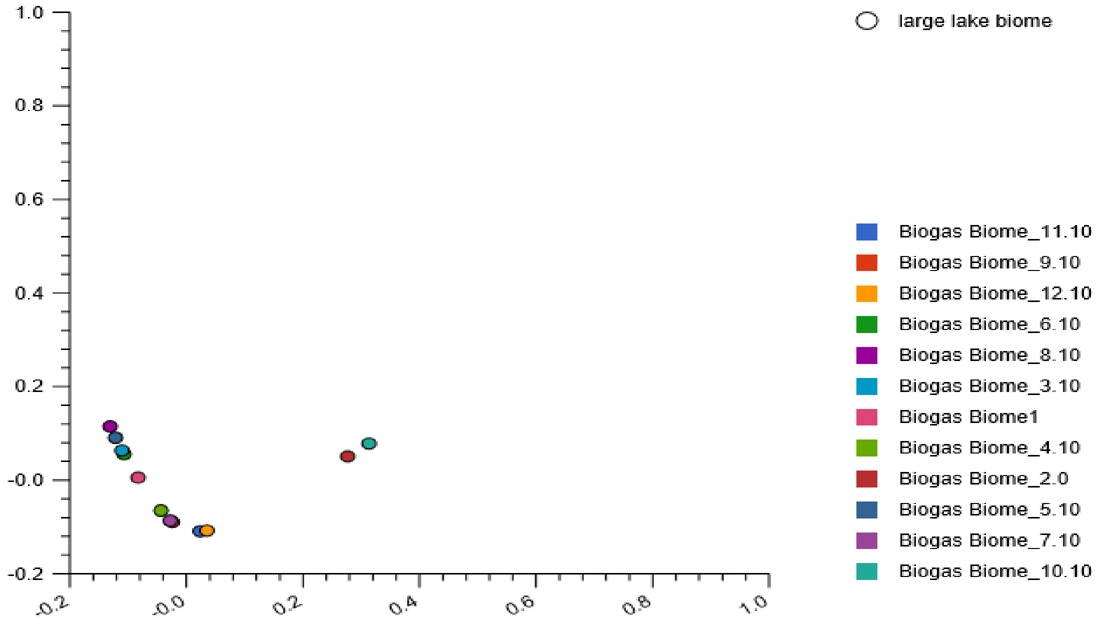


**SFig. 27:** Stacked barchat (a) showing two *Deinococcus-Thermus* orders, the relative abundances and the PCoA plots (b) based on the Euclidean model. The PCoA plots revealed clustering of the nucleotide composition of reactor 3 and 6, and those in reactor 7 and 9 on the upper left quadrant and lower left quadrant of the plot respectively. The nucleotide composition in reactor 4 were in close proximity with those in reactor 7. The nucleotide composition in reactor 11 and 12 partially clustered on the lower right quadrant of the plot. However, those detected in reactor 2 and 10 were the only nucleotide reads located in the upper right quadrant of the plot .
