## Supplementary Fig. 28 for "Metagenomics survey unravels diversities of biogas’ microbiomes with potential to enhance its’ productivity in Kenya"

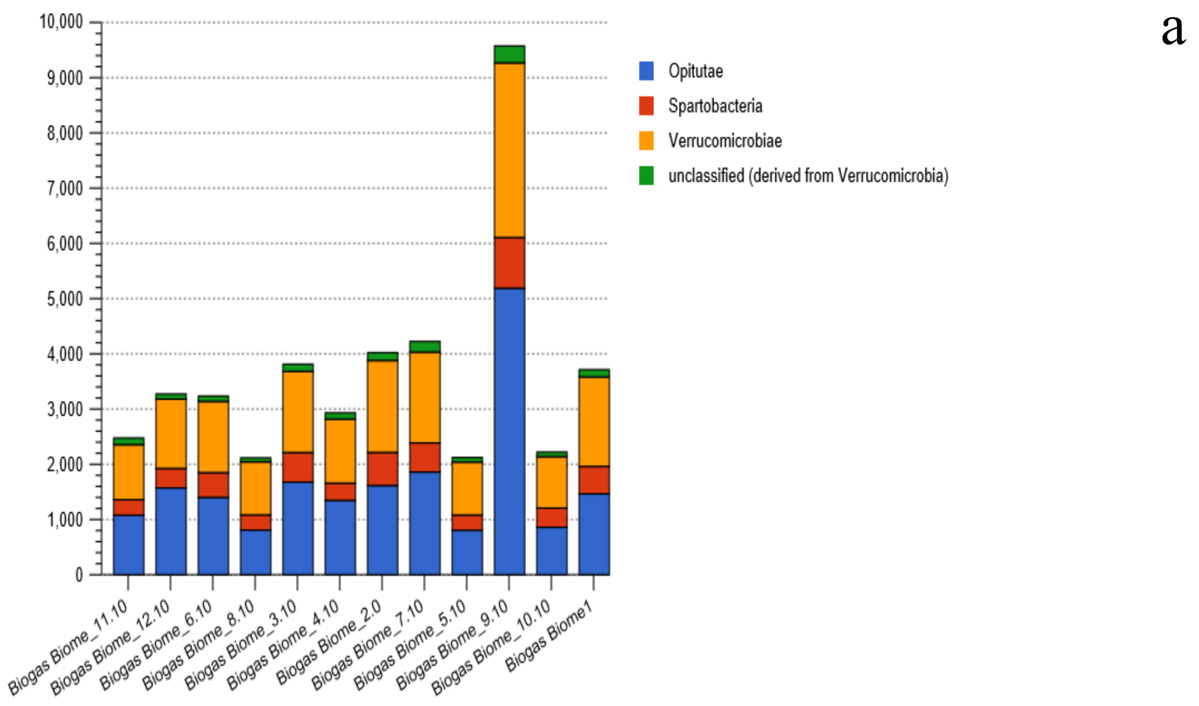


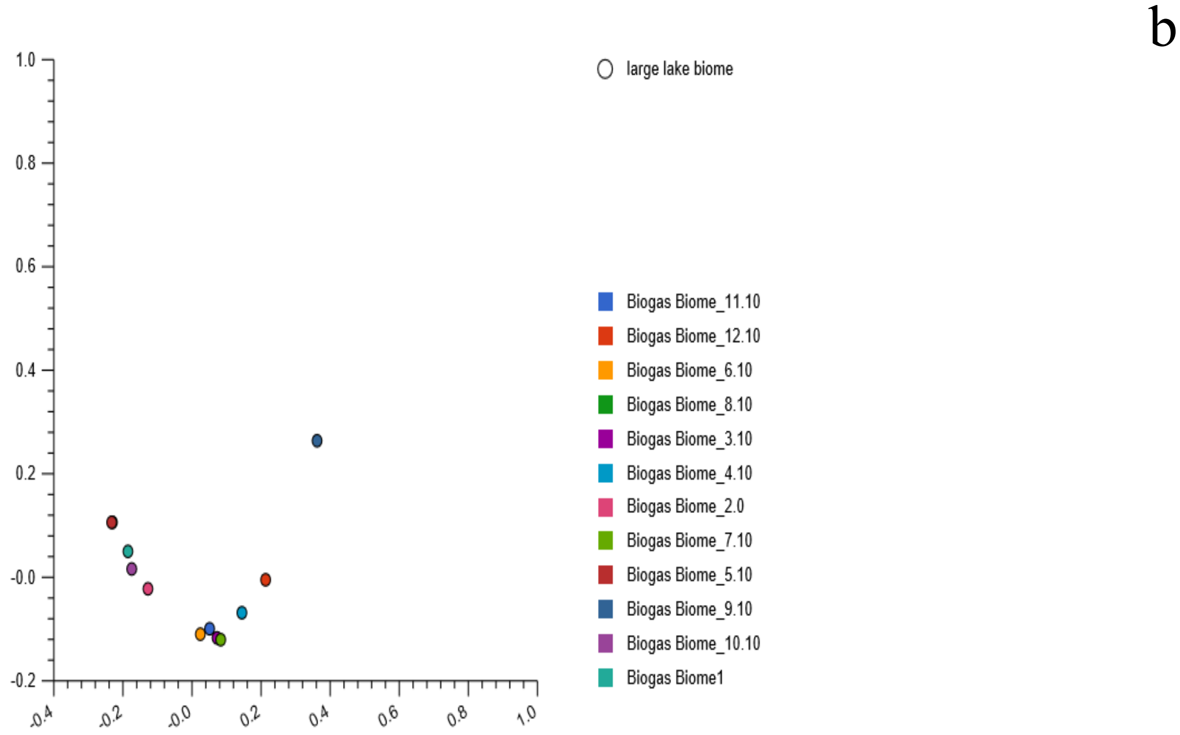


**SFig. 28:** Stacked barchat (a) showing *Verrucomicrobia* classes, relative abundances and their PCoA plot (b) based on the Euclidean model. The PCoA plot revealed nucleotide composition dissimilarities among the reactors, except those detected in reactor 3 and 7 that partially clustered in the lower right quadrant of the plot.
