## Supplementary Fig. 29 for "Metagenomics survey unravels diversities of biogas’ microbiomes with potential to enhance its’ productivity in Kenya"

a

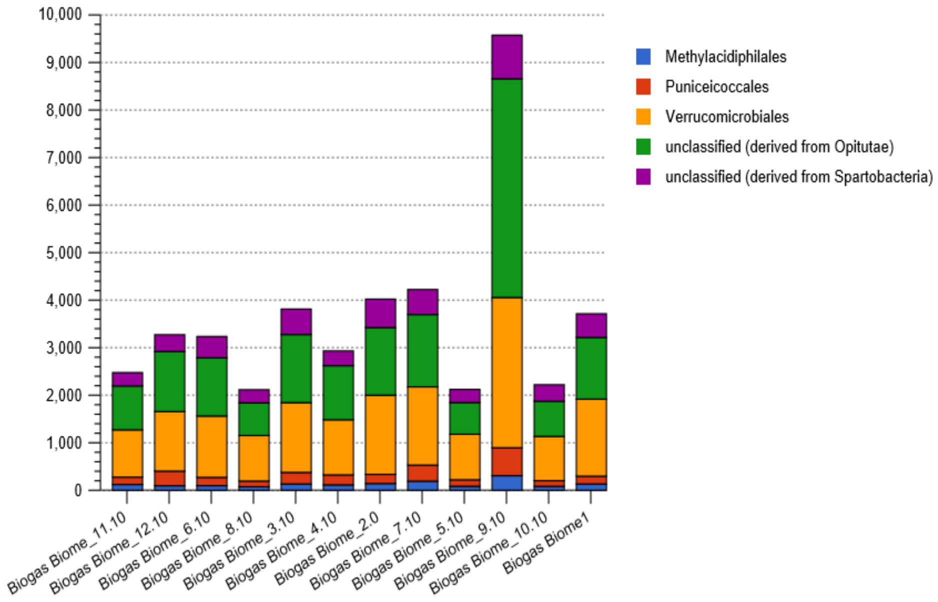


b

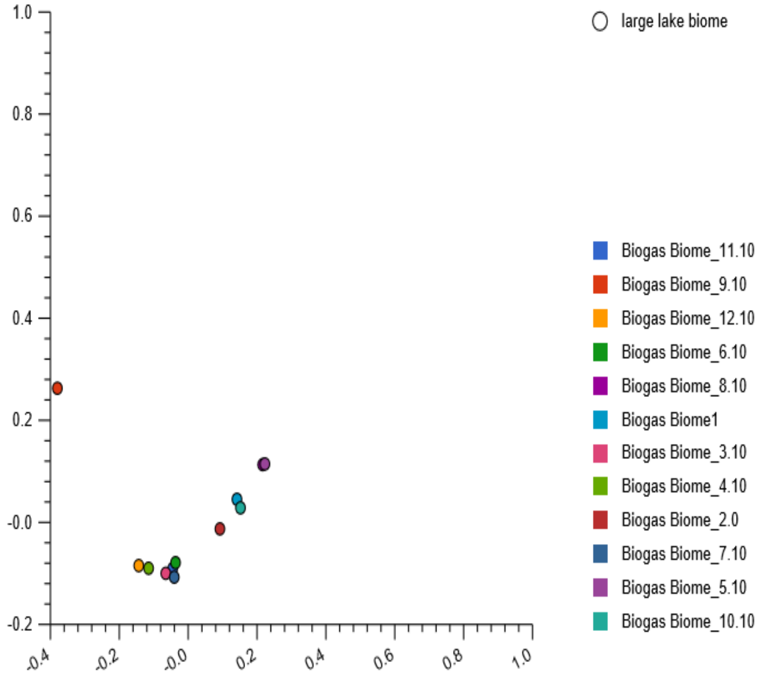


**SFig. 29:** Stacked barchat (a) showing three *Verrucomicrobia* orders, relative abundances and their PCoA plot (b) based Euclidean model. The nucleotide composition in reactor 3, 6, 7 and 11 partially clustered, lower left quadrant; those in reactor 1 and 10 partially clustered, upper right quadrant of the plot while the reads in reactor 5 and 8 clustered in the upper right quadrant of the plot.
