## Supplementary Fig. 30 for "Metagenomics survey unravels diversities of biogas’ microbiomes with potential to enhance its’ productivity in Kenya"

a

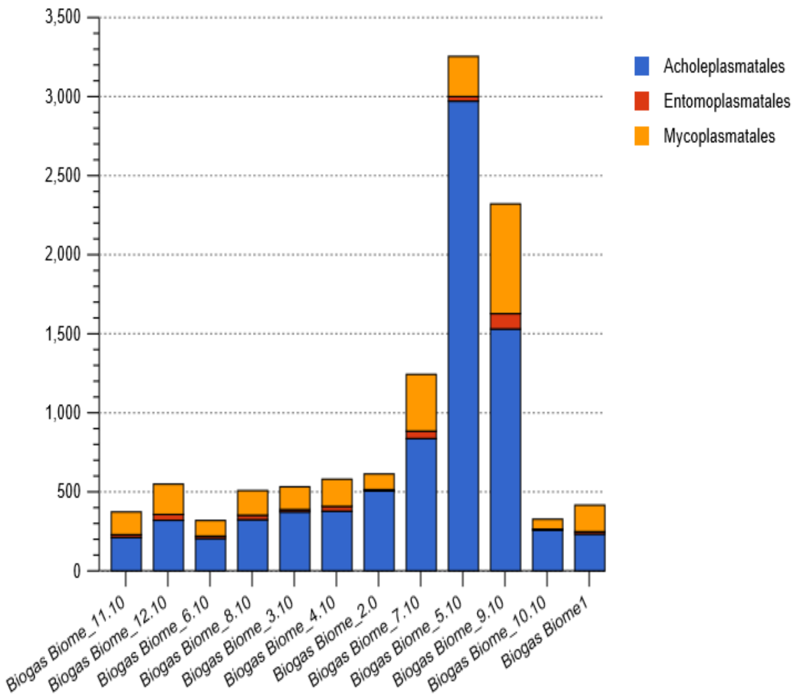


b

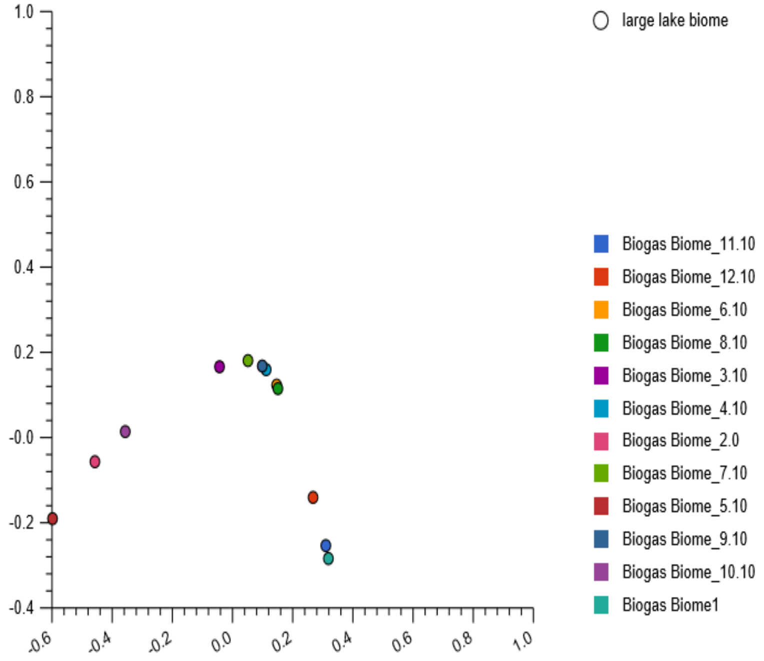


**SFig. 30:** Stacked barchat (a) showing three *Terneicutes* orders, relative abundances and their PCoA plot (b) based on the Euclidean model. The nucleotide composition in reactor 4 and 9 and those detected in reactor 6 and 8 partially clustered in the upper right quadrant of the plot. The other reactors comprised dissimilar nucleotide composition.
