## Supplementary Fig. 31 for "Metagenomics survey unravels diversities of biogas’ microbiomes with potential to enhance its’ productivity in Kenya"

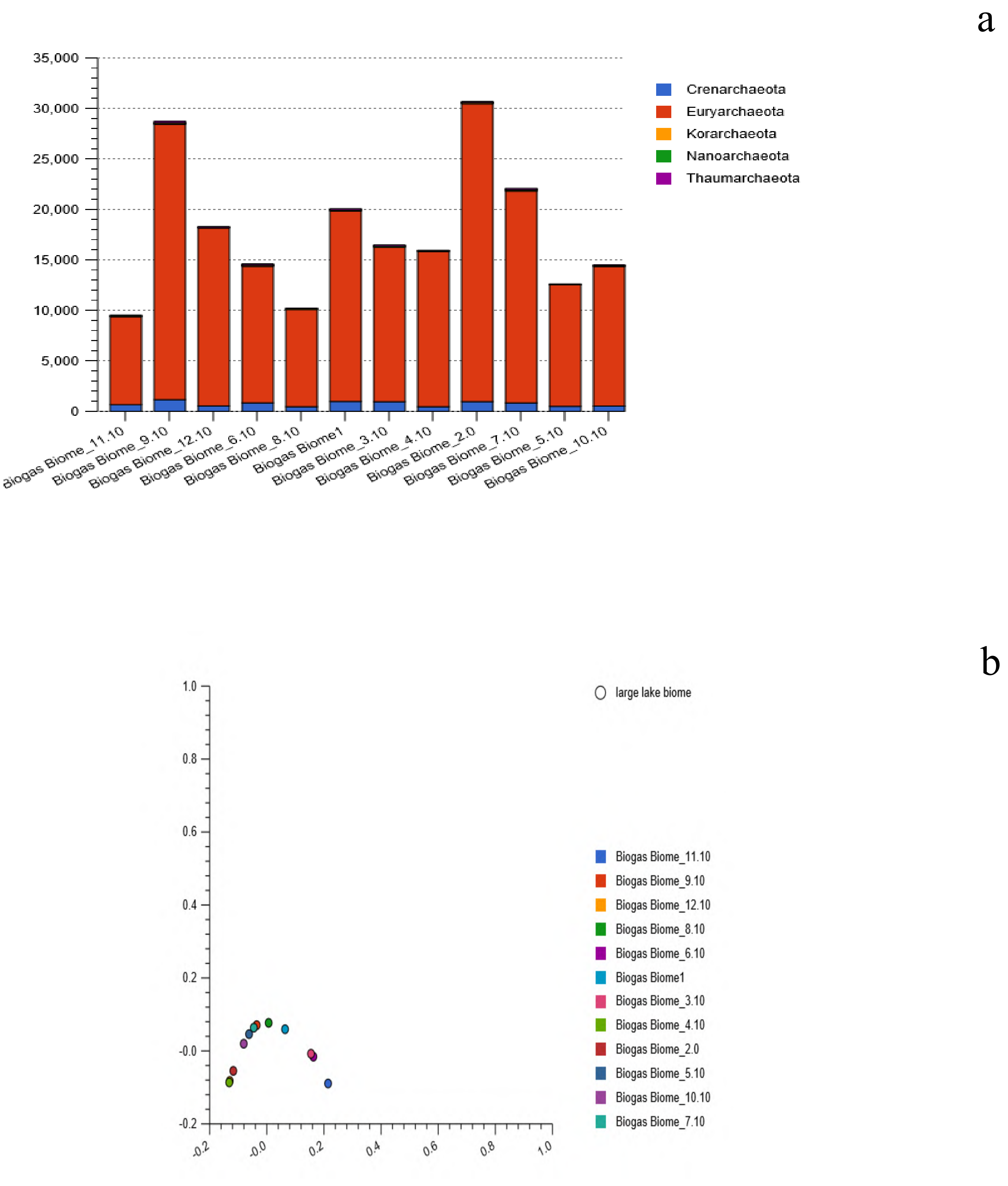


**SFig. 31:** The stacked barchart (a) showing the four Archaeal phyla, relative abundances and their PCoA plot (b) based on the Euclidean model. The nucleotide composition in reactor 4 and 12 clustered, in the lower left quadrant; while those in reactor 3 and 6 and reactor 7 and 9 partially clustered; lower right quadrant and upper left quadrant of the plot. The nucleotide composition in the
