## Supplementary Fig. 32 for "Metagenomics survey unravels diversities of biogas’ microbiomes with potential to enhance its’ productivity in Kenya"

a

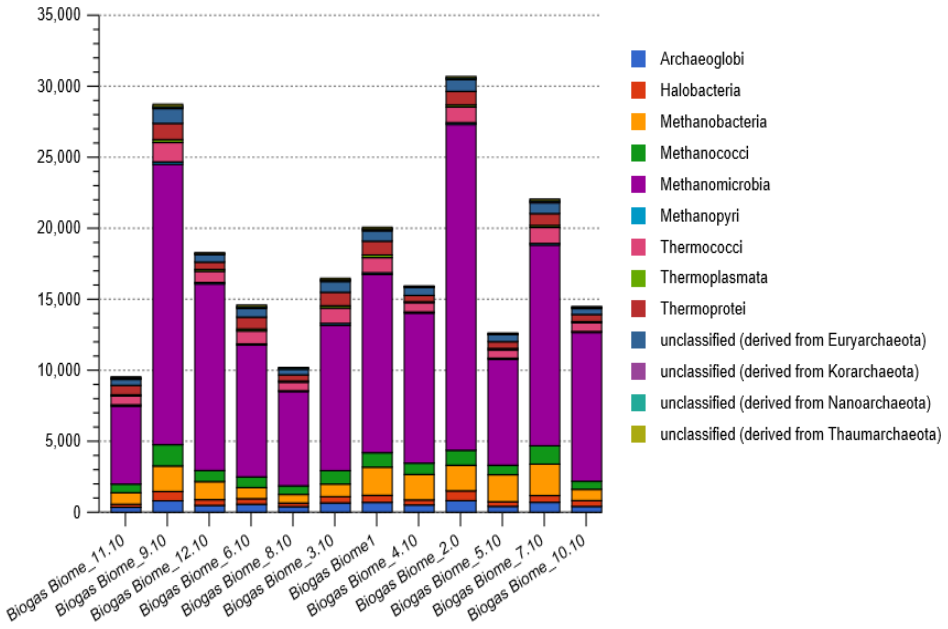


b

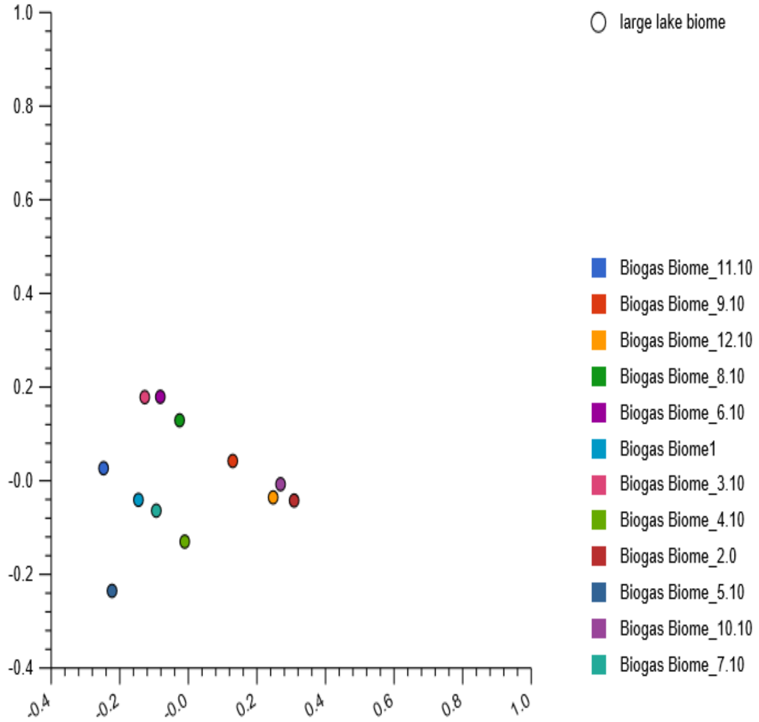


**SFig. 32:** Stacked barchat (a) showing nine archaea classes, and the affiliates of the unclassified reads, relative abundances and their PCoA plot (b) based on the Euclidean model. The nucleotide composition in the respective reactors were distinctively dissimilar.
