## Supplementary Fig. 33 for "Metagenomics survey unravels diversities of biogas’ microbiomes with potential to enhance its’ productivity in Kenya"

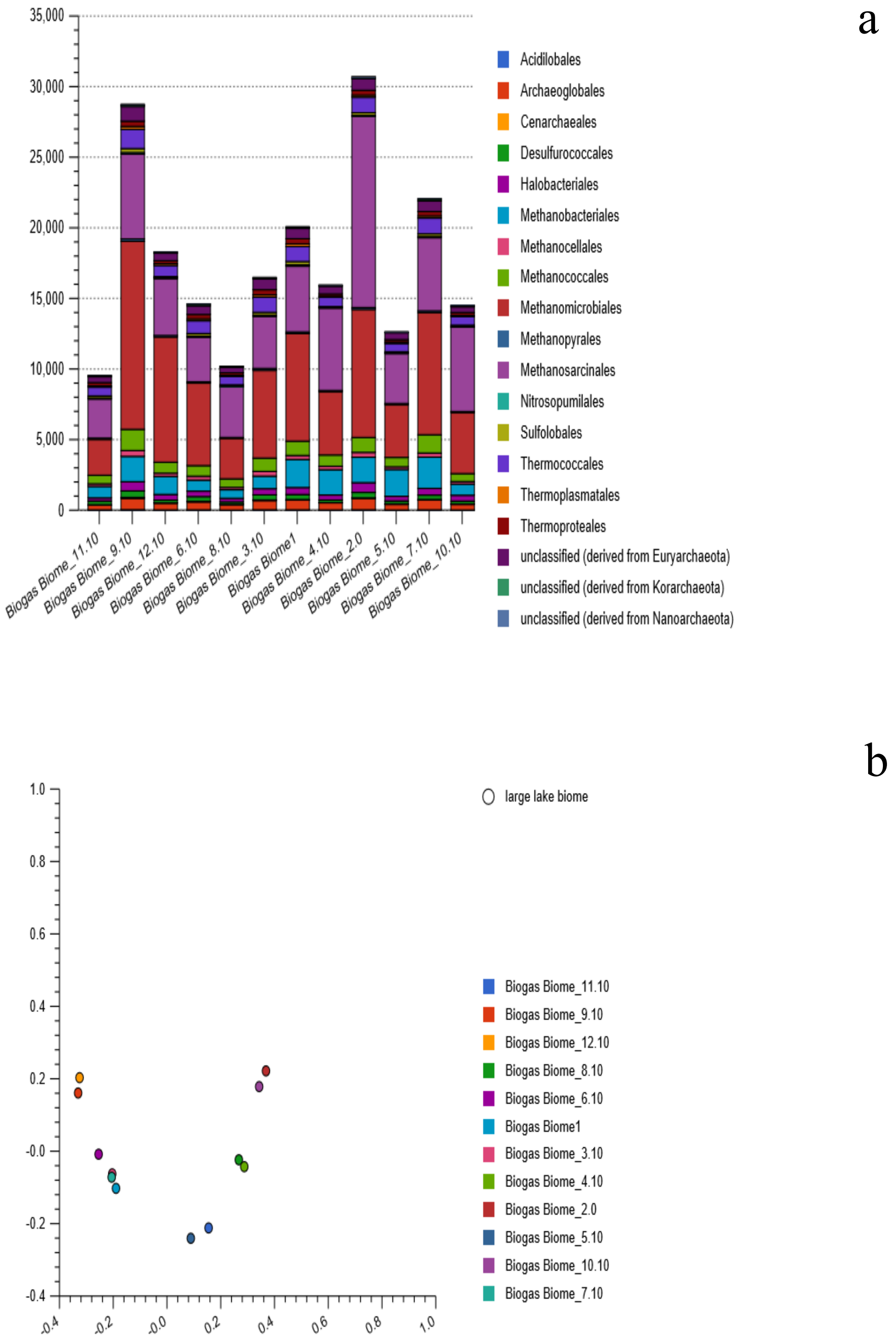


**SFig. 33:** Stacked barchat showing sixteen Archaeal orders, relative abundances and their PCoA plot based on the Euclidean model. The plot revealed partial clustering of the nucleotide in reactor 3 and 7, at lower left quadrant. The nucleotide
