## Supplementary Fig. 34 for "Metagenomics survey unravels diversities of biogas’ microbiomes with potential to enhance its’ productivity in Kenya"

a

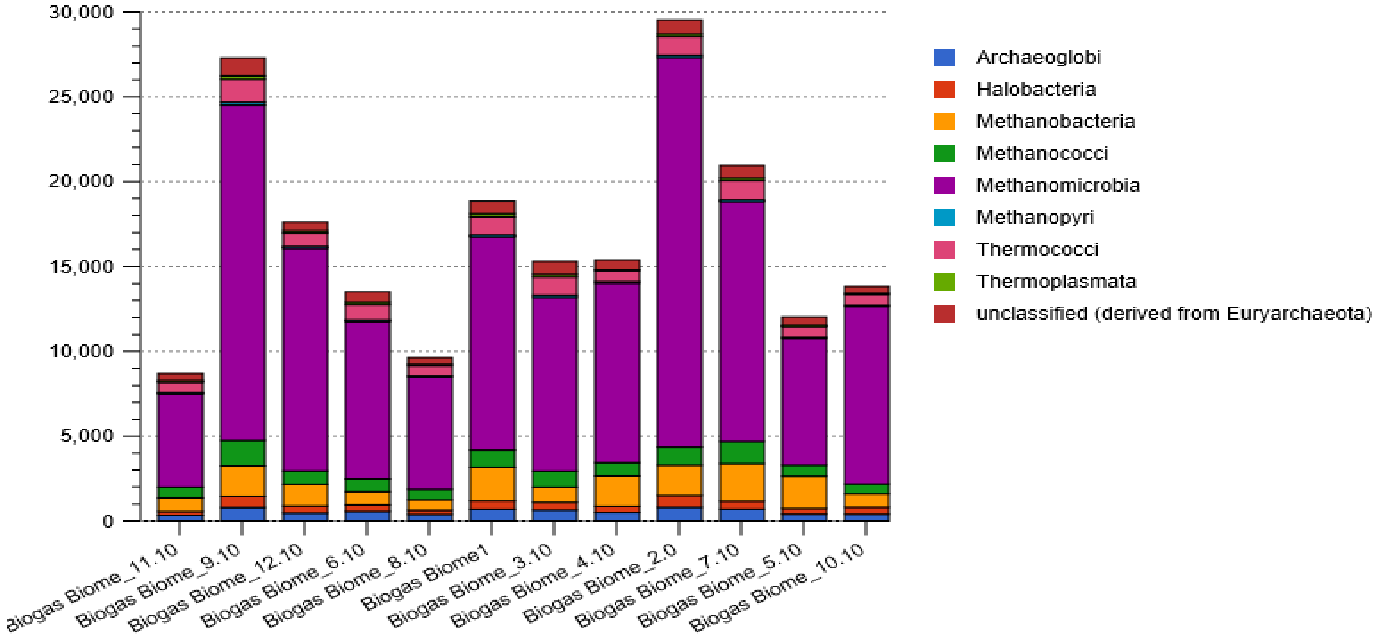


b

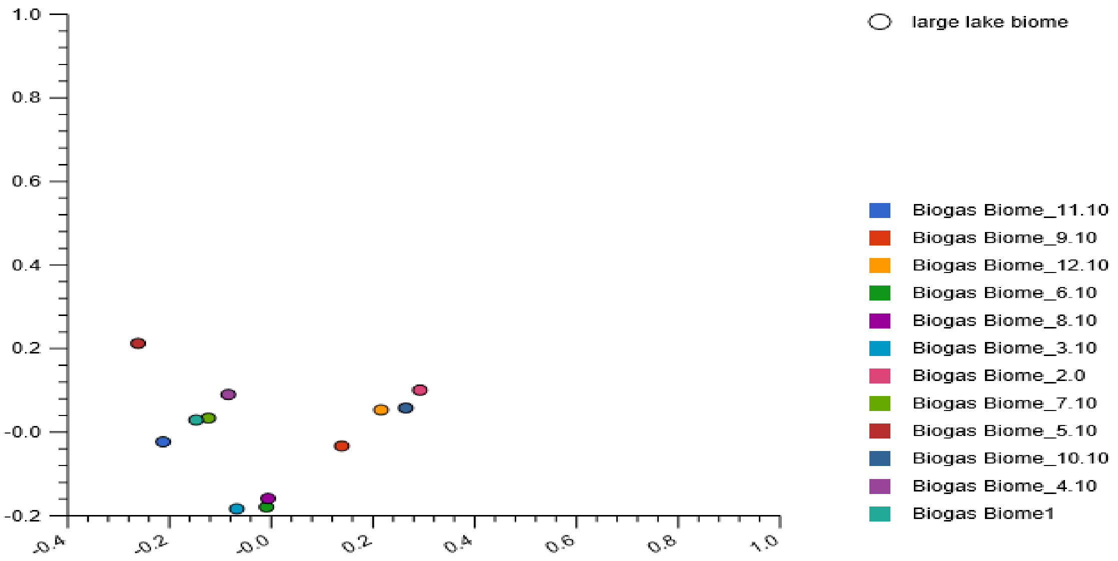


**SFig. 34:** The stacked barchat (a) showing the eight *Eurychaeota* classes, the proportion of relative abundances and their PCoA plots (b) revealing dissimilarities of the nucleotide composition among the twelve studied reactors. The plot revealed the nucleotide composition in reactor 1 and 7, located in the upper left quadrant and those in reactor 6 and 8, located in the lower left quadrant of the plot, near the y-axis were in close proximity.
