## Supplementary Fig. 35 for "Metagenomics survey unravels diversities of biogas’ microbiomes with potential to enhance its’ productivity in Kenya"

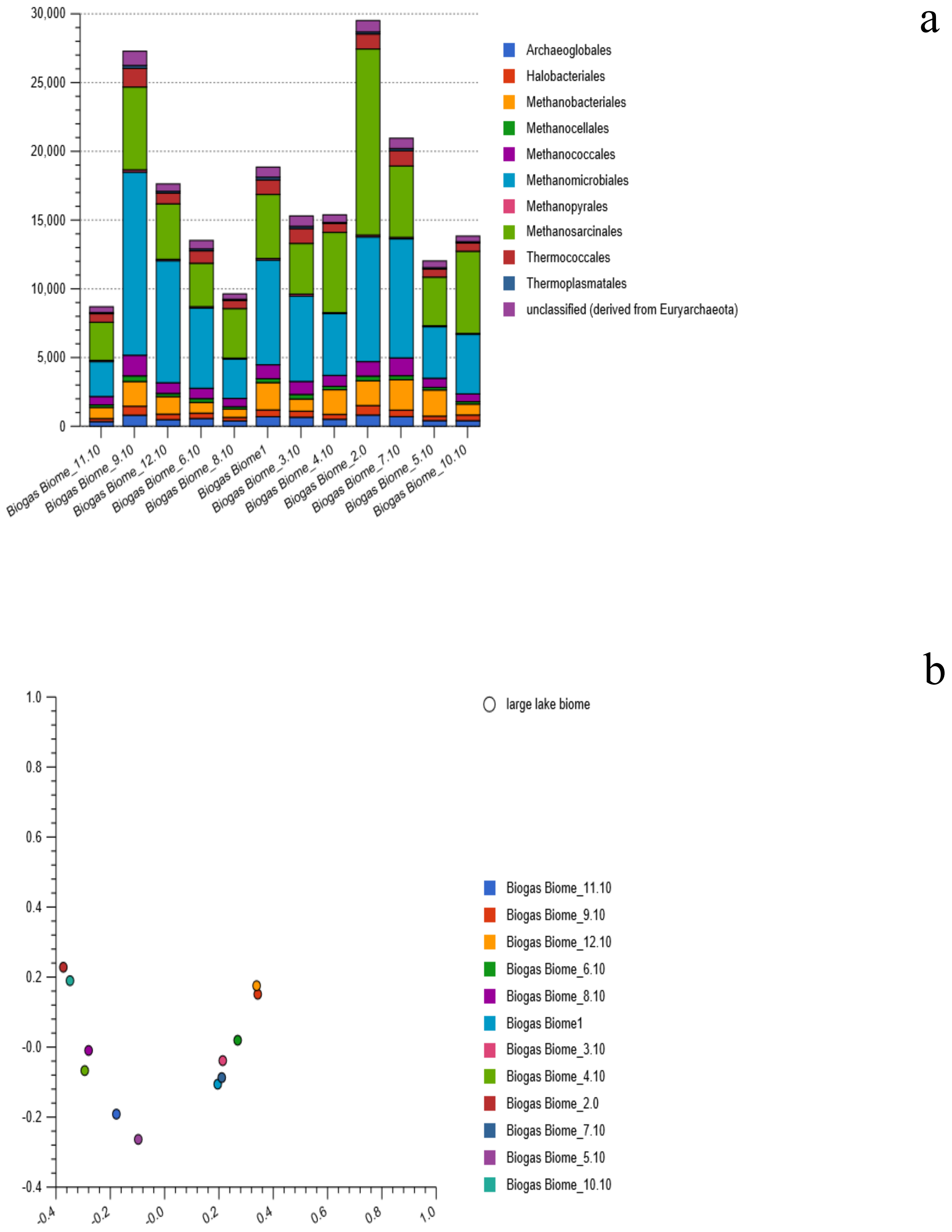


**SFig. 35:** Stacked barchat (a) showing ten *Euryarchaeota* orders, relative abundances and their PCoA plot (b) based Euclidean model. Their nucleotide composition were dissimilar, distributed in the four quadrant.
