## Supplementary Fig. 36 for "Metagenomics survey unravels diversities of biogas’ microbiomes with potential to enhance its’ productivity in Kenya"

a

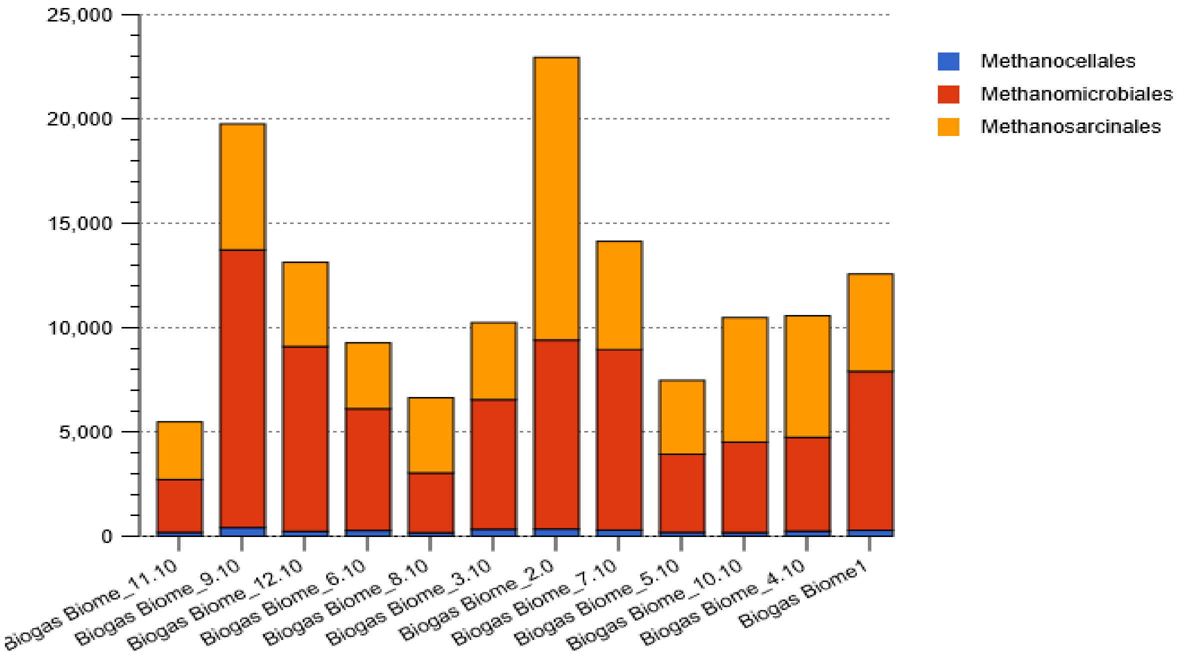


b

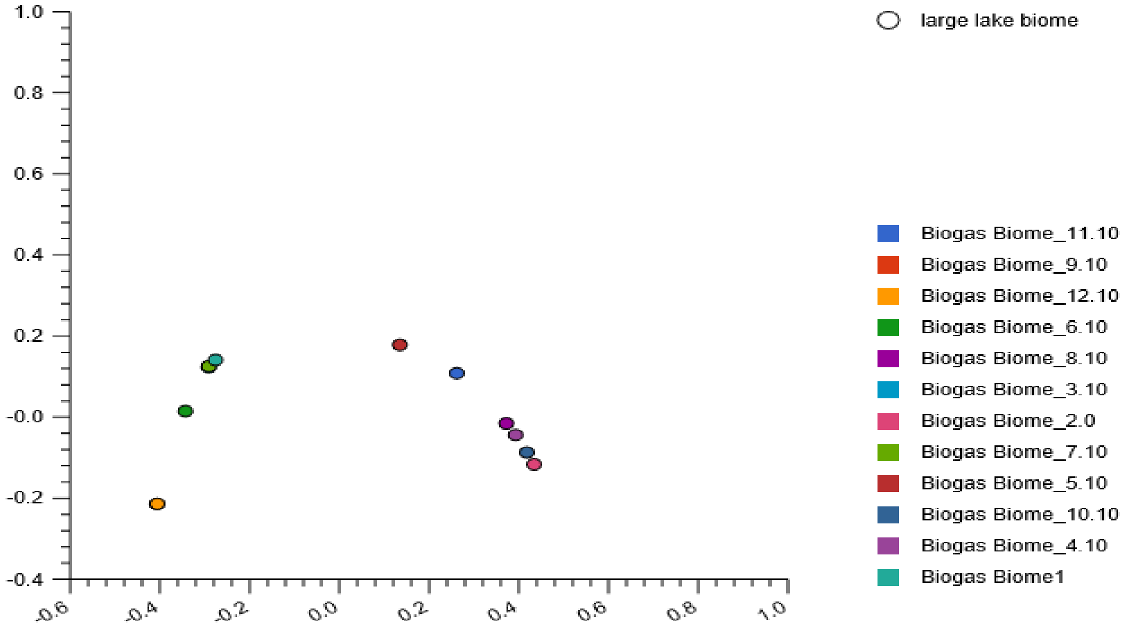


**Fig. 36:** The stacked barchat (a) showing three Methanomicrobia orders, the proportion of relative abundances and their PCoA plot (b) based on the Euclidean model. The PCoA plot revealed that the nucleotide composition in reactor 1 and 7 clustered partially on the upper left of the plot. Further the nucleotide composition in reactor 2 and 10 and those in reactor 4 and 8 were in close prox-
