## Supplementary Fig. 37 for "Metagenomics survey unravels diversities of biogas’ microbiomes with potential to enhance its’ productivity in Kenya"

a

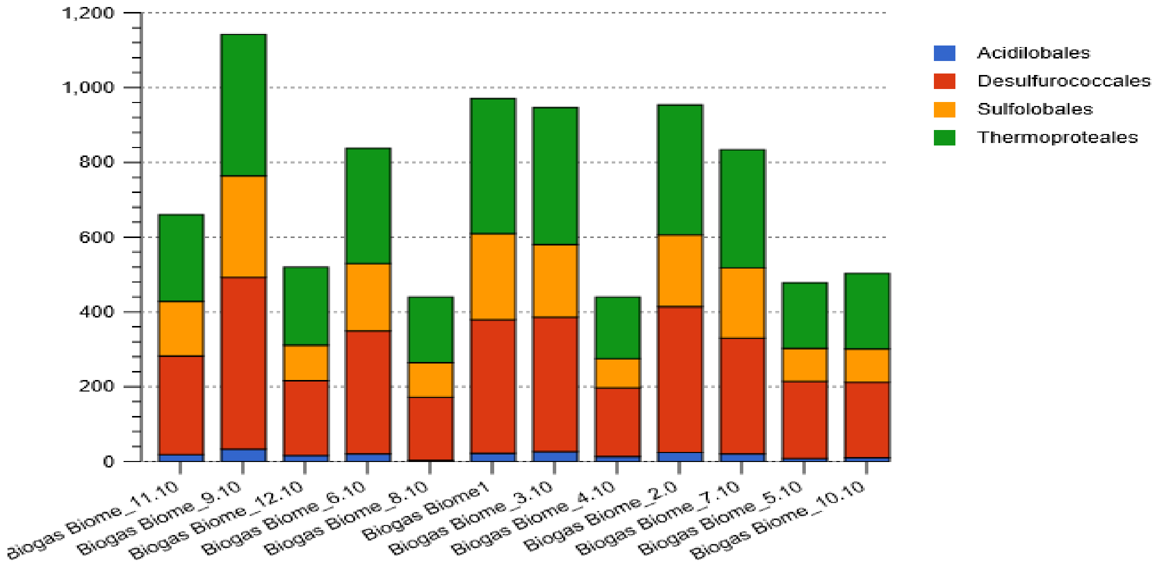


b

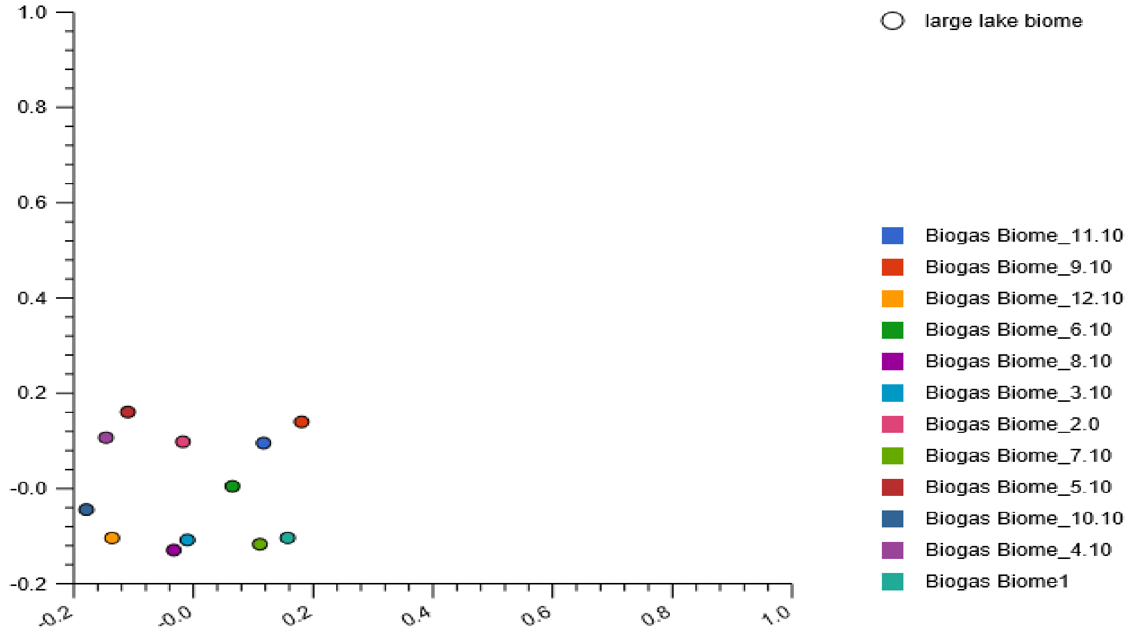


**SFig. 37:** Stacked barchat (a) showing four Thermoprotei orders and their PCoA plot (b) based on the Euclidean model. The plot reveal nucleotide composition dissimilarities among the twelve reactors. However, the nucleotide composition in reactor 3 and 8 were closely located in the lower left quadrant of the plot.
