## Supplementary Fig. 38 for "Metagenomics survey unravels diversities of biogas’ microbiomes with potential to enhance its’ productivity in Kenya"

a

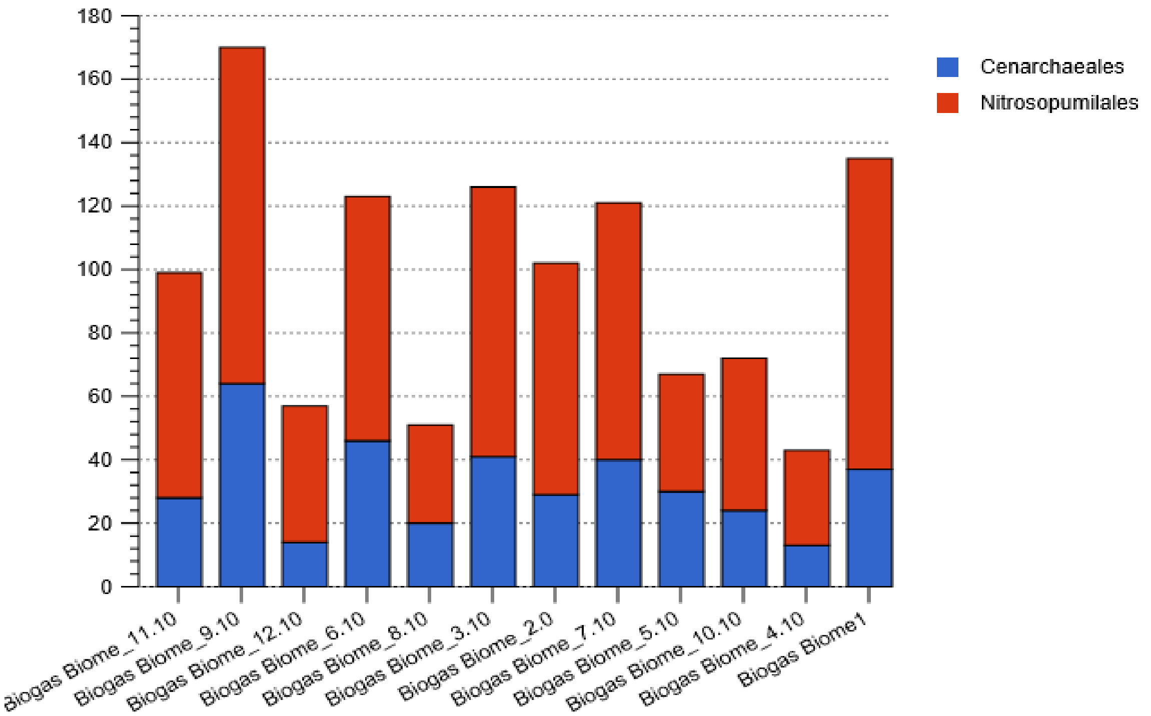


b

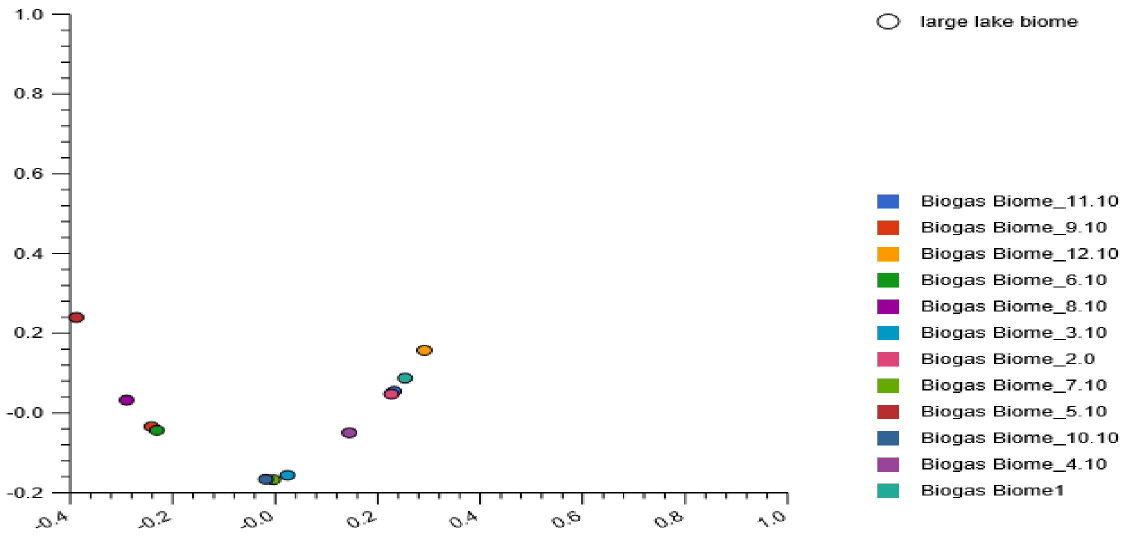


**SFig. 38:** Stacked barchat (a) showing two *Thaurmarchaeota* orders, the relative abundances, and their PCoA plot (b) based on the Euclidean model. The plot revealed partial clustering of the nucleotide composition in reactor 6 and 9, located in the lower left quadrant of the plot, reactor 7 and 10, located near the y-axis, and reactor 2 and 11 located on the upper right quadrant of the plot. The nucleotide composition in reactor 3 were in close proximity to those detected in reactor 7, located on the lower right quadrant of the plot. The nucleotide composition in the rest of the reactors were dissimilar.
