## Supplementary Fig. 39 for "Metagenomics survey unravels diversities of biogas’ microbiomes with potential to enhance its’ productivity in Kenya"

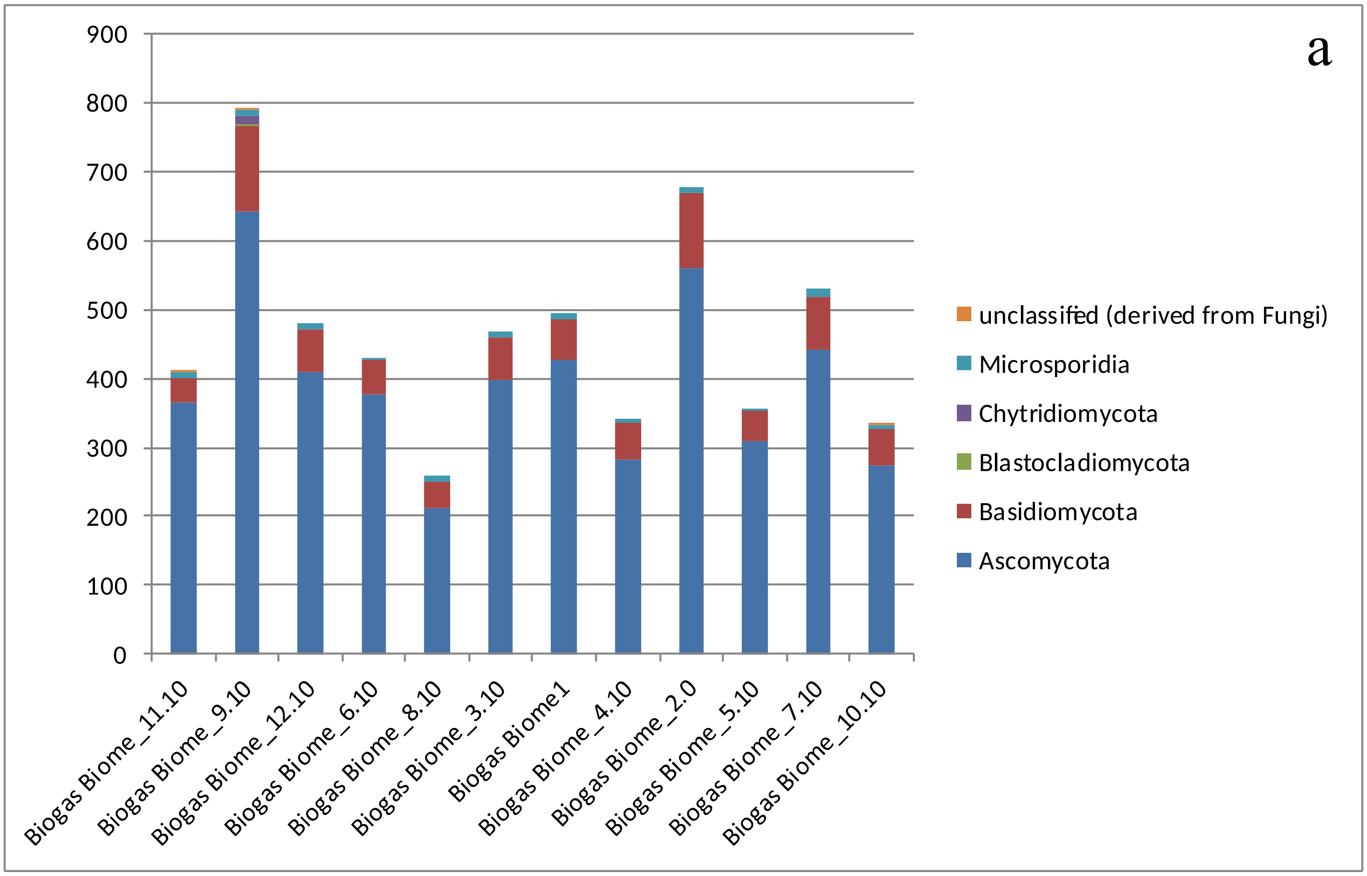


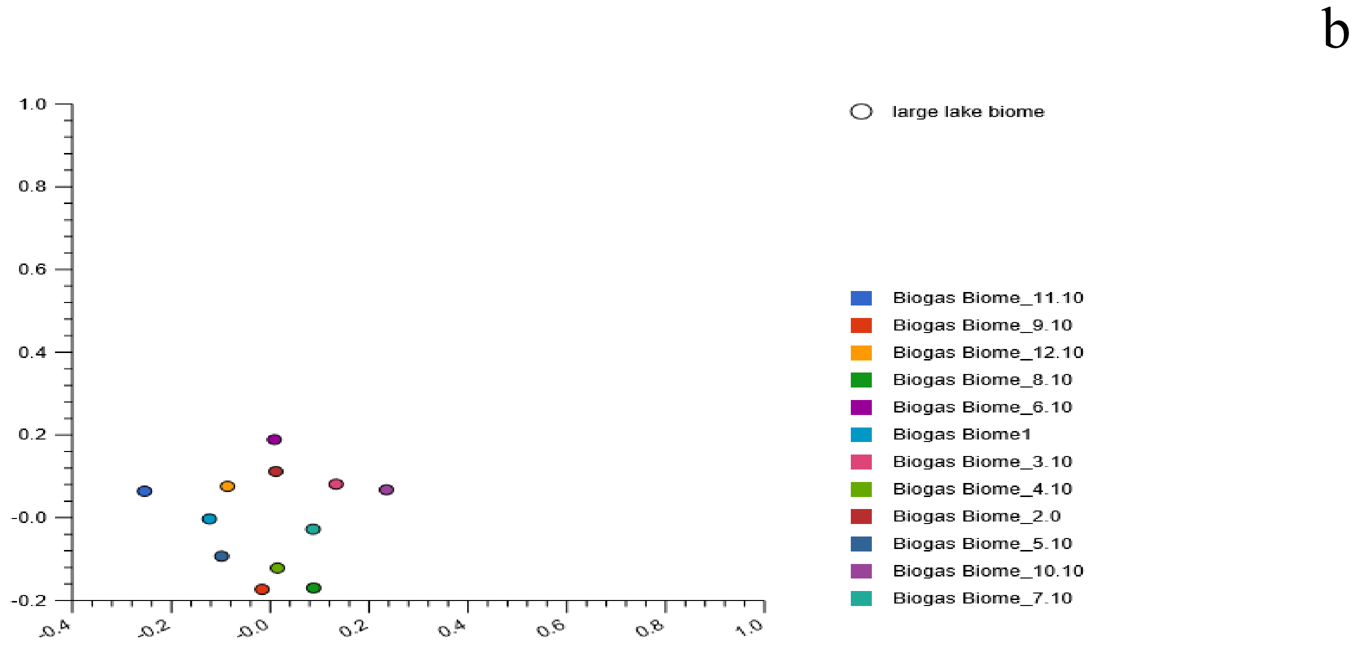


**SFig. 39:** Stacked barchat (a) showing five Fungal phyla, proportion of the relative abundances and their PCoA plots (b) based on Euclidean model. The plot revealed dissimilarities of the nucleotide composition among the studied reactors. distributed within the four plot quadrant.
