## Supplementary Fig. 40 for "Metagenomics survey unravels diversities of biogas’ microbiomes with potential to enhance its’ productivity in Kenya"

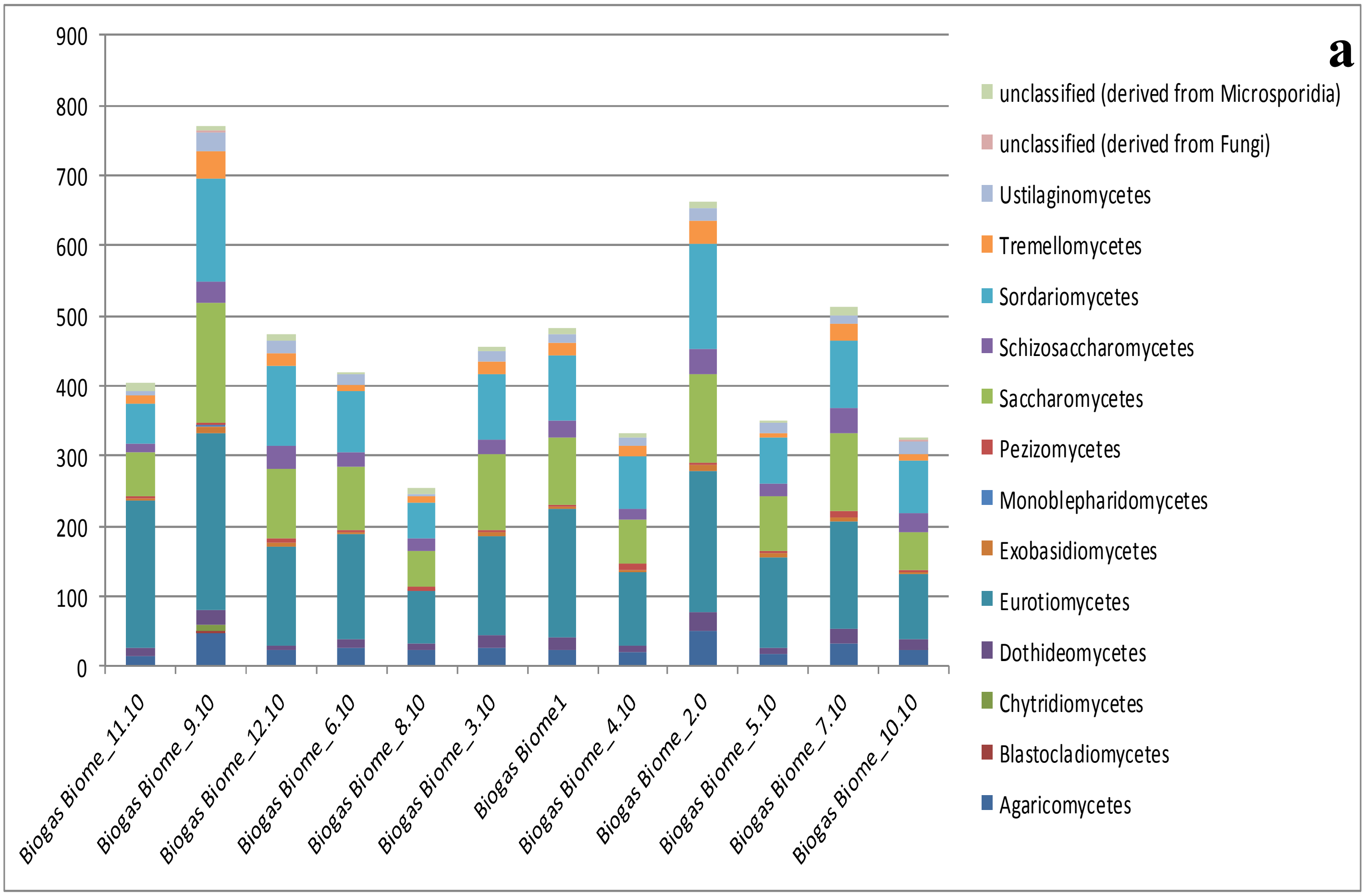


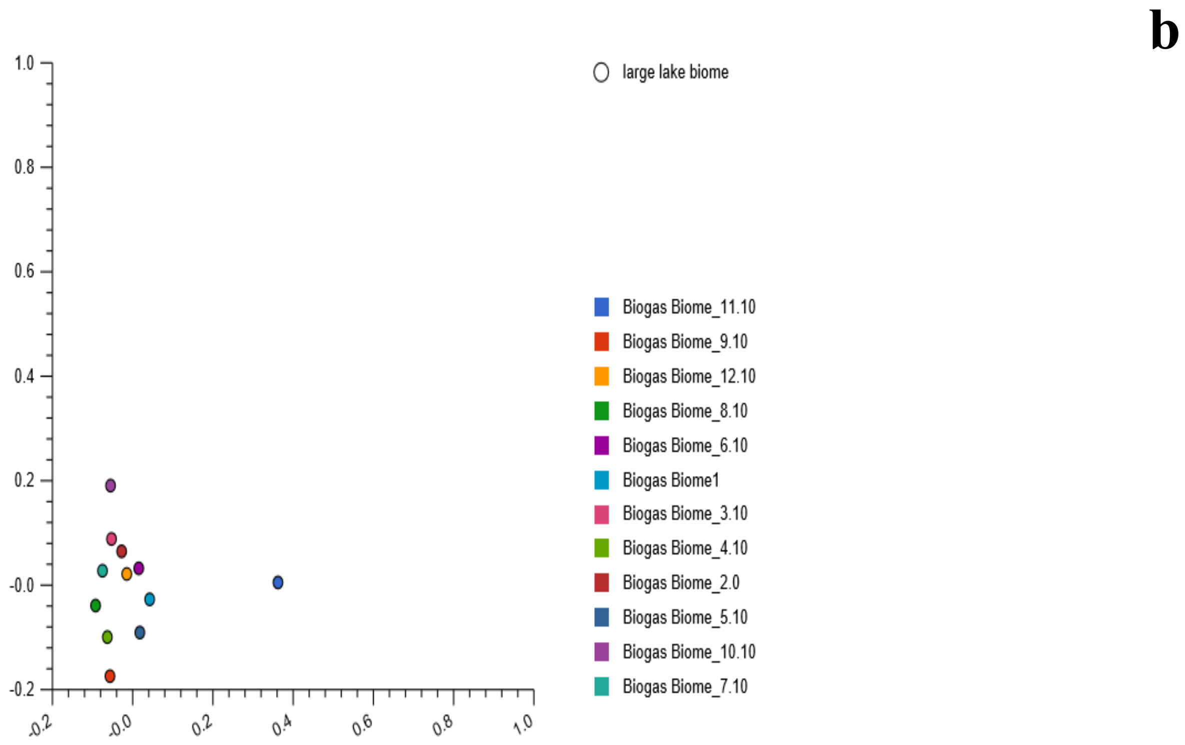


**SFig. 40:** Stacked barchat (a) showing thirteen eukaryotes classes, relative abundances and their PCoA plot (b) based on the Euclidean model. The PCoA plot revealed nucleotide composition dissimilarities among the studied twelve reactors.
