## Supplementary Fig. 41 for "Metagenomics survey unravels diversities of biogas’ microbiomes with potential to enhance its’ productivity in Kenya"

a

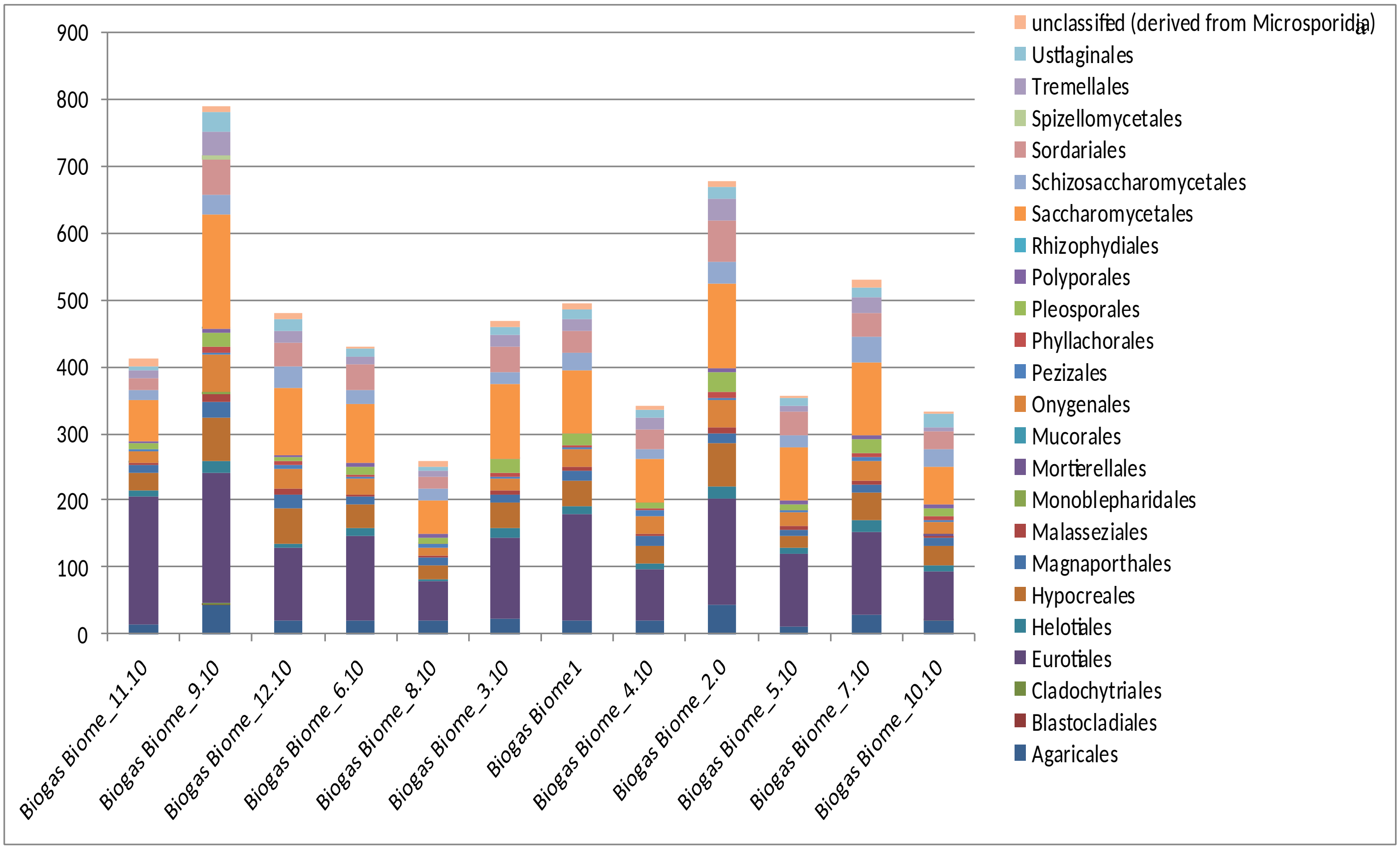


b

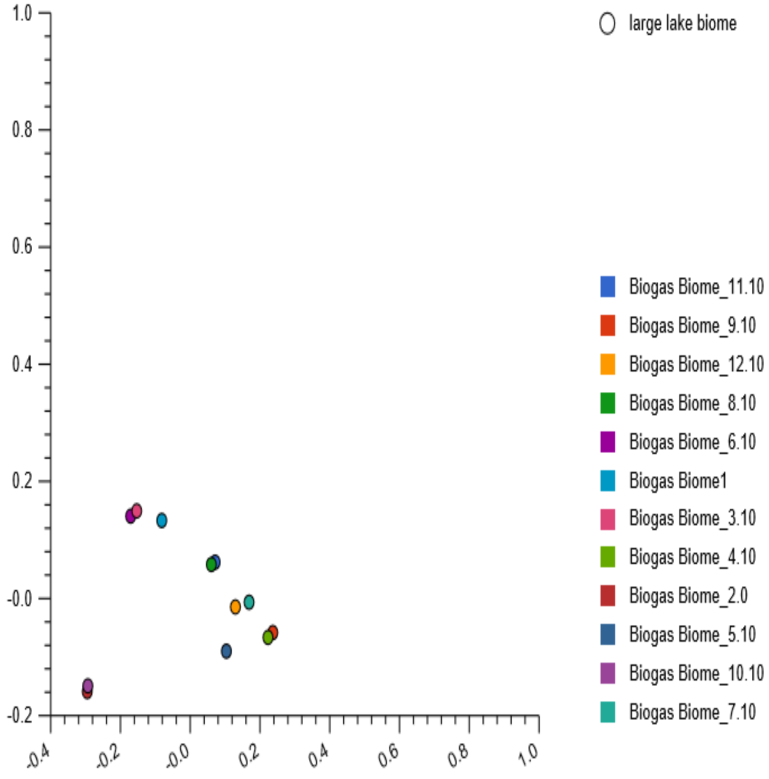


**SFig. 41:** Stacked barchat (a) showing 23 fungal orders, relative abundances and their PCoA plot (b) based on the Euclidean model. The nucleotide composition for reactor 2 and 10, lower left quadrant; reactor 3 and 6, upper left quadrant; reactor 4 and 9, upper right quadrant; and reactor 8 and 11, lower right quadrant of the plot, partially clustered.
