## Supplementary Fig. 42 for "Metagenomics survey unravels diversities of biogas’ microbiomes with potential to enhance its’ productivity in Kenya"

a

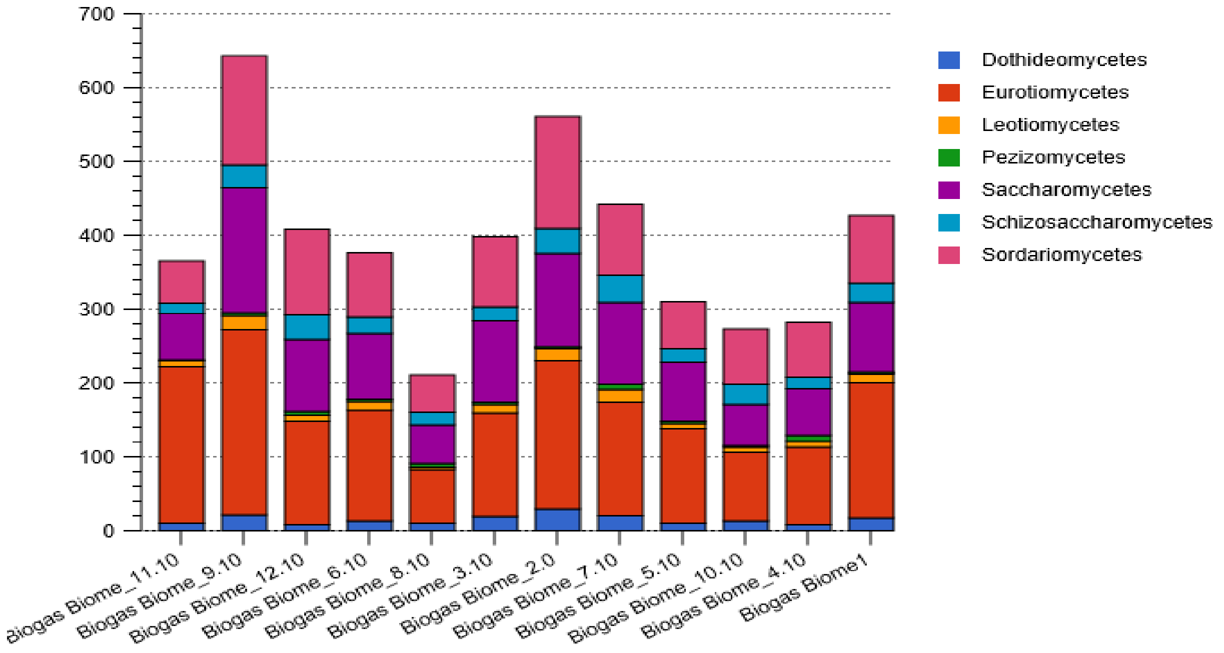


b

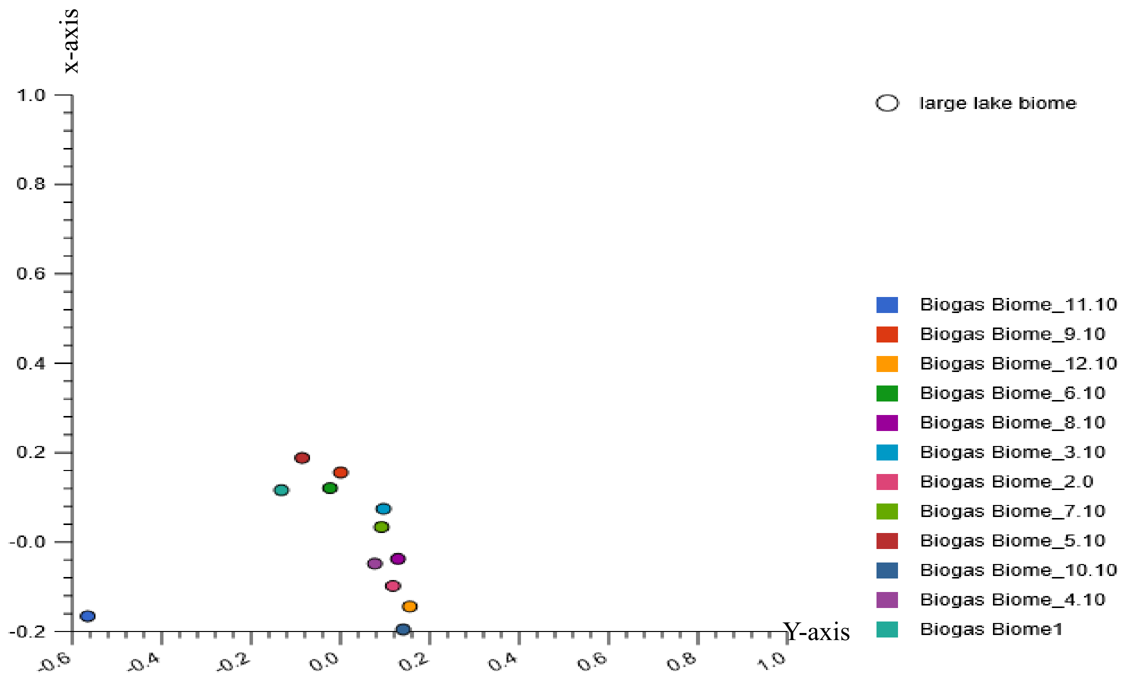


**SFig. 42:** The stacked barchat (a) showing the five Ascomycota classes, relative abaundances and their PCoA plots (b), based on the Euclidean model at the class level. The PCoA plots revealed dissimilar nucleotide composition among the twelve reactors. However, reactor 11 was located singly in the lower left quandrant of the plot while the nucleotide in reactor 10 were located along the y-axis (Negative PCoA 1 and Negative PCoA 2).
