## Supplementary Fig. 43 for "Metagenomics survey unravels diversities of biogas’ microbiomes with potential to enhance its’ productivity in Kenya"

a

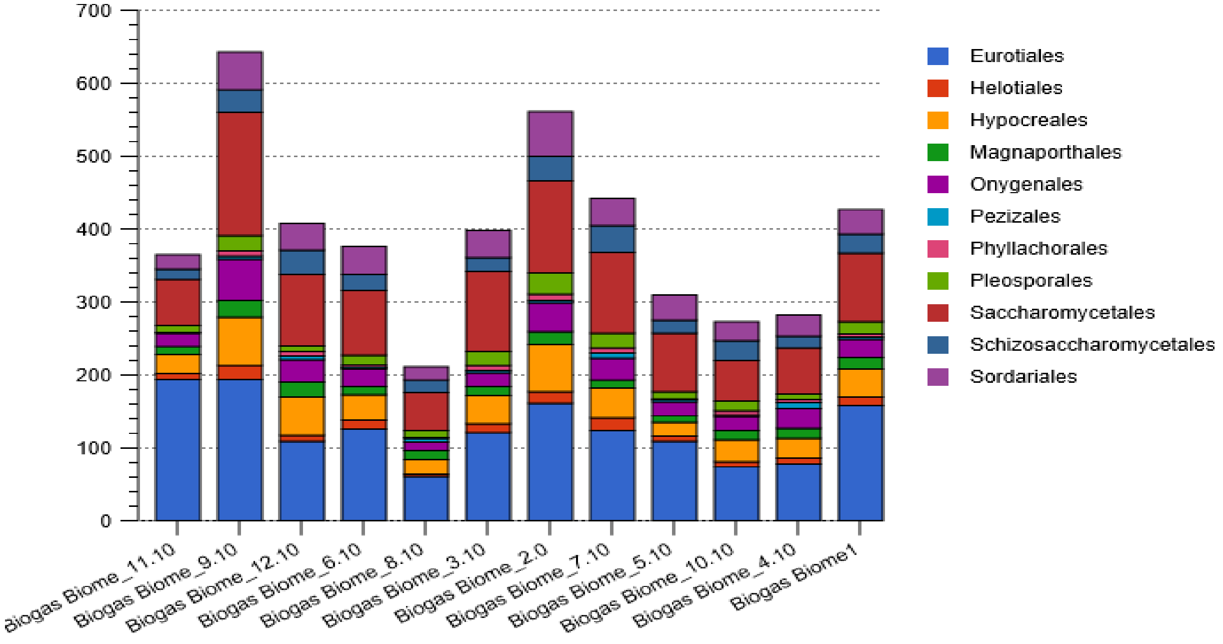


b

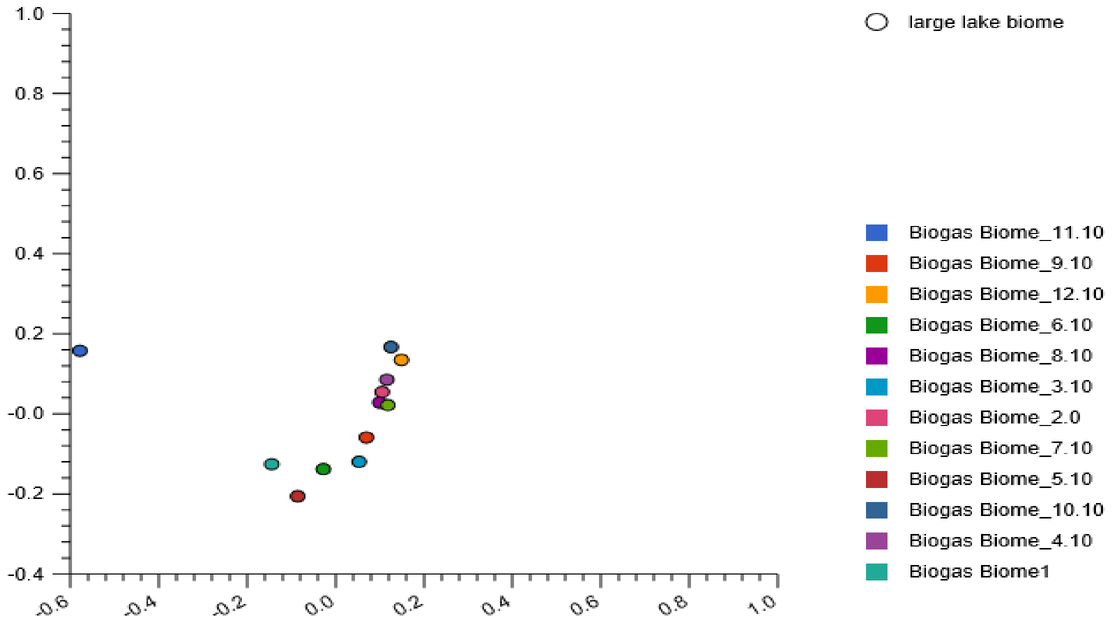


**SFig. 43:** The stacked barchat (a) revealing eleven Ascomycota orders, proportions of relative abundances and their PCoA plot (b); based on the Euclidean model revealed dissimilarities among the 12 reactors. However, the model revealed that the nucleotide composition in reactor 7 and 8 partially clustered while those in reactor 2 and 8 and reactor 2 and 4 were in close proximity, located in the upper right quadrant of the plot. The *Ascomycota*'s nucleotide composition in reactor 11 were singly located in the
