## Supplementary Fig. 44 for "Metagenomics survey unravels diversities of biogas’ microbiomes with potential to enhance its’ productivity in Kenya"

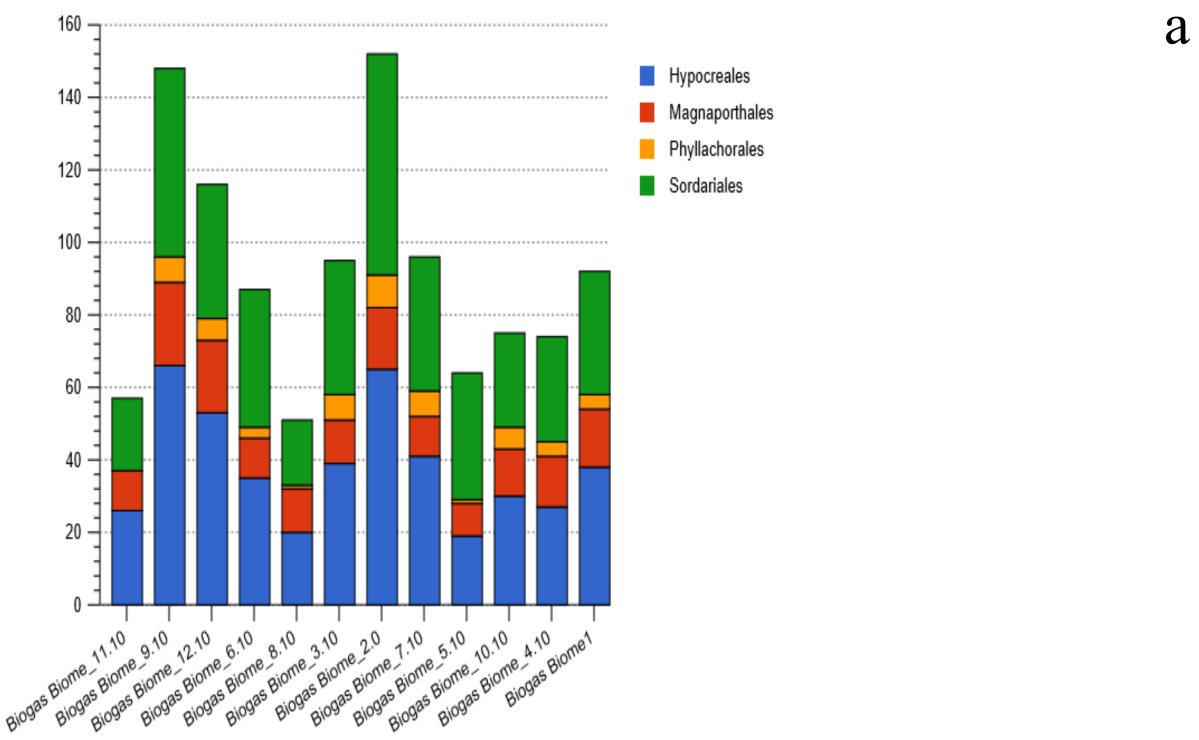


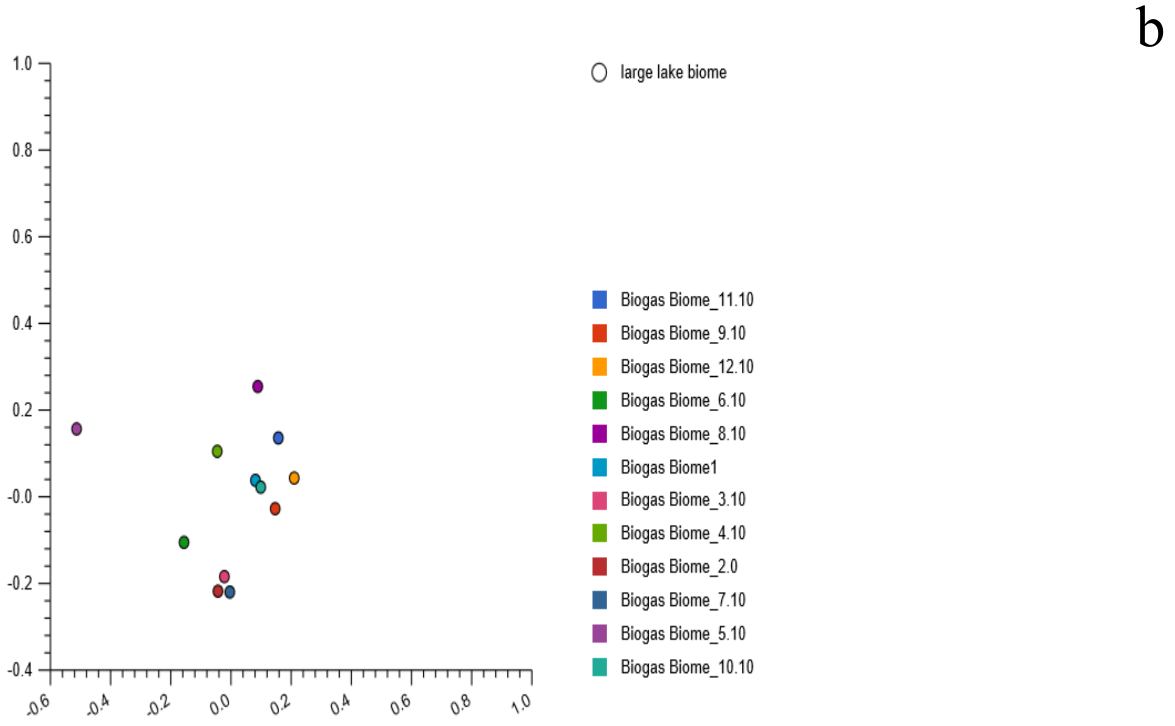


**SFig. 44:** Stacked barchat showing three Sordariomycetes orders, relative abundances and their PCoA plot based on the Euclidean model. The PCoA plots revealed partial clustering of the their nucleotide reads in reactor 1 and 10 while in other nucleotide composition in the other reactors were dissimilar.
