## Supplementary Fig. 45 for "Metagenomics survey unravels diversities of biogas’ microbiomes with potential to enhance its’ productivity in Kenya"

a

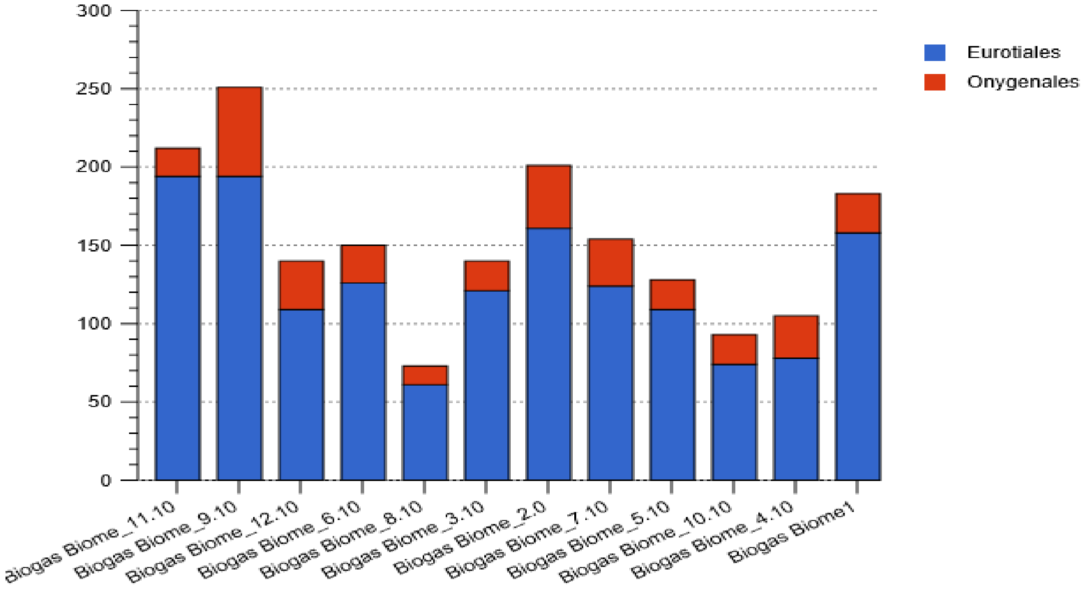


b

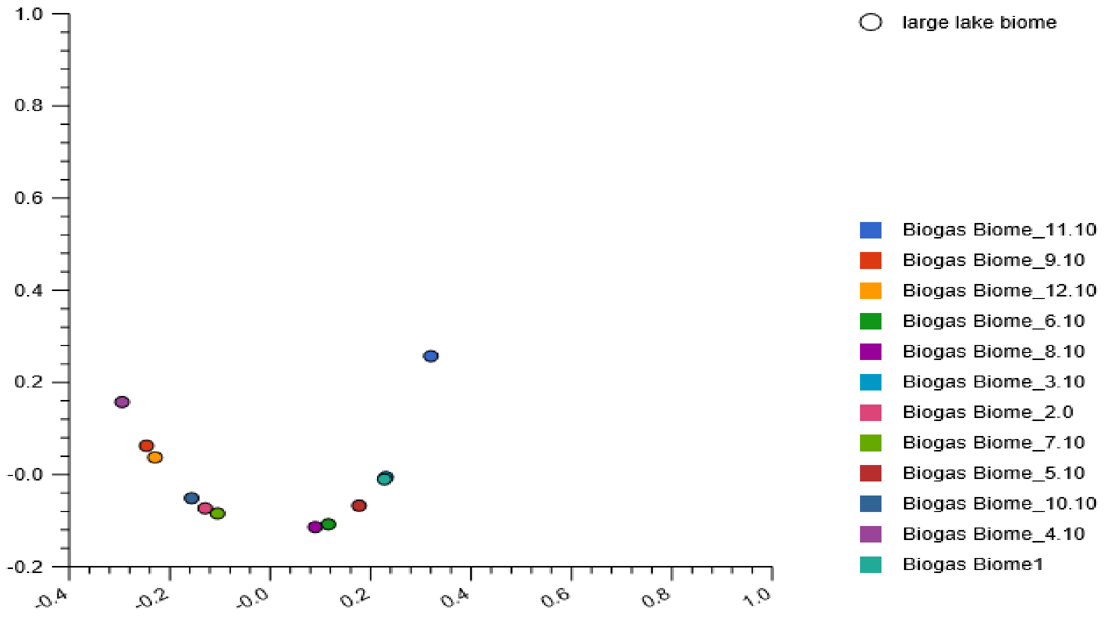


**SFig. 45:** Stacked barchat (a) showing the two *Eurotiomycete*s orders, the proportion of relative abundances and their PCoA plot (b) based on the Euclidean model. The nucleotide composition in reactor 1 and 3 clustered on the x-axis (positive PCoA 1 and Positive PCoA2), while those in reactor 2 and 7, located on the lower left quadrant and reactor 6 and 8, located on the lower right quadrant of the plot were in close proximity. However, the nucleotide composition in reactor 11 were singly located on the upper right quadrant of the plot.
