## Supplementary Fig. 46 for "Metagenomics survey unravels diversities of biogas’ microbiomes with potential to enhance its’ productivity in Kenya"

a

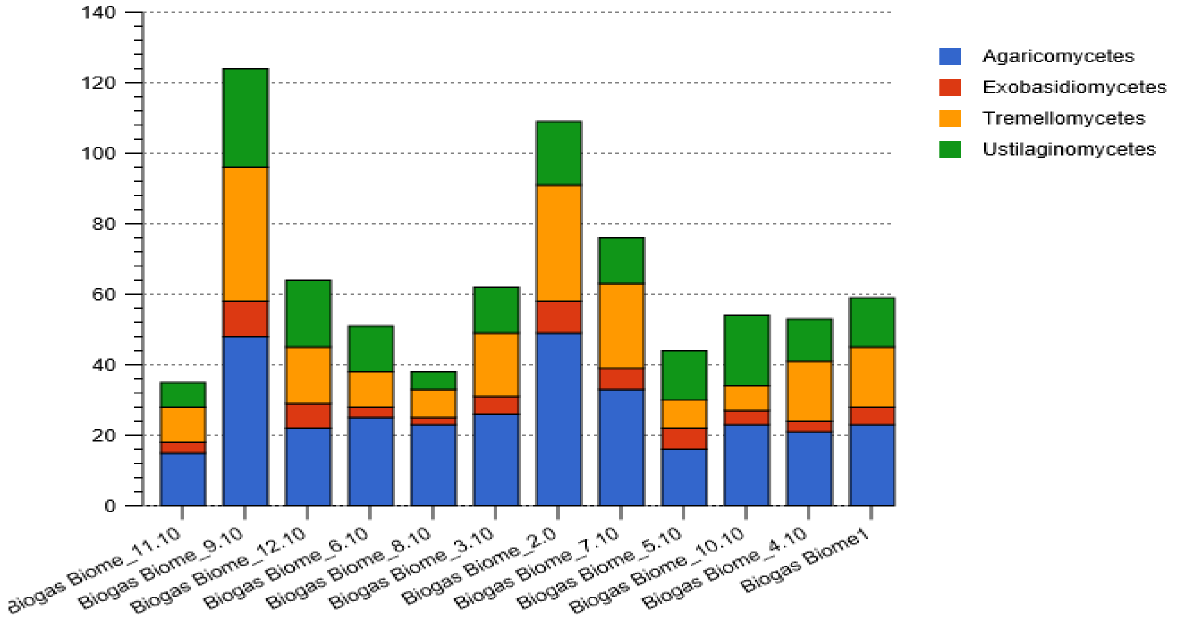


b

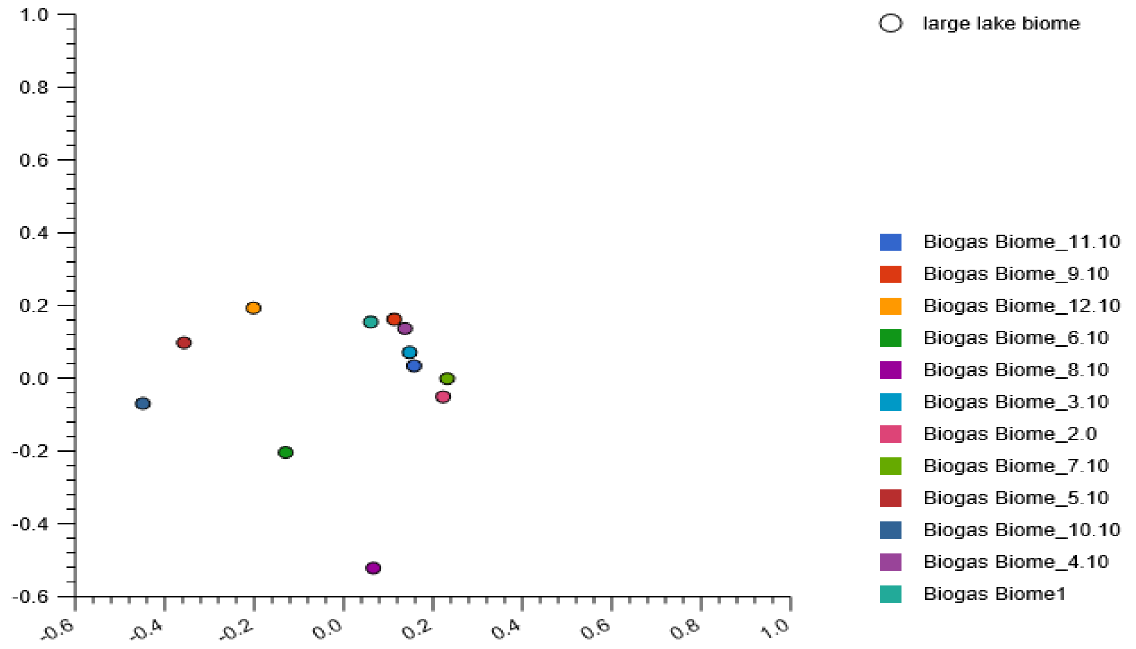


**SFig. 46:** Stacked barchat (a) showing four *Basidiomycota* classes, the proportion of the relative abundances and their PCoA plot (b) based on the Euclidean model. The nucleotide composition in reactor 4 and 9 were in close proximity, located on the upper right quadrant of the plot. The nucleotide composition in reactor 3 and 11 were also closely located in the upper right quadrant, near the x-axis. The rest of the nucleotide composition in other reactors were distributed in the four plot quadrant.
