## Supplementary Fig. 47 for "Metagenomics survey unravels diversities of biogas’ microbiomes with potential to enhance its’ productivity in Kenya"

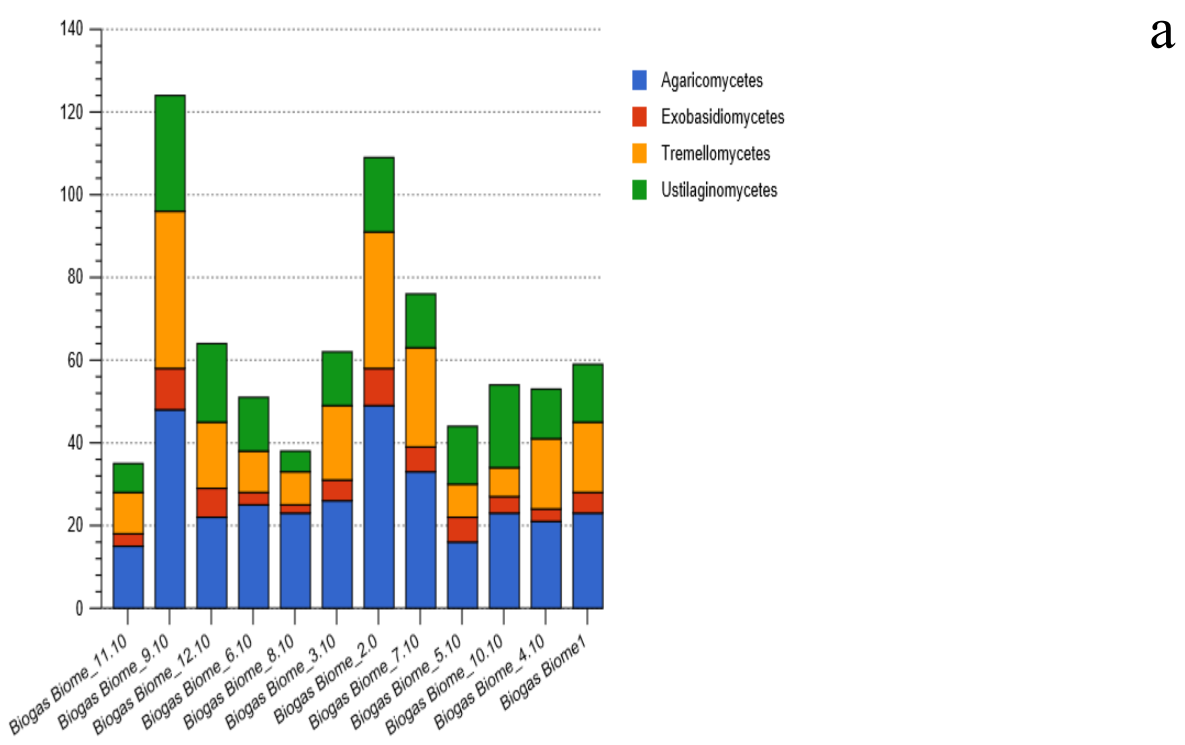


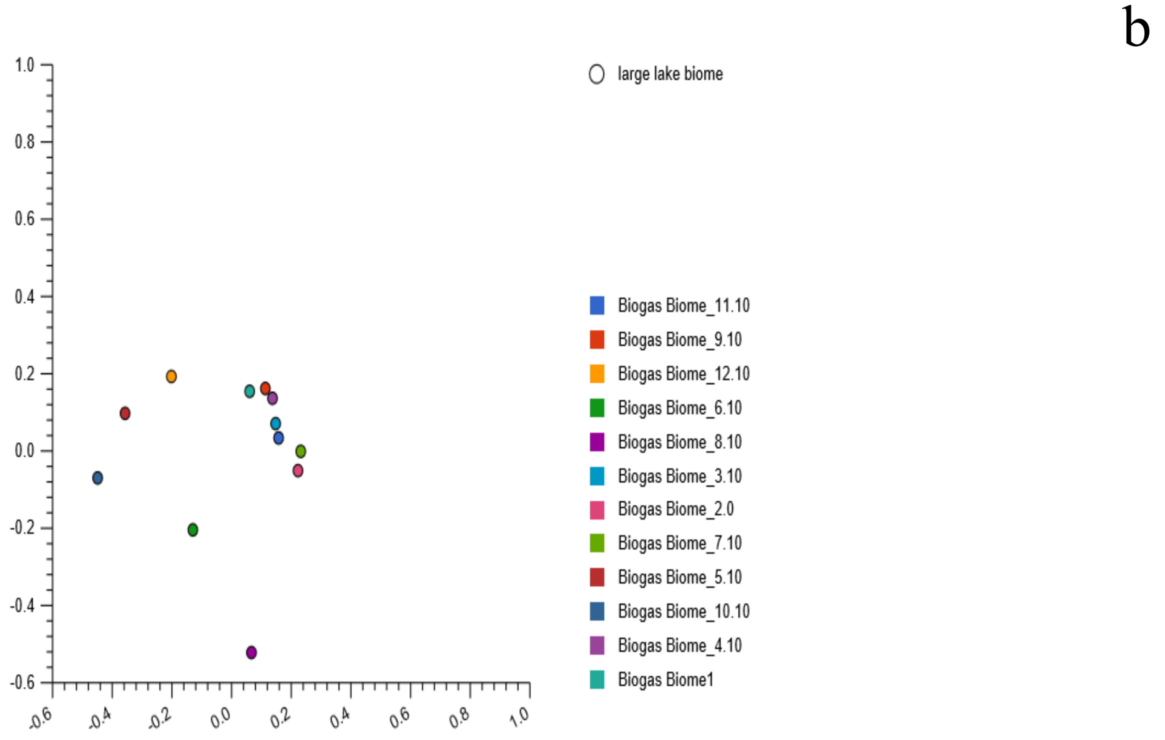


**SFig. 47:** Stacked barchat (a) showing four *Basidiomycota* orders, relative abundances and their PCoA plot (b) based on the Euclidean model. The nucleotide composition in twelve reactors were dissimilar. Located in the four quadrants.
