## Supplementary Fig. 48 for "Metagenomics survey unravels diversities of biogas’ microbiomes with potential to enhance its’ productivity in Kenya"

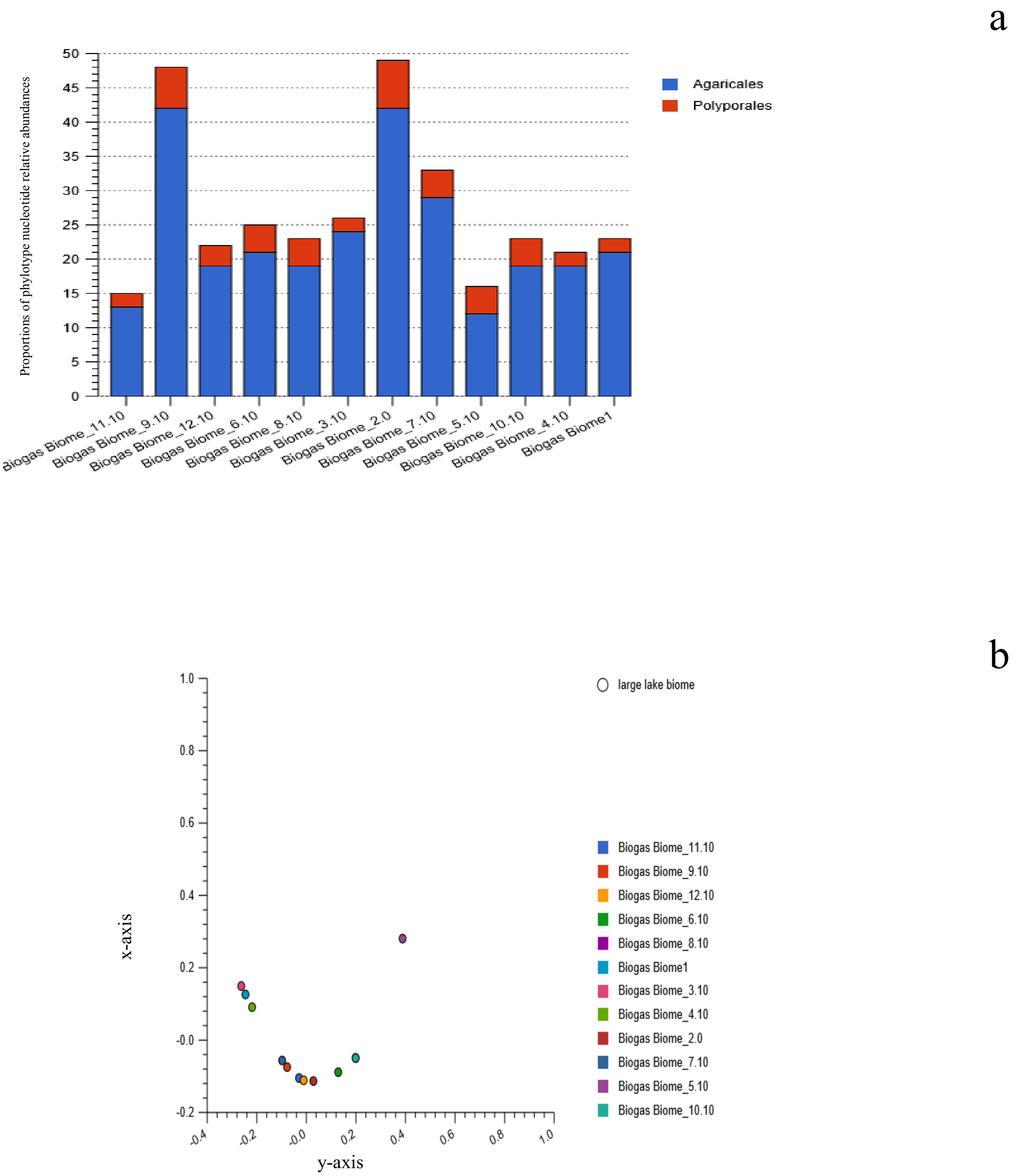


**SFig. 48:** Stacked barchart (a) showing the proportion of the agaricomycetes relative abundances and their PCoA plot (b) based on the Euclidean model at the order level: The nucleotide composition for *agaricomycete*s in reactor 1 and 3 were almost similar, located in the upper left quandrant of the Euclidean plot. Their nucleotide composition in reactor 7 and 9 were in close proximity, located in the lower left quandrant of the plot. The *agaricomycete*s nucleotides in reactor 11 and 12
