## Supplementary Fig. 49 for "Metagenomics survey unravels diversities of biogas’ microbiomes with potential to enhance its’ productivity in Kenya"

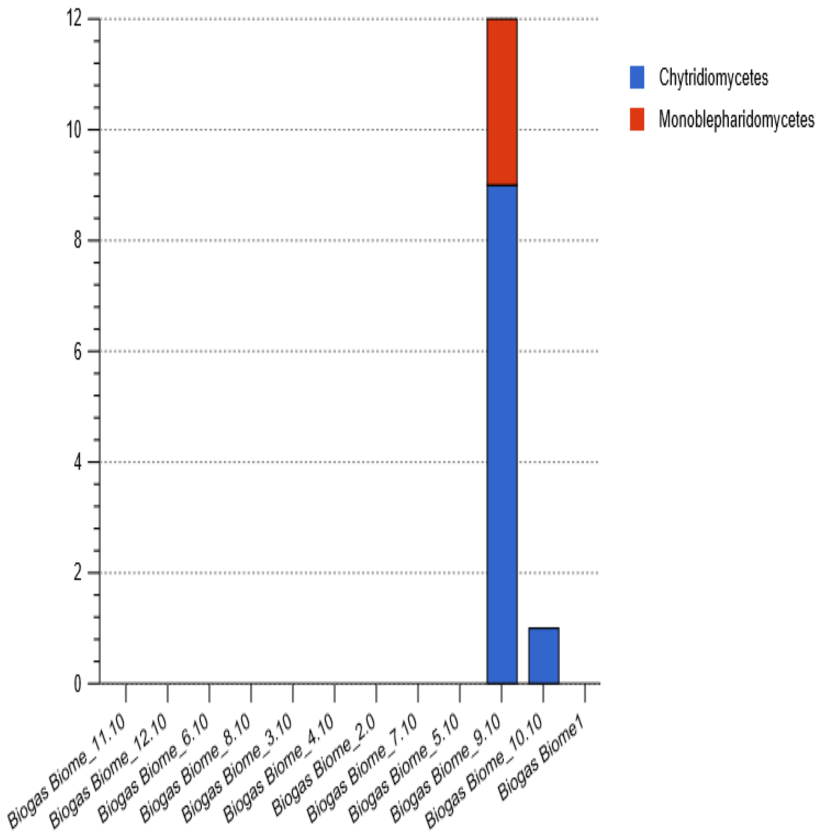


**SFig. 49:** Stacked barchat showing the two *Chytridiomycota* classes and their relative abundances in the two reactors.
