## Supplementary Fig. 50 for "Metagenomics survey unravels diversities of biogas’ microbiomes with potential to enhance its’ productivity in Kenya"

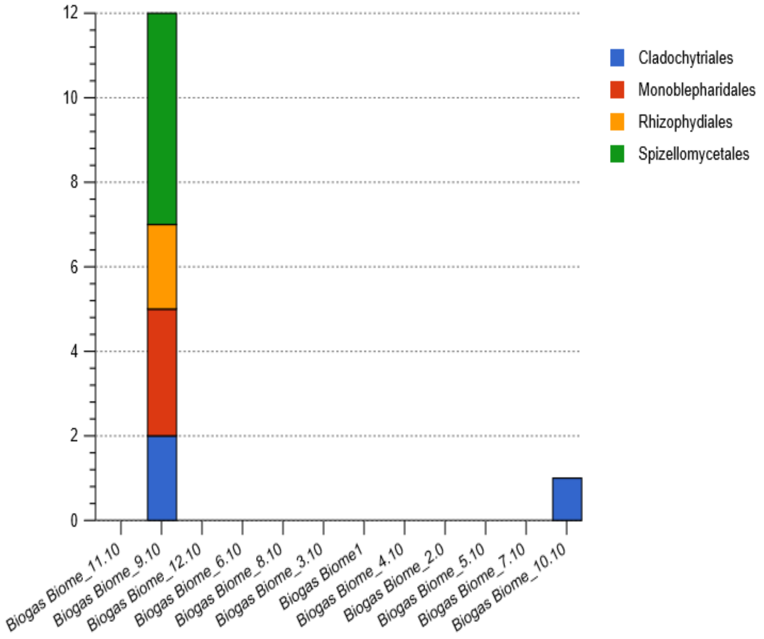


**SFig. 50:** Stacked barchat showing the four Chytridiomycota orders, and their relative abundances in the two reactors.
