## Supplementary Fig. 51 for "Metagenomics survey unravels diversities of biogas’ microbiomes with potential to enhance its’ productivity in Kenya"

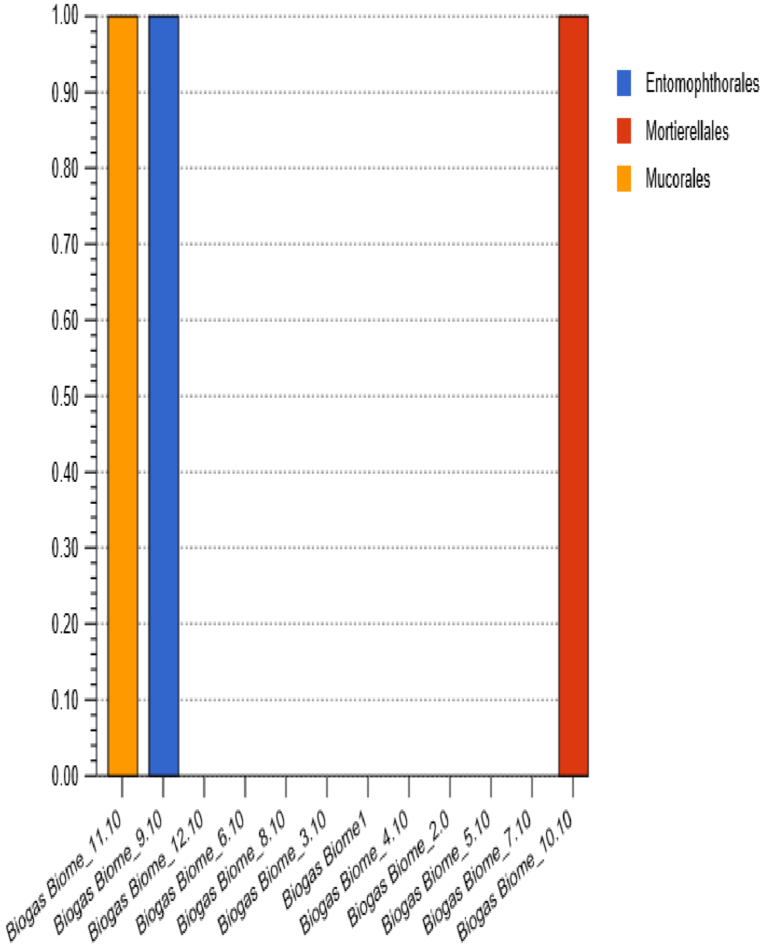


**SFig. 51:** Stacked barchat showing the three orders affiliated to unclassified fungal nucleotide reads. The orders are considered rare due to the fact that they were detected in only three reactors.
