## Supplementary Fig. 52 for "Metagenomics survey unravels diversities of biogas’ microbiomes with potential to enhance its’ productivity in Kenya"

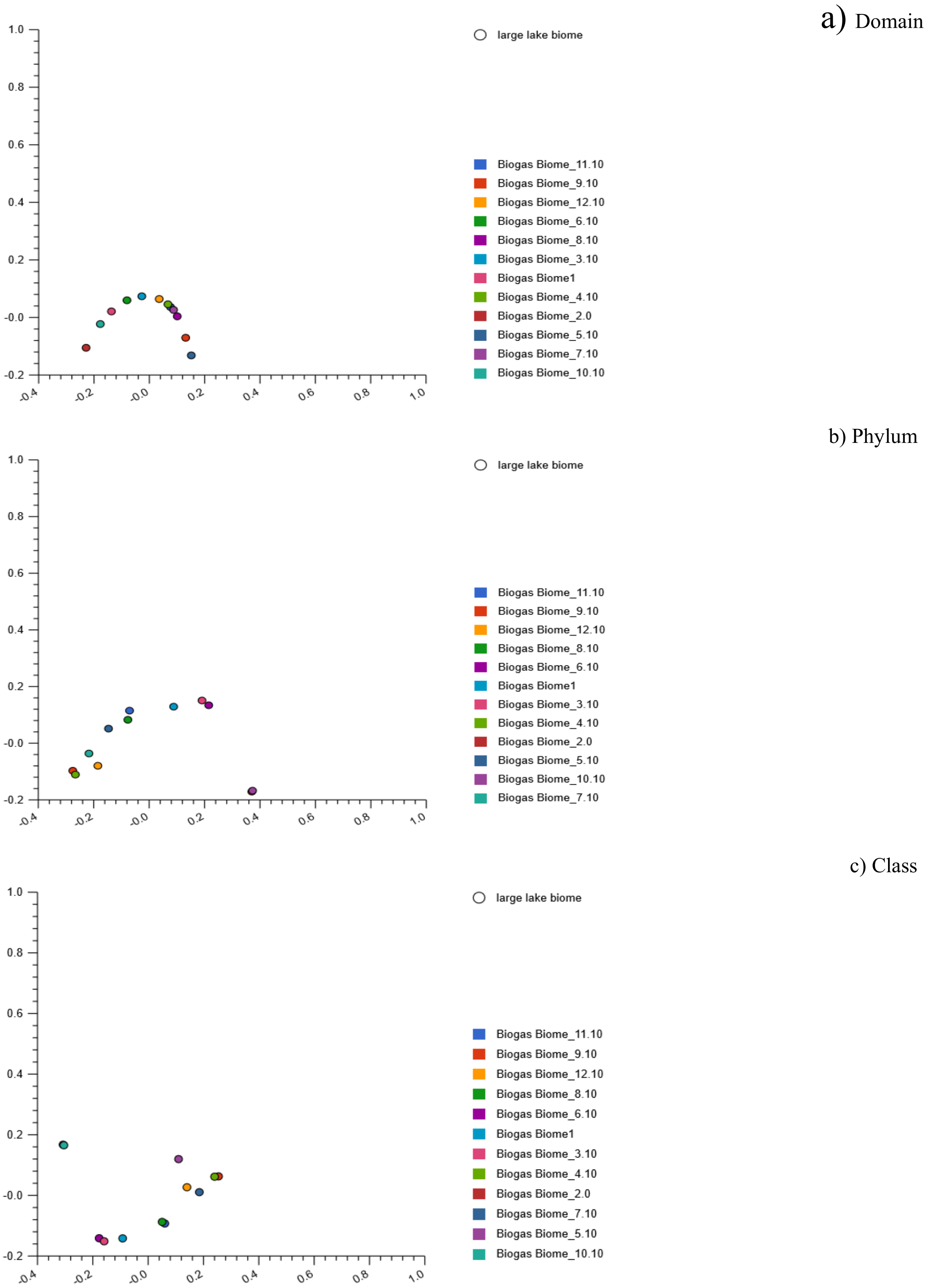


SFig. 52: The PCoA analysis revealing β-diversity of the twelve reactors at the three taxa level: a) Domain, b) Phylum, and c) class level. At the class level, the reads in reactor 2 and 10 were partially similar, located in the lower left quadrant, those in reactor 8 and 11 located in the upper right quadrant, reactor 4 and 9 in the lower right quadrant, while those in reactor 3 and 6 were located in the lower left quadrant of the plot.
