## Supplementary Table 1 for "Metagenomics survey unravels diversities of biogas’ microbiomes with potential to enhance its’ productivity in Kenya"

| STable 1: The quality control statistics of the utilized scaffolds of the twelve treatments | | | | | | | | | | |
|---|---|---|---|---|---|---|---|---|---|---|
| | **Statistics of the filtered scaffolds for downstream analysis** | | | | | **Statistics of scaffolds GC content and ambiquity** | | | | |
| **Sample id** | **Base pair** | **Sequence** | **Length (bp)** | **Av. Length Bp** | **Av. Std** | **GC%** | **Std dev.** | **GC-ratio** | **Std dev.** | **Amb. Reads** |
| S_1 | 89,283332 | 313,836 | 56-136,866 | 284 | 348.635 | 52.385 | 11.093 | 1.009 | 0.492 | 2,600 |
| S_2 | 107,701,975 | 394,463 | 56-64,851 | 273 | 189.197 | 55.033 | 11.236 | 0.908 | 0.463 | 1,5580 |
| S_3 | 90,908,777 | 310,336 | 56-55621 | 292 | 276.697 | 52.446 | 10.810 | 1.00 | 0.471 | 2,800 |
| S_4 | 83,626,041 | 275,624 | 55-34,847 | 303 | 372.306 | 49.967 | 10.900 | 1.108 | 0.514 | 1,220 |
| S_5 | 74,806,357 | 280,930 | 55-5,606 | 266 | 85.933 | 49.220 | 11.339 | 1.154 | 0.560 | 470 |
| S_6 | 75,808,027 | 253,949 | 56-55,621 | 298 | 355.114 | 53.626 | 10.547 | 0.948 | 0.441 | 10,026 |
| S_7 | 133,718,872 | 428,821 | 56-95,185 | 311 | 510.825 | 49.603 | 11.172 | 1.133 | 0.549 | 5,020 |
| S_8 | 63,285,799 | 212,099 | 56-56,718 | 298 | 292.399 | 50.066 | 10.639 | 1.100 | 0.507 | 1,620 |
| S_9 | 188,414,920 | 621,662 | 56-44,371 | 303 | 305.181 | 49.315 | 11.075 | 1.145 | 0.548 | 7,284 |
| S_10 | 56,387,075 | 217,027 | 56-64,831 | 259 | 171.992 | 55.568 | 11.027 | 0.884 | 0.439 | 340 |
| S_11 | 59,129,493 | 198,327 | 56-216,425 | 298 | 996.110 | 48.738 | 10.866 | 1.167 | 0.546 | 2,491 |
| S_12 | 96,394,918 | 312,853 | 56-238,965 | 308 | 1024.420 | 50.749 | 10.631 | 1.065 | 0.468 | 3,865 |

*Legend: S_*1: *reactor 1 or Biogas Biome* 1.10; *S_2*: *reactor 2.0 or Biogas Biome* 2.0; *S_3*: *reactor 3 or Biogas Biome* 3.10; *S_4*: *reactor 4 or Biogas Biome* 4.10; *S_5*: *reactor 5 or Biogas Biome* 5.10; *S_6*: *reactor 6 or Biogas Biome* 6.10; *S_7*: *reactor 7 or Biogas Biome* 7.10; *S_8*: *reactor 8 or Biogas Biome* 8.10; *S_9*: *reactor 9 or Biogas Biome* 9.10; *S_10*: *reactor 10 or Biogas Biome* 10.10; *S_11*: *reactor 11.10 or Biogas Biome* 11.10. *S*_12: *reactor 12 or Biogas Biome* 12.10. NB: Reactors are treatments
